# Single-Cell Roadmap of Early Hemato-Endothelial Development: Functions of Atf3, Zfp711 and Bcl6b

**DOI:** 10.1101/2025.02.23.639715

**Authors:** Ridvan Cetin, Giulia Picco, Jente van Staalduinen, Eric Bindels, Remco Hoogenboezem, Gregory van Beek, Mathijs A Sanders, Yaren Fidan, Ahmet Korkmaz, Joost Gribnau, Jeffrey van Haren, Danny Huylebroeck, Eskeatnaf Mulugeta, Frank Grosveld

## Abstract

Hematopoiesis is the process of producing blood cells. In mammalian embryos, hematopoiesis occurs in three consecutive overlapping waves (Neo et al. 2021; Dzierzak and Bigas 2018) that are regulated by transcription factors (TFs) and signaling proteins. We investigated the functions of three relatively poorly studied TFs in early embryonic hematopoietic development at single-cell resolution: Activating transcription factor 3 (Atf3), Zinc finger protein 711 (Zfp711), and B cell CLL/lymphoma 6, member B (Bcl6b). These TFs are upregulated early in development when hematopoietic and endothelial lineages separate from cardiac and other mesodermal lineages. We combined multiplexed single-cell RNA sequencing (scRNA-seq) and cell identity analysis using Flow Cytometric Analysis (FCA) with TF knockouts (KO) in *in-vitro* differentiating mouse embryonic stem cells (mESCs) to dissect the function of these TFs in lineage induction, specification, and separation. The *Atf3*-KO showed an increase of distinct mesodermal subpopulations, but a decrease of endothelial and erythro-myeloid progenitors (EMPs) by downregulation of important genes (i.e. *Runx1, Mafb, Egr1, Jun, Jund, Fos, Batf3*, and *Zf608)* that likely explains the effects on EMPs and the increased expression of interferon-related genes. In *Zfp711*-KO cells, the number of blood progenitor cells and erythroid cells increased, while the number of endothelial cells decreased. Furthermore, *Hoxa* expressing mesoderm decreased, while *Hoxb* expressing mesoderm increased. In contrast, the *Bcl6b*-KO had no observable effects on early hematopoiesis. In conclusion, we report the function of these three TFs function at different stages of hemato-endothelial lineage specification.

**HIGHLIGHTS:**

- Combined scRNA-seq and FCA on KO mESCs *in vitro* differentiation enabled unbiased identification of the respective TF functions during specific stages of hematoendothelial lineage specification.
- <u>*Atf3*-KO:</u> Increased abundance of mesodermal lineages and decreased endothelial lineages and EMPs.
  – Downregulation of key TF-encoding genes (e.g., *Runx1, Mafb, Egr1, Jun, Jund, Fos, Batf3, Zf608*) explain the effects on EMPs.
  – Generation of a distinct mesodermal subpopulation with high expression of sets of interferon and antiviral-related genes.
  – Upregulation of interferon and anti-viral-related genes, indicating Atf3 acts as a negative regulator of these genes.
- <u>*Zfp711*-KO:</u> Increased numbers of blood progenitor and erythroid cells. Decreased numbers of endothelial cells.
  – Shift in mesodermal populations: *Hoxa* expressing mesoderm decreased, and *Hoxb* expressing mesoderm increased.
  – *Hoxa9, Hoxa10, Hoxa13, Jun, Jund* and *Atf3* amongst the downregulated genes. The *Atf3* promoter has a potential Zfp711-binding site.
  – Increased expression in a given Endothelium subtype in endocardium *in vivo* and *in vitro*.
- <u>*Bcl6b*-KO:</u> No observable impact on early hematopoiesis.

## INTRODUCTION

After oocyte fertilization, the zygote of mammals slowly generates multi-cellularity before implantation of the blastocyst in the uterine wall. From that moment, gastrulation is accompanied by increased cell proliferation leading to drastic re-allocation of embryonic cells to the three embryonic germ layers and the formation of a primitive body plan. The need for nutrients and oxygen is initially met by diffusion via extra-embryonic tissues, but increasing numbers of cells and morphogenetic changes of the growing embryo rapidly exceeds the capabilities of diffusion. Therefore, vessel formation and blood cell formation are required to sustain embryonic development. The close functional relationship between the formation of blood vessels (which include endothelial cells) and hematopoiesis is reflected by shared developmental trajectories during the three embryonic hematopoietic waves. Hematopoietic Stem Cells (HSCs) are absent in early development and appear only after embryonic day (E)10.5 in the mouse embryo (Dzierzak and Bigas 2018). Thus, temporary solutions evolved during early development (Yamane 2020). The first wave of hematopoiesis and endothelial cell development (E7.5) starts with the differentiation of a group of mesodermal cells called Hemato-Endothelial Progenitors (HEPs), which differentiate into Blood Progenitors (BPs) and Endothelial cells. BPs generate nucleated primitive erythrocytes, primitive megakaryocytes and macrophages in blood islands that form in and from the inner mesodermal leaflet of the visceral yolk sac, not of the amnion. In the second wave, endothelial-like cells called Hemogenic Endothelium undergo an Endothelial-to-Hematopoietic Transition (EHT) to generate Erythro-Myeloid Progenitors (EMPs). These cells detach from the vascular bed and enter the circulation as precursor cells. The third wave occurs at E10.5 in the Hemogenic Endothelium of the dorsal aorta and vitelline/umbilical arteries, where HSCs are formed that will supply the blood cells throughout adult life (Tober et al. 2006; Neo et al. 2021; Harland et al. 2021).

Understanding the regulation of these embryonic waves of hematopoiesis is essential to generate hematopoietic progenitors/cells using protocols for instructed differentiation in cell culture. This approach could limit the dependency on blood donors and ultimately reduce the risks of complications related to bone marrow transplantation. The developmental regulation of the hemato-endothelial lineage is controlled by DNA-binding Transcription Factors (TFs) (Latchman 1997) (i.e., GATA1 binding to [A/T]GATA[A/G] to regulate, e.g., hemoglobin production (Evans et al. 1988)). Therefore, investigating the roles of such individual TFs, their gene regulatory network, and their cooperation with other TFs is crucial to understanding the regulatory mechanisms behind hemato-endothelial development. Technical improvements in single-cell analytic technologies, instructed stem cell differentiation mimicking embryonic development in cell culture, and the publicly available transcriptomics and proteomics (including TF interactomes) datasets/databases, lists of TF/TF-partners and their direct target genes, and surveys of epigenetic regulations, make it possible to investigate the roles of the TFs in developmental roadmaps at single-cell resolution (Karlsson et al. 2021; Uhlén et al. 2015; Kronman et al. 2024; Haniffa et al. 2021; Consortium 2004; Pijuan-Sala et al. 2019; Imaz-Rosshandler et al. 2023).

Using murine ESC differentiation *in vitro* we previously observed that several TFs are upregulated in embryoid bodies (EBs) when the cells of the future hematoendothelial lineage split from the cardiac mesodermal cells at day 4 of differentiation. Here the results of three relatively poorly studied TFs that are upregulated at this split in early embryonic hematopoietic development, are shown at single-cell resolution: Activating transcription factor 3 (*Atf3*), Zinc finger protein 711 (Zfp711) (the ortholog of human*ZNF711*), and B cell CLL/lymphoma 6, member B (Bcl6b). We answer three questions: **1**) are these three TFs important for early lineage choices, i.e. the hemato-endothelial versus late mesodermal lineages? **2**) are they important for later stages of differentiation, i.e., EHT, primitive hematopoiesis, and possibly other mesodermal lineages? **3**) which genes and pathways are changed upon their respective perturbation providing insight into their functional role(s)?

ATF3 is a member of the ATF/cAMP-responsive TF family (Hummler et al. 1994). It binds to cAMP response elements and contains a leucine zipper (bZIP) domain. ATF3 interacts with the bZIP domain of other proteins, including other ATF/CREB family members. While homodimers of ATF3 act as transcriptional repressors, ATF3 hetero-dimers (i.e., with ATF2, JUN, JUNB, and JUND) act as activators (Hai et al. 1999; Liang et al. 1996; Chen et al. 1994). ATF3 plays various roles in multiple cellular processes, including metabolism, in the immune system, during development, and in oncogenesis (Ku and Cheng 2020).

ZFP711 contains multiple C2H2-type zinc-finger domains and an acidic transactivation domain (Mardon et al. 1990), *Zfp711/ZNF711* has a similar binding motif as its paralogs ZFX and ZFY (and mouse *Zfx* and *Zfy*) (Ni et al. 2020). It binds to CpG-islands and may play a role in regulating promoter architecture (Ni et al. 2020; Rhie et al. 2018). Its acidic transactivation domain can recruit the histone lysine demethylases, resulting in the local removal of H3K4me1/me2 and H3K9me2 methylation marks, thereby enhancing the transcriptional activation of target genes (Wu et al. 2021;Kooy 2022).

BCL6B is a paralog of the TF BCL6 that acts as a transcriptional repressor and binds to the same DNA motif as BCL6. It has five Krüppel-type zinc fingers in the C-terminal end to bind DNA. BCL6B needs to dimerize with BCL6 for repressor activity, but their mRNA expression profiles across tissues differ, suggesting they may have other and/or additional specific roles (Hartatik et al. 2001; Takenaga et al. 2003; Takamori et al. 2004; Chattopadhyay et al. 2006; Ohnuki et al. 2012).

Using a workflow (see Materials and Methods) to analyze the three genes in one experiment, we show that Atf3 plays a role in EMP abundance/maturation, Endothelium abundance, and the generation of various new cell types in late mesodermal lineages, some of which show strong expression of interferon pathways-related genes; Zfp711 effects are mainly observed in mesodermal cell types abundance and on *Atf3* expression, one of the genes affected by the inactivation of *Zfp711*; Bcl6b shows no observable impact on either hematoendothelial lineage or mesodermal development. These findings highlight the strength of the adapted workflow to dissect TF functions in hematoendothelial lineage specification.

## RESULTS

### Experimental design and workflow with adapted multiplexed scRNA-seq and Flow Cytometric Analysis of Multiple TF knockouts during early hematopoietic development

We adapted available multiplexing technology for singlecell RNA sequencing (scRNA-seq), with multiplexing knockout (KO) and Control conditions (with biological replicates) and combined it with Flow Cytometric Analysis (FCA), to study the effects of the inactivation of each of three TFs in early embryonic hematopoietic development. Doing so, the depth and coverage of this study can be placed between large-scale multiplexed CRISPR-based perturbation studies covering multiple candidate genes together (Datlinger et al. 2017; Jaitin et al. 2016; Dixit et al. 2016; Adamson et al. 2016) and single-gene perturbation studies, the latter usually involving in-depth functional studies using multiple experimental procedures (Harland et al. 2021). With our adaption, we studied the role of three TF-encoding genes in one experiment (including three biological replicates), and capture sufficient numbers of cells to detect changes in low-abundance cell types, minimize potential batch effects (e.g., each replicate generating a separate library), and prevent the loss of biological information by skipping specific computational integration steps. Following the scRNA-seq analysis, its results were compared with publicly available datasets, enabling us to categorize our findings. This categorized information will markedly facilitate future indepth follow-up studies of the studied TFs in hematoendothelial development.

Based on observations of increased expression of *Atf3, Bcl6b*, and *Zfp711*, both *in vitro* and *in vivo* and that their roles remain poorly understood in hematoendothelial development, we investigated their role in this complex process by genetically inactivating each of their genes using the same mouse embryonic stem cells (mESCs) differentiation in which we observed increased expression of the three TFs.

The separate KO mESC lines were generated using CRISPR-Cas9, resulting in *Atf3*-KO, *Bcl6b*-KO, and *Zfp711*-KO mESC cells, and their differentiation was compared to their parent non-targeted Control (NTControl). FCA was used to probe multiple biological replicates (4x for the TF-KO clones, 6x for the NTControl clones) of their *in vitro* hematopoietic differentiation. For FCA, the differentiation process was repeated three times independently, thereby minimizing intra- and inter-assay variation in differentiation efficiencies. Two time points (days 4 (D4, non-pulsating EBs) and 7 (D7, pulsating EBs)) were used to score the efficiency of differentiation. This was done by using CD309 (encoded by *Kdr/Flk1*) and CD140a (*Pdgfra)* markers (Harland et al. 2021), which determined the percentage of HEPs (Flk1^+^/Pdgfra^-^) on D4 and mesodermal lineages (Flk1^+^/Pdgfra^+^). The next assessment used a set of 7 markers (CD41 (encoded by *Itga2b*), CD71 (*Tfrc*), CD144 (*Cdh5*), CD102 (*Icam2*), CD45 (*Ptprc*), CD140a (*Pdgfra*), and CD140b (*Pdgfrb*); these markers were combined from the literature and compared with the expression of markers *in vitro* (Harland et al. 2021; Sievert et al. 2014; Psaila et al. 2016) (Pijuan-Sala et al. 2019)), that identify the cells involved in the first wave and early-phase second wave of hematopoiesis (Fig. 1A,C; for details, see also Supplemental Fig. S1A-E). Next, scRNA-seq was carried out at D7 as three biological replicates (triplicates), using cell multiplexing oligonucleotides (CMO, Fig. 1B). This captured transcriptomic changes at the most appropriate time point, considering the asynchronous nature of the ESC differentiation after D7 (*data not shown*).

**Figure 1.**
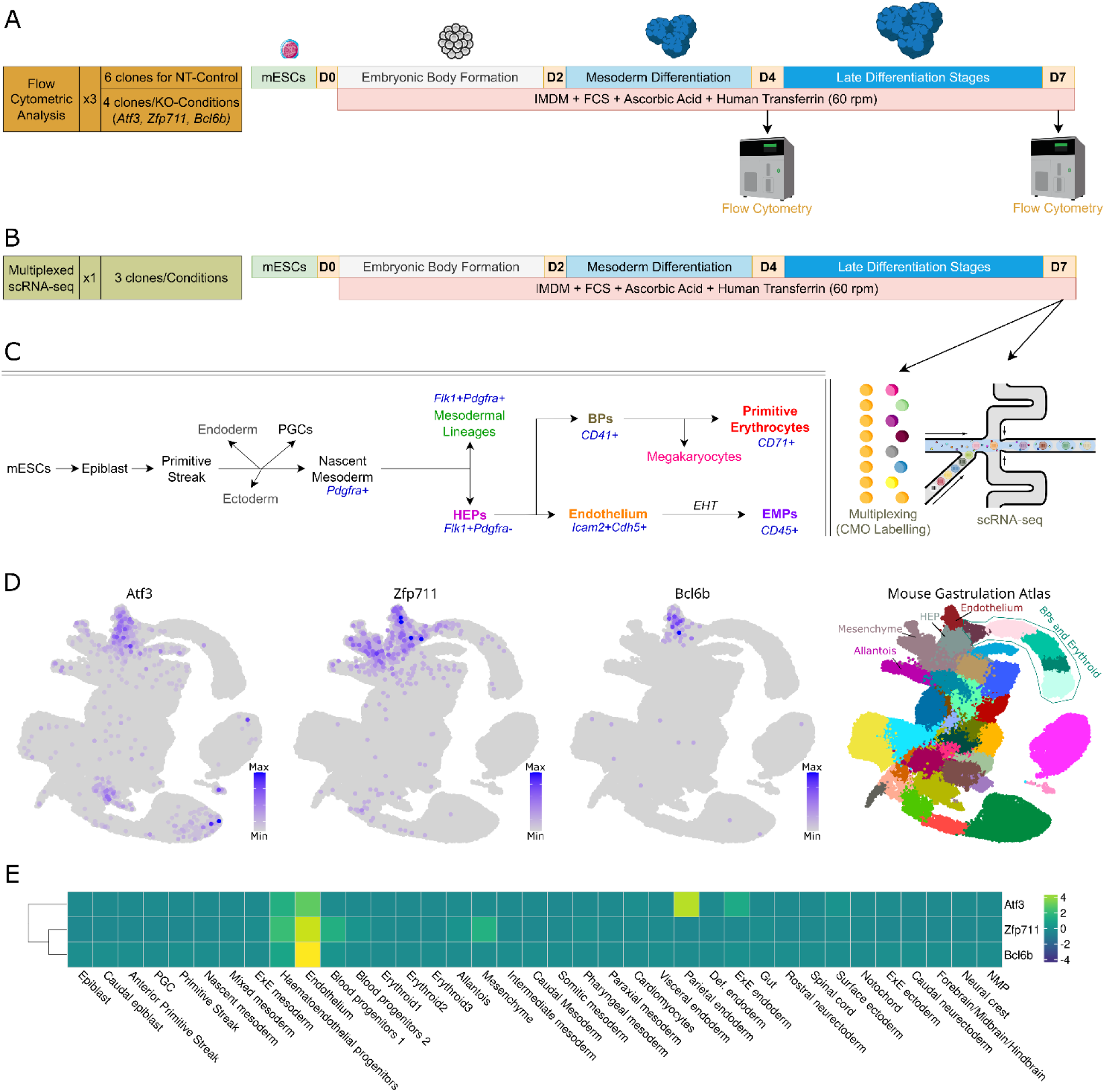
Experimental workflow, differentiation stages, and TF gene expression *in vivo*. **Panel A** summarizes the experimental workflow and time points of the Flow Cytometric Analysis. **Panel B** summarizes the experimental workflow and time point of multiplexed single-cell RNA sequencing. **Panel C** shows the developmental trajectory of the mesodermal and early hematopoietic lineages, including the markers used for the Flow Cytometric Analysis. **Panel D-E** shows expression of *Atf3, Zfp711*, and *Bcl6b*, and cell type annotation plots/heatmaps from an available atlas of mouse gastrulation.

This also minimized batch effects in scRNA-seq processing, made the biological replicates cost-efficient, and minimized clonal bias. Five scRNA-seq libraries/lanes were generated to capture sufficient cells, notably including small subpopulations of cells.

### Atf3, *Zfp711*, and *Bcl6b* exhibit increased expression in HEP and Endothelial cells both *in vivo* and *in vitro*

In a separate set of scRNA-seq experiments, it was clear that the expression of *Atf3, Bcl6b*, and *Zfp711* increased in HEP and endothelial cells when compared to other mesodermal lineages during mESC at D4 of *in vitro* differentiation, i.e. when the hematoendothelial lineage splits from other mesodermal lineages. This is also seen at D7 when further differentiation into cells that can be annotated to mesodermal, erythroid, and endothelial lineages occurs. The scRNA-seq results on mouse embryos (including E6.5-8.5 embryos) showed similar expression profiles for our three TFs of interest (Pijuan-Sala et al. 2019). *Bcl6b* is expressed almost exclusively in mESC-derived HEPs, whereas *Atf3* and *Zfp711* are each expressed in other mESCs-derived mesodermal lineages (Fig. 1D,E; see also Supplemental Fig. S2A-D).

### Multiplexed scRNA-seq captures cell types from naïve mESCs to mature hemato-endothelial lineages, without observable batch effects

After demultiplexing the scRNA-seq data, (Supplemental Fig. S3A), and performing other quality controls (removal of low-quality cells, ambient RNA, cell doublets, and cell cycle effects, a total of 71,107 cells were captured in 5 libraries and divided among 12 samples. These 12 samples represent 3 x biological replicates per condition (about 6,000 cells/clone, with a minimum of 4,954 and maximum of 7,457 cells) and four conditions (*Atf3*-KO, *Zfp711*-KO, *Bcl6b*-KO, and NT-Control) (16,404-18,948 cells per condition) (see Supplemental Fig. S3B). No significant differences in the resulting scRNA-seq-based cell distributions were observed between each of the libraries and respective conditions (Supplemental Fig. S3C-E). Therefore, no computational batch correction and integration methods were applied. The scRNA-seq data show a continuous differentiation pattern starting from naive pluripotent mESCs (*Zfp42*^+^, *Esrrb*^+^) to multiple differentiated cellular states (e.g., Erythroid (*Hba-x*^+^), EMP (*Ptprc*/CD45^+^), Endothelium (*Cdh5*/CD144^+^), Megakaryocytes (*Vwf* ^+^), extra-embryonic tissues (i.e., ExtraEmbryonic Ectoderm and Extra-Embryonic Endoderm) and late mesodermal lineages (for each selected marker gene, see Fig. 2A; see also Supplemental Fig. S4AC). The cell subpopulations were annotated at three levels for analysis, i.e., “groups,” “clusters,” and “sub-clusters” (Fig. 2B,C). There are three groups: A first group of cells related to early differentiation stages of the starting naive ESCs (Early Differentiation), a second group of cells with hematoendothelial markers (Hematopoietic) and a third group of cells at later stages of their mesodermal development (Late Mesoderm) (Fig. 2B; see also Fig. 3). Likewise, 15 clusters (Fig. 2B) and 40 subclusters (Fig. 2C) could be discerned by virtue of highly expressed, acknowledged marker genes (as listed in Fig. 3, columns 1-5).

**Figure 2.**
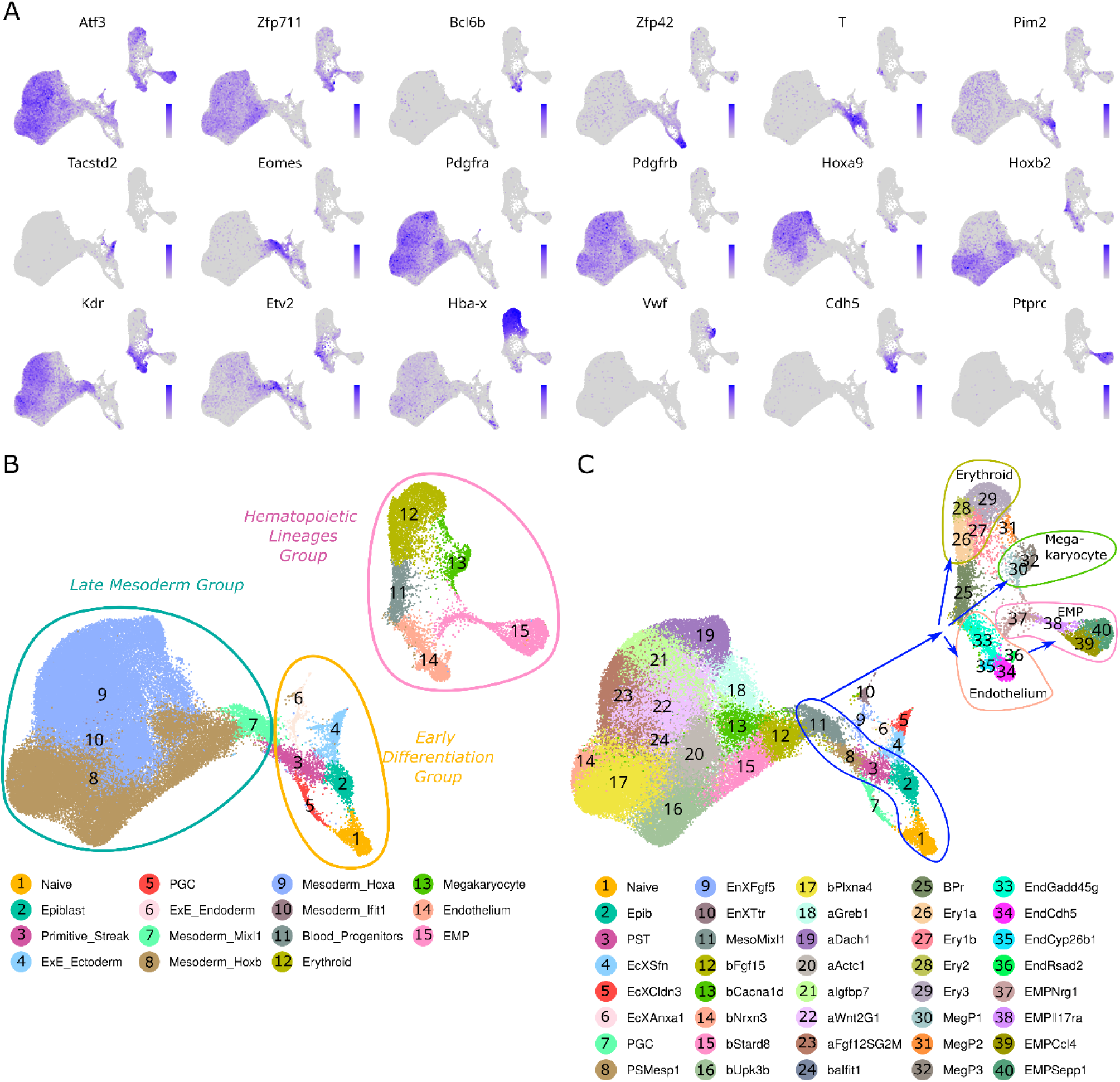
Identification of cell groups and clusters and sub-clusters therein, based on marker gene expression in scRNA-seq results of D7 embryoid bodies. **Panel A** shows the expression of each of the three TFs studied here, plus known individual marker genes, on UMAPs. **Panel B** shows the annotation of the cells at group and cluster levels. The three groups are bordered with lines (orange, turquoise, and pink), and the 15 discernable clusters are colored by their cluster annotation and numbered according to the color codes below each panel. **Panel C** similarly shows the 40 discernable sub-clusters. Blue bordered lines and arrows highlight the predicted differentiation trajectory starting from “Naïve,” following the mesoderm emergence (2,3,8,11) and ending in Erythroid (26,27,28,29), Megakaryocyte (30,32), Endothelium (33, 34,35,36) and transitioned EMP cell (37,38,39,40).

**Figure 3.**
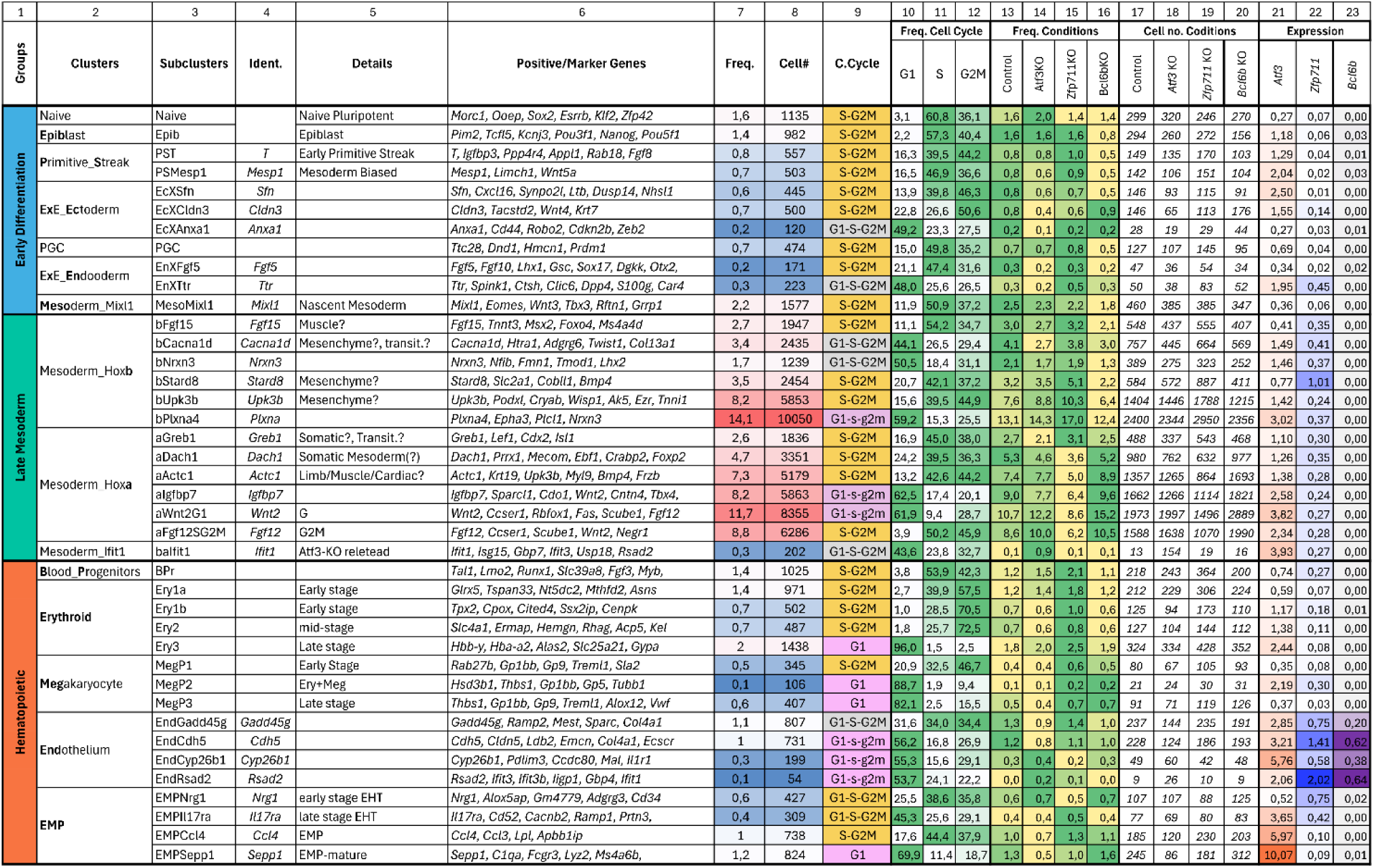
Details of cell lineage, such as group, cluster, and sub-cluster, the applied nomenclature, and characteristics of the cell populations. **Columns 1-3** show the annotation levels; **4** shows the genes used for identification/annotation of the sub-clusters; **5** shows the additional details; **6** shows the marker genes for each sub-cluster; **7** shows the frequency of the sub-clusters; **8** shows the number of cells; **9** shows dominant cell cycle stage(s); **10-12** show the frequency of the cell cycle stages (low [white] high [green]); **13-16** show the frequency of each condition in each sub-cluster (low [yellow] high [green]); and **17-20** show the number of the cells from each condition. **21-23** show average expression of TFs *Atf3, Zfp711, and Bcl6b* in each sub-cluster, expression level are colored highlighted with low (light) high (dark) tones of orange, blue, and purple.

The annotation of the 40 sub-clusters was done using gene expression, division rate and quiescence states, and transition and maturation stages (for a compilation, see Fig. 3, columns 6, 9-12). To name them (listed in Fig. 3, column 3), one of the following two options was used on top of the incorporated abbreviation of the cluster itself (Fig. 3; also listed in Fig. 2C): either sub-clusters were given an extra number corresponding to differentiation maturity (e.g., Ery3 for Erythroid 3, being the late stage) or the sub-cluster name incorporated one of the highly-expressed genes in those sub-clusters, such as aGreb1 (of the *Hoxa*^+^ Mesoderm, with high *Greb1* expression) and EMPSepp1 (EMP hematopoietic cells, with high *Sepp1*). Details of the marker genes for sub-clusters, their pseudo-time trajectory, the documented cell cycle distribution, and UMAPs for each group, are shown in Supplemental Fig. S4 to S7. Sub-cluster annotations are particularly useful for downstream analysis, and the data sets were divided into three groups (Early Differentiation, Hematopoietic, and Late Mesoderm) (Supplemental Fig. S5F-H).

### Deletion of ATF3 shows an increased expression that parallels the progression to EHT

Both alleles of *Atf3* were knocked out by using Crispr/Cas9 and two gRNAs, removing approximately 8 kb encompassing exons 2-3 and part of the terminal coding exon-4, resulting in the absence of detectable *Atf3* mRNA (Fig. 4A). scRNA-seq showed that the average expression of *Atf3* was highest in EMPs on D7, followed by Endothelium (Endothelium and EMPs constitute EHT) and Late Mesoderm. Within the EMP sub-clusters, the levels of *Atf3* increased with EMP maturation (EMPNrg1 = 0.52, EMPIl17ra = 3.65, EMPCcl4 = 5.97, EMPSepp1 = 10.07). The lowest *Atf3* expression was observed in naïve pluripotent cells and cells with primordial germ cell (PGCs) signature (Fig 3, column 21; see also Supplemental Fig. S4C, green box).

**Figure 4.**
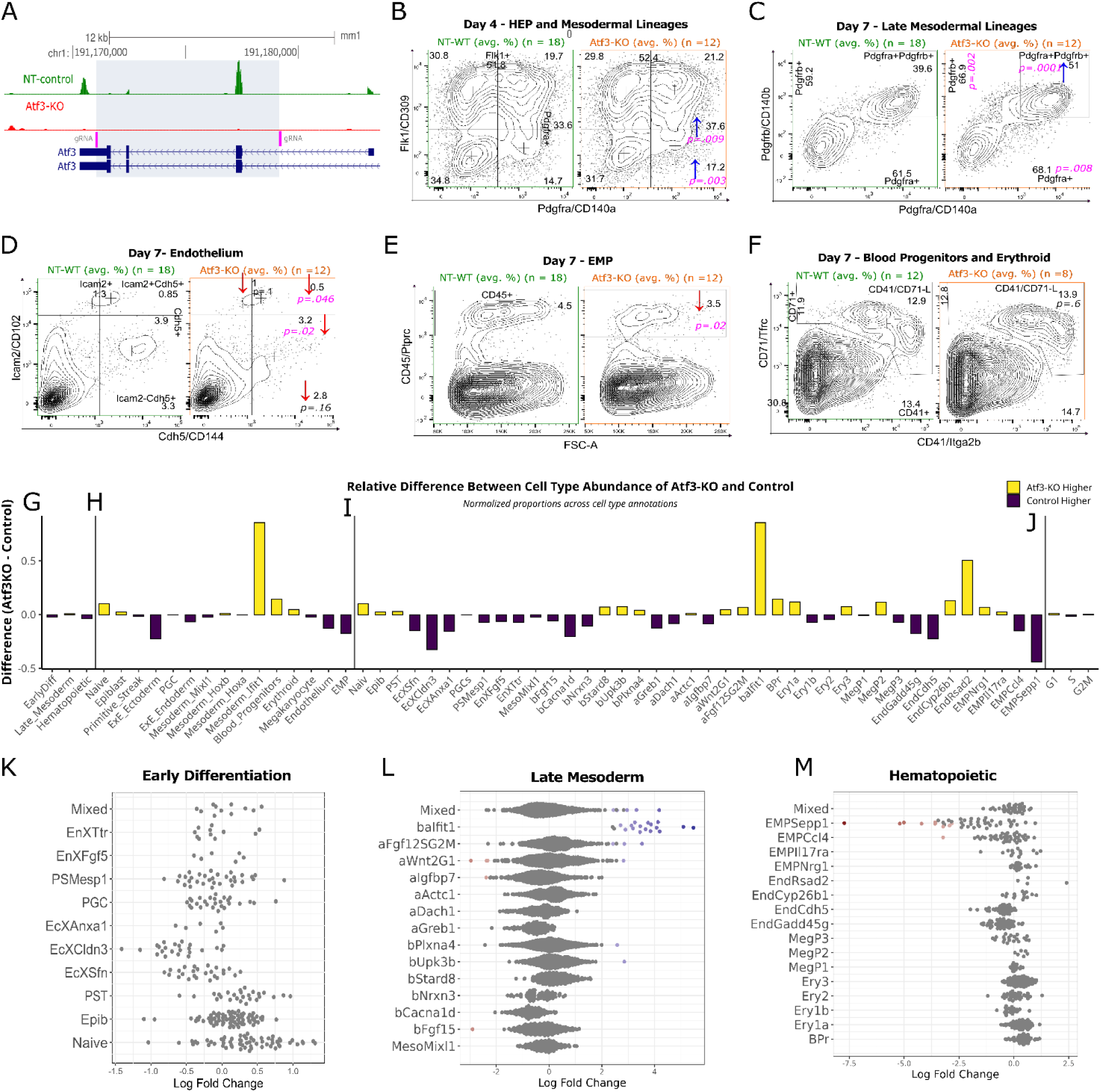
Results of the Flow Cytometric Analysis and differential abundance analysis of the *Atf3*-KO ESCs upon differentiation. **Panel A** shows the CRISPR-Cas approach for knocking out *Atf3*. The positions of the gRNAs are shown in pink. The peaks represent the scRNA-seq result for the control mESCs (in green) and the KO (red). The *Atf3* gene structure is shown in dark blue. **Panel B** shows examples of FACS plots from Atf3-KO and NT-Control after four days of differentiation, detecting the cell surface markers Pdgfra and Flk1. The numbers in the four corners show the average percentage of the respective quadrant. The numbers on the middle-right and top-right show the mean percentages of Pdgfra^+^ or Flk1^+^ cells. The arrows indicate an increase (in blue) or decrease (red) in the percentage of cells in a quadrant, together with its significance. Panels **C-F** show similar plots for day 7 Late Mesodermal lineages, Endothelium, EMP, and Blood Progenitors - Erythroid cells. **Panels G-I** compare the number of NT-Control cells versus the *Atf3*-KO based on groups, clusters, and sub-clusters. Panel **J** compares the number of cells in the respective phases of the cell cycle. **Panels K-M** show the fold-change in the differential abundance analysis in the different groups upon creating small neighborhoods (small groups of cells; shown as dots, with each dot representing in most cases 50-100 cells). Spatial FDR < 0.1 neighborhoods are shown in blue (*Atf3*-KO enriched) or red (*Atf3*-KO depleted); the rest is shown in grey. The text on the y-axis indicates sub-clusters, and the log fold change is shown on the x-axis. “Mixed” on the y-axis indicates a neighborhood that does not belong to any sub-cluster (using a 50% threshold).

### *FCA shows that Atf3* deficiency affects the abundance of the Mesodermal/Late-Mesodermal cells at D4 and D7

Surface marker staining and FCA of D4 cells showed that the ESC-derived Pdgfra^+^ cells (used as a general mesodermal marker) and Flk1^-^ Pdgfra^+^ cells (early mesoderm and/or endodermal lineages) had increased significantly upon *Atf3* inactivation, whereas Flk1^+^Pdgfra^-^ (HEPs), Flk1^+^Pdgfra^+^(mesodermal lineages), and Flk1^-^Pdgfra^-^ cells did not (Fig. 4B; see also Supplemental Fig. S8 and S9; Supplemental Table S1A,B). Thus, *Atf3* deficiency increased mesodermal differentiation: with 11.8% in the Pdgfra^+^ population (33.7 for NT-Control versus 37.6 for *Atf3*-KO cells (33.7/37.6)) and with 17% (14.7/17.2) in Flk1^**-**^Pdgfra^**+**^ cells. There was a decrease in Flk1^-^Pdgfra^-^ cells of 8.7% (34.8/31.7) at D4. Flk1^+^Pdgfra^-^ cells (representing HEPs) decreased slightly (2.9% (30.8/29.9)), whereas Flk1^**+**^Pdgfra^**+**^ cells (other Mesodermal Lineages, including cardiac, somatic, mesenchymal cells) increased by 7.3% (19.7/21.2), indicating a limited shift towards Flk1^**+**^Pdgfra^**+**^ cells.

The differentiation protocol strongly enriches mesoderm formation but is known to still produce small numbers of cells of extra-embryonic identity. For example, the number of extra-endodermal cells that also express *Pdgfra* might complicate the use of Pdgfra as a *bona fide* mesodermal marker in the scRNA-seq at D7. Hence, staining of Pdgfra alone cannot lead to firm conclusions. At D7, the staining of not only Pdgfra^**+**^ (10.6% (61.5/68.1)) but also Pdgfrb^**+**^ (13% (59.2/66.9)) and CD140a^**+**^CD140b^**+**^ (representing mesodermal lineages) (28.7% (39.6/51.0)) had increased significantly relative to NT-Control cells (Fig. 4C; see also Supplemental Fig. S9; Supplemental Table S1A,B). The combination of the D4 Flk1/Pdgfra and D7 Pdgfra/Pdgfrb staining thus indicated that Atf3 deficiency increased the mesodermal differentiation efficiency by up to 10%.

### FCA shows that Atf3 deficiency affects the abundance of the Endothelium and EMP, but not Erythroid Lineages at D7

On D7, cells stained for the endothelial markers Icam2 and Cdh5 decreased significantly (Cdh5^**+**^Icam2^+^: 41% (0.85/0.50)) compared to the NT-Control (Fig. 4D). Staining for CD45 on EMPs showed a significant decrease from 4.5% in the NT-Control to 3.5% in the *Atf3*KO-derived cells (a 21.6% change; Fig. 4E). Staining of Blood Progenitors and Erythroid cells for CD41 and CD71 showed a non-significant 7.8% change (NT-Control = 12.9%, *Atf3*-KO = 13.9%) (Fig. 4F, see also Supplemental Fig. S9; Supplemental Table S1A,B). Thus, the absence of Atf3 affects Endothelial and EMP lineages, but not the primitive wave of hematopoiesis (Blood Progenitors and Erythroid).

### scRNA-seq shows that Atf3 deficiency affects the abundance of Mesoderm-Ifit1, Endothelium, and EMP at D7

Changes in the composition of the various cell populations, detected by scRNA-seq, were further checked with two different Differential Abundance Analyses (DAA), i.e. annotated cell type proportion comparison (speckle/propeller Phipson et al. 2022) (Fig. 4G-J; see also Supplemental Fig. S10; Supplemental Table S2) and using k-nearest neighbor graphs (miloR - Dann et al. 2022) (Fig.4K-M; see also Supplemental Fig. S11; Supplemental Table S3), respectively.

DAA at group-level cell annotation showed no significant changes, but revealed a minor increase in Late Mesoderm and a slight decrease in Hematopoietic and Early Differentiation (Fig. 4G; Supplemental Fig. S10A). At the cluster level, DAA shows that a Late Mesodermal Mesoderm cluster (Ifit1) had formed. The Early Differentiation cluster ExE-Ectoderm shows a decrease in the *Atf3*-KO cells upon differentiation. Concerning Hematopoietic lineages, a minor increase was observed in BP and Erythroid cells, whereas Endothelium and EMP showed a decrease (Fig. 4H; Supplemental Fig. S10B).

In addition to the observations for the ExE-Ectoderm and Hematopoietic lineages, DAA showed a similar profile of cell type abundance of the mesoderm-Ifit1 (balfit1, at subcluster annotated level) as also seen in the Endothelium sub-cluster EndRsad2 (albeit not as strong as baIfit1), which was increased in the *Atf3*-KO cells. The two major Endothelium sub-clusters (EndGadd45g, EndCdh5) decreased in abundance. The decrease in the EMP cluster showed that three sub-clusters of the EMPs were affected (Fig. 4I; Supplemental Fig. S10C), whereas there was no change in the total cell cycle phase (Fig. 4J; Supplemental Fig. S10D). Immature EMPs (EMPNrg1) showed some increase, while the more mature EMPs were decreased (EMPSepp1, EMPCcl4) (Fig. 4I,M), suggesting that Endothelial-Hematopoietic Transition was affected (Supplemental Table S2A-H; Supplemental Fig. S10). A similar result was obtained by k-nearest neighbor graphs: there are no significant changes in Early differentiation, whereas Late Mesoderm (baIfit1) and Hematopoietic cells (EMPSepp1, EMPCcl4, EndRsad2) showed significant changes ((Fig. 4 K-M; Supplemental Fig. S11; Supplemental Table 3A-C).

When combined, FCA and DAA of the *Atf3*-KO versus NT-Control differentiated mESCs further showed that the results of both single-cell-based methods aligned and showed a decreased proportion of CD45^+^ cells and a decreased abundance of EMP clusters and sub-clusters. As mentioned, EMP was the highest *Atf3*-expressing cluster, and its sub-clusters show a change (increase) from immature EMPNrg1 (expression level 0.5), EMPIl17ra (3.60), EMPCcl4 (6) to mature EMPSepp1 (10.1), i.e. correlating with EMP maturation. These findings are in agreement with previous reports that used zebrafish (Yin et al. 2020).

### The absence of Atf3 shows upregulation of genes in Late Mesoderm and downregulation in EMPs

Pseudo-bulk Differential Gene Expression Analysis (DGEA) was performed to document changes in gene expression at the group, cluster, and sub-cluster levels to compare the *Atf3*-KO and NT-Control mESC differentiation. After removing the genes expressed at very low levels (which cannot be analyzed reliably with DGEA), the Differentially Expressed Genes (DEGs) were determined at a statistically significance level using the Benjamini-Hochberg method of p-value adjustment of < 0.05 in DESeq2 (Love et al. 2014).

The number of DEGs in the *Atf3*-KO condition is higher in EMP, Late Mesoderm, and their derivatives. EMP has the highest average *Atf3* expression and the highest number of DEGs (Fig. 5A, red circles). EMP and Late Mesoderm show different profiles of DEGs; most of the Late Mesoderm DEGs are upregulated (69 genes up versus 20 down), whereas EMP DEGs are downregulated (17 up versus 107 down) (Fig. 5A,B; Supplemental Fig. S12A,B; Supplemental Table S4A-H). Usually, larger populations of cells have a higher number of genes included in the DGEA, which affects the number of DEGS. In our case, this results in Late Mesoderm with 16,803 genes versus EMP with 9,631 genes. Even though the EMP cell number is lower, the increase in DEGs seen in EMP is therefore a biological. Also, the two subpopulations representing mature EMPs (EMPCcl4 and EMPSepp1) had the largest number (150 DEGs) when compared to the whole EMP cluster (124 DEGs) and the hematopoietic group (45 DEGs) (Fig. 5A, Table 1; Supplemental Table S4AH; Supplemental Table S9A).

**Figure 5.**
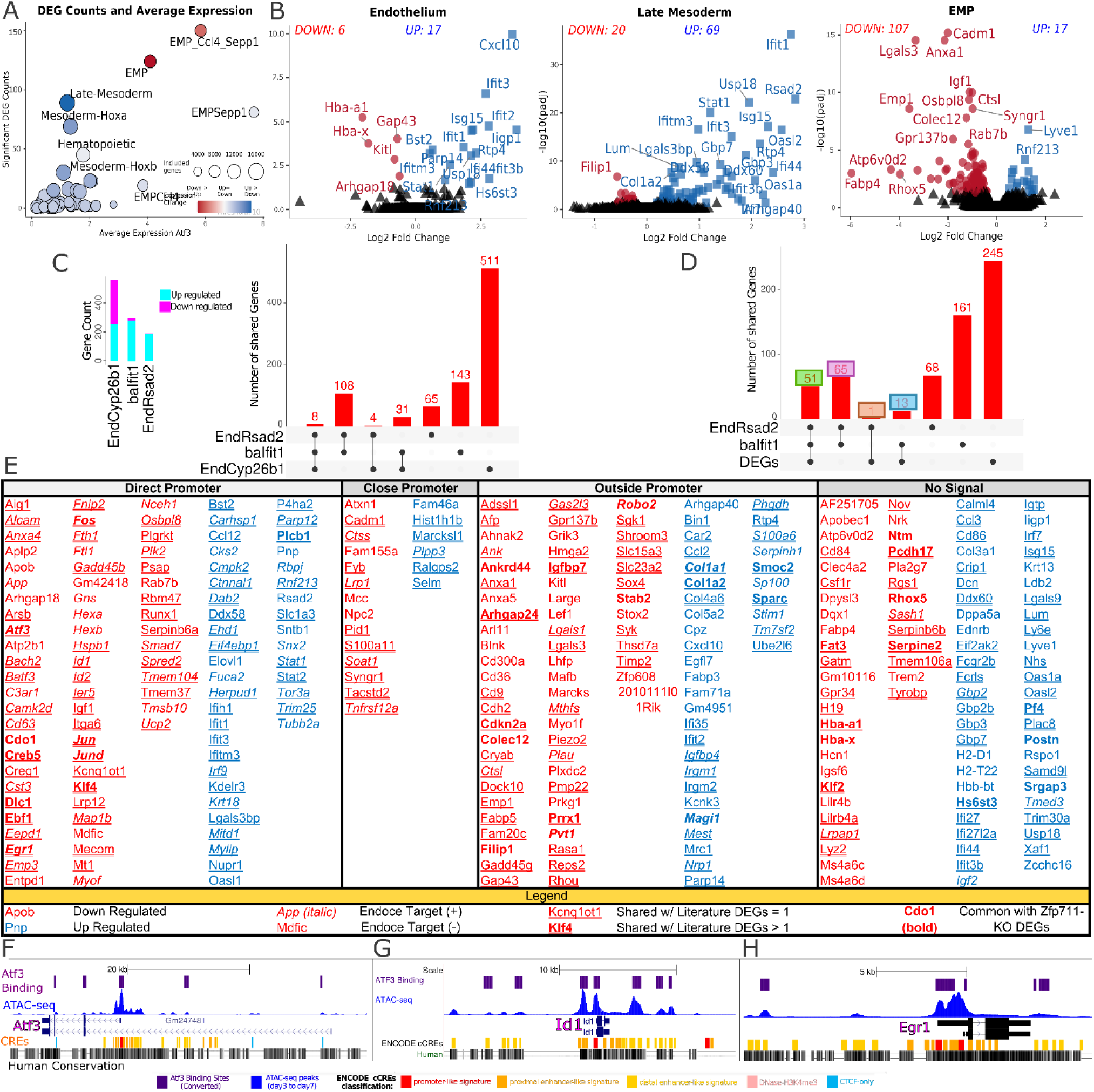
Results and categorization of the Differential Gene Expression Analysis (DGEA) of the *Atf3*-KO cells. **Panel A** shows the average expression level and number of significant DEGs (padj < 0.05) in groups, clusters, and sub-clusters. The size of the circles indicates the number of genes included in the analysis (after removing low-expressed genes) at group, cluster, or sub-cluster level. The color of the circle indicates up- (in blue) or downregulation (red). **Panel B** shows the up- or downregulation of genes in the Endothelium and EMP clusters and Late Mesoderm group as volcano plot. **Panel C** shows the number of up- or downregulated genes in three unique sub-clusters and the number of shared genes between the sub-clusters. **Panel D** shows the number of shared genes between the unique sub-cluster potentially caused by the *Atf3*-KO and the total DEGs. The colored boxes refer to the number of intersected genes shown in Supplemental Figure 12F. **Panel E** shows the total list of significant pseudobulk DEGs from scRNA-seq at day 7, which categorized/visualized with Atf3 binding sites, up-downregulation, intersected with ATF3 targets list from Harmonize - ENCODE Transcription Factor Targets (Rouillard et al. 2016), and shared with ATF3/Atf3 related literature DEGs. **Panel F** to **G** shows three examples of Atf3 binding sites with ATAC-seq peaks in three example DEGs.

### The *Atf3*-KO severely affects a small number of cells in the Late Mesoderm, resulting in the formation of a new cell type

The population of cells in a new cell type baIfit1 (Hoxb^**+**^ Hoxa^**+**^ Ifit1^**+**^) and the EndRsad2 sub-clusters were more prominent in the differentiated *Atf3*-KO cells (Fig. 3, columns 3 and 13-18; and rows sub-cluster balfit1 and EnsRsad2, Fig. 4I,L). Due to the small number of cells in these sub-clusters (and even fewer cells in the NT-Control), a pseudo-bulk DGEA was uninformative because most genes with low counts were filtered out. Therefore, a search for markers of these sub-clusters was done while comparing their neighbors in a complete single-cell dataset, which included all conditions with differential gene expression based on the non-parametric Wilcoxon rank sum test. The comparison of cluster baIfit1 to the other mesodermal clusters (*Hoxa*^**+**^ and *Hoxb*^**+**^) showed 267 upgenes (e.g., *Irf7, Stat1, Stat2, Irf9, Rsad2*, and *Ifit1*) versus 23 downregulated (Supplemental Fig. S12C; Supplemental Table S4I).

When the EndRsad2 sub-cluster is compared to the rest of the sub-clusters of the Endothelium, 182 genes were found upregulated (e.g., *Irf7, Stat1, Stat2, Trim30a, Sp100, Ifit1*, and *Rsad2*) versus 3 downregulated (Supplemental Fig. S12D; Supplemental Table S4J). A different result was obtained for EndCyp26b1 (253 genes upregulated (e.g., *Hand2, Sox6, Gata4, Id2 Zfp711, Mef2c, Tbx20, and Meis2*), 301 downregulated) (Supplemental Fig. S12E; Supplemental Table 4K). When comparing these three small sub-clusters, 8 genes were shared by all three, and EndRsad2 and balfit1 shared an extra 108 genes (Fig. 5C).

Thus, both the DAA and the expression of selected, significant marker genes indicate a significant overlap between these two sub-clusters that belong to two distinct groups (Late mesoderm and Hematopoietic lineages). The specific and, therefore, significant markers of the EndRsad2 and baIfit1 sub-clusters overlap with the list of all DEGs of the *Atf3*-KO versus NT-Control cells (Fig. 5C,D; Supplemental Fig. S12F). These 51 genes are primarily involved in interferon-alpha and gamma signaling and response to viruses (i.e. *Bst2, Ifit1, Ifit2, Ifit3, Ifitm3*), suggesting that Atf3 is important in this process and that the absence of Atf3 can lead to the emergence of baIfit1 and strongly affects the cell type abundance of EndRsad2.

### *Atf3*-KO DEGs categorization by their distance to Atf3 DNA binding peaks

In order to categorize and link each DEG to *Atf3*, the DEG list was intersected with different types data. First, single-nearest-gene-neighbor analysis was used with GREAT (McLean et al. 2010) within 1Mb of a combination of ChIP-seq data (Supplemental Table S8A), and this for the mouse and human genome (see Material Methods), ATAC-seq data (*our unpublished data*), and the presence of an Atf3-binding motif. DEGs with an Atf3 binding site close to or in the promoter/and transcription start site (TSS) (defined here as −300 bp to +125 bp of the TSS) are likely regulated directly by Atf3, and are called “Promoter” peaks. An up- or downstream distance of 2kb from the TSS is assigned “Close Promoter”. The other category binds at a further distance from the TSS and is called “Outside Promoter”. DEGs without a neighboring Atf3 binding site are called “No Signal” and likely represent the result of downstream effects of the perturbed genes directly regulated by Atf3. This analysis showed 223 genes to be potentially directly regulated by Atf3, and 87 as secondary genes or regulated by very distant Atf3 binding sites (Fig. 5E-H and Table 2; see also Supplemental Fig. S13; Supplemental Table S7A-E). The UCSC genome browser shows that Atf3 binding and ATAC-seq regions are accessible for mouse Atf3 and human ATF3, see https://genome.ucsc.edu/s/mdrcetin/mm10_Atf3 and https://genome.ucsc.edu/s/mdrcetin/hg38_ATF3, respectively.

**Table 2.**
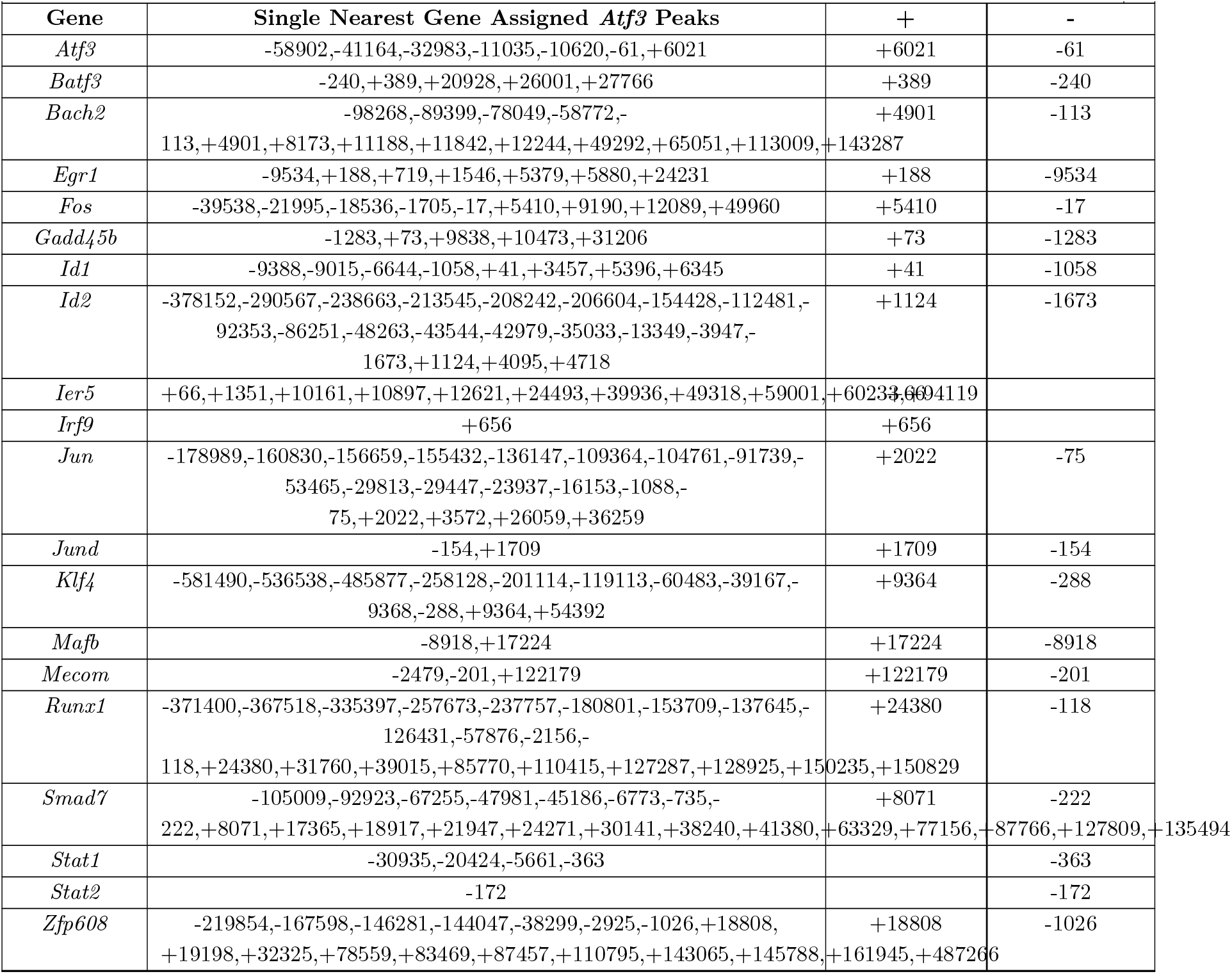
*Atf3*-KO Selected DEGs and Neighboring *Atf3* Peaks with GREAT Single Nearest Gene Assignment.

**Table 3.**
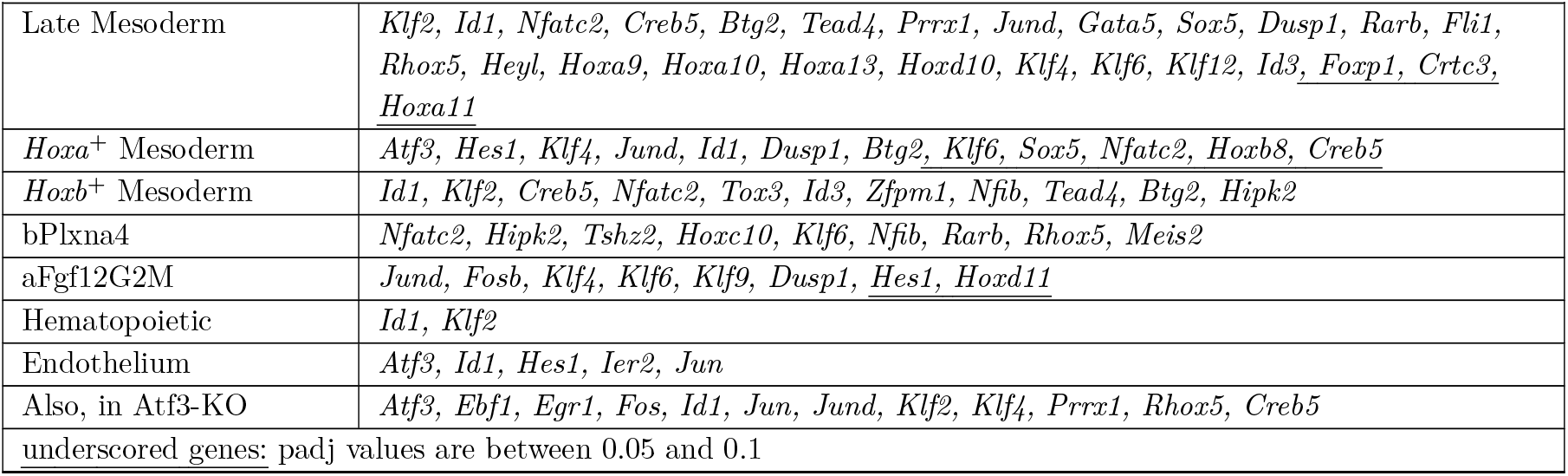
*Zfp711*-KO cells: selected DEGs (TFs and other important genes)

**Table 4.** *Zfp711*-KO cells: selected DEGs and Neighboring Zfp711 peaks with GREAT Single Nearest Gene Assignment.

|  |  |  |  |
| --- | --- | --- | --- |
| Gene | Single Nearest Gene Assigned *Zfp711* Peaks | + | - |
| *Atf3* | -41032,+104 | +104 | -41032 |
| *Btg2* | -299,+94,+18070,+18425 | +94 | -299 |
| *Cited2* | -777,-47,+300 | +300 | -47 |
| *Creb5* | +186 | +186 |  |
| *Egr1* | +34,+27443,+27827 | +34 |  |
| *Gata5* | -151,+19 | +19 | -151 |
| *Hbegf* | -305,-160,+85,+455 | +85 | -160 |
| *Hes1* | -119243,-118817,+17,+1052,+5161 | +17 | -118817 |
| *Heyl* | +51 | +51 |  |
| *Hoxa10* | +808 | +808 |  |
| *Hoxa13* | -272,+570,+966 | +570 | -272 |
| *Hoxa9* | -3196,-2299,+1292 | +1292 | -2299 |
| *Hoxc10* | +68 | +68 |  |
| *Hoxd11* | -103470,-26354,-26154,-14319,-13794,-9459,-2524,+61,+1402 | +61 |  |
| *Jun* | -226412,-109275,-99651,-24349,-987,-514,-310,+64 | +64 | -310 |
| *Junb* | +27 | +27 |  |
| *Jund* | -241,+211 | +211 | -241 |
| *Klf2* | -258,+194 | +194 | -258 |
| *Klf6* | -336,+110 | +110 | -336 |
| *Pard6b* | -601,+300,+34150,+36345,+36608,+47529,+47817 | +300 | -601 |

### 63 DEGs are shared in multiple datasets in *Atf3*- KO differentiated cells

The “ENCODE Transcription Factor Targets” dataset from the Harmonizome 3.0 database (Rouillard et al. 2016; Consortium 2004, 2011) was used to intersect DEGs after converting ATF3 targets from human to mouse orthologs (Supplemental Table S6C). In addition, existing *Atf3* knockout or knockdown data (see Mat&Meth and Supplemental Table S8A) were intersected with our DEGs list (Supplemental Table S6A). Finally, all three intersections were combined (Fig. 5E; Supplemental Fig. S13; Supplemental Table S4A) and showed that 63 out of 310 genes are present in all three data sets. They are therefore very likely direct targets of Atf3, and include *Egr1, Id1, Id2, Irf9, Smad7, Stat1, Trim25*, and the bZIP-protein encoding genes *Fos, Bach2, Batf3, Jun, Jund* and *Atf3* itself.

### EMP and Late Mesoderm Gene Set Enrichment Analysis

Next, a Gene Set Enrichment Analysis (GSEA) was carried out on EMP and Late Mesoderm. Three gene sets were used for GSEA from The Molecular Signatures Database: (MSigDB, https://www.gsea-msigdb.org/, Hallmark, Gene Ontology/Biological Processes, and Reactome) (Subramanian et al. 2005; Mootha et al. 2003). The gene sets that were affected in the GSEA (padj < 0.001) were categorized into six main categories: Cellular Processes and Cell Structure, Metabolism, Nucleus-DNA-Chromatin, Development, Immune-Myeloid System, and Signal Transduction/Pathways, respectively.

The top-GSEA results for the EMP cluster gave significant insights in cell cycle, RNA, ribosome, interferon-alpha and gamma related pathways. They exhibited highly statistically significant positive normalized enrichment scores (NES). Conversely, apoptosis, hypoxia, cellular import, endocytosis, and neutrophil degranulation-related pathways showed negative NES (Fig. 6A-C; Supplemental Fig. S14A-C; Supplemental Table S5A-C). A diverse range of findings was observed within the “Cellular Processes and Structure” category (Supplemental Fig. S15A).

**Figure 6.**
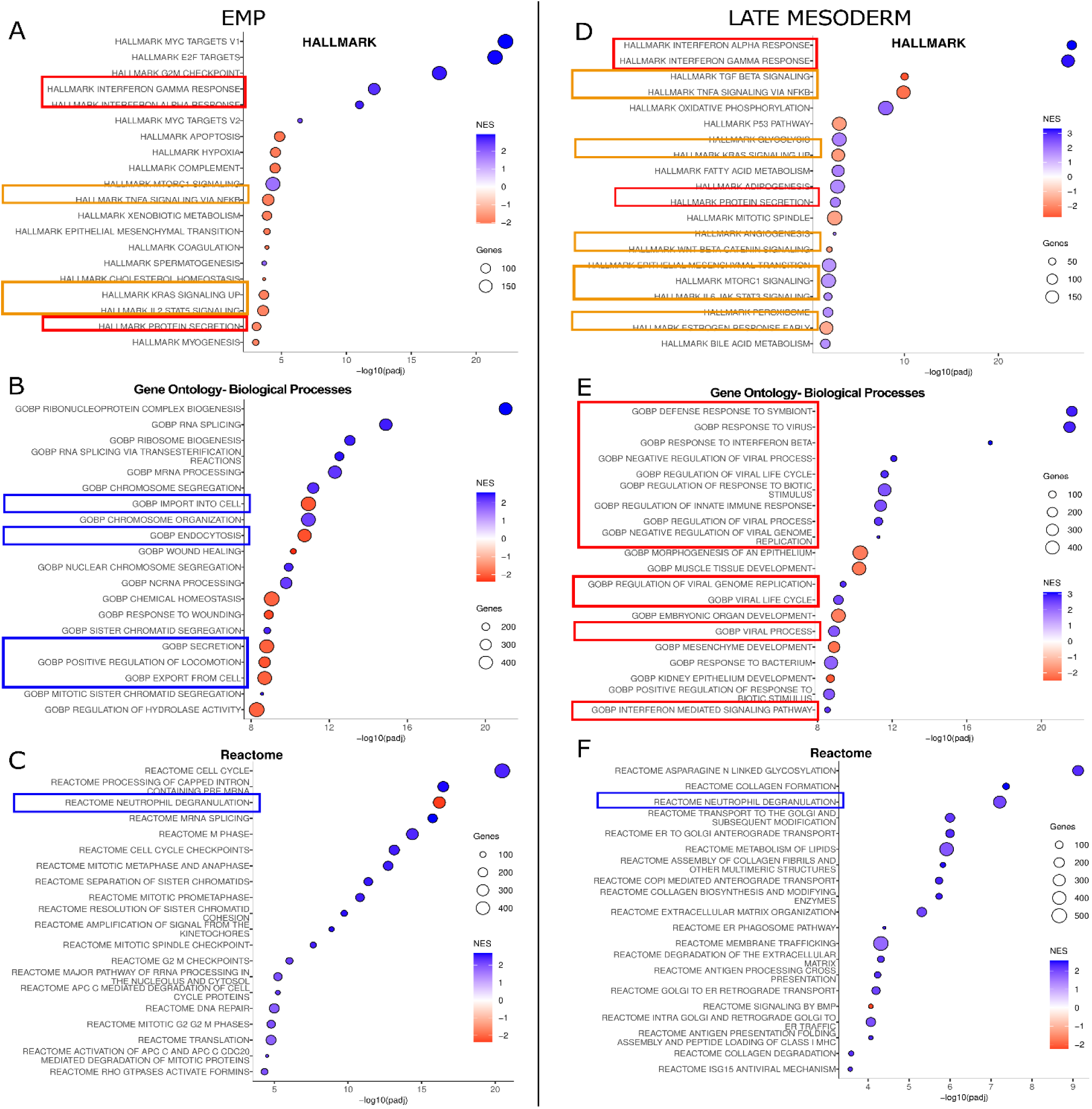
GSEA results of the differentiated *Atf3*-KO cells on EMP and Late Mesoderm. **Panels A-C** show the significance (−log10 (padj)) of the GSEA for three collections of gene sets (from MSigDB [https://www.gseamsigdb.org/]) in EMP and the top 20 gene sets in that collection (on the left). The size of the circle indicates the number of genes in a gene set (after removal of the genes that are expressed at a very low level). Blue to red indicates the increase or decrease of the Normalized Enrichment Score. Red-colored boxes highlight “interferon and antiviral related pathways,” blue-colored boxes highlight “degranulation, cellular transport, and secretion related pathways,” and orange-colored boxes highlight “signaling pathways.” **Panels D-F** show, on Late Mesoderm, GSEA results similar to panels A-C.

The developmental category, hematopoiesis, angiogenesis, and epithelial-to-mesenchyme transition each showed negatively-enriched gene sets. In the Immune and myeloid categories, two groups showed positive enrichment, “interferon-related pathways” and “response to viruses.” Other groups in this category showed negative enrichment, such as “myeloid lineages”, “cell type differentiation”, “migration” and “activation”, “phagocytosis”, “IL1B, IL2-STAT5 pathway”, “IL1, IL6, and cytokine production” (Supplemental Fig. S15A).

The positively enriched sets in the “Signal Transduction and Pathways” category included “E2F targets”, ‘MYC targets”, “GTPase-mediated signaling”, and “Runx2 and Runx3 expression’. The negatively enriched genesets included “Receptor tyrosine-kinase related signaling”, “ERRB signaling”, “MAPK Cascade”, “ERK1 and ERK2 Cascade”, “TNF signaling”, “TNF signaling with NFKb”, “KRAS”, “PI3K/Akt pathways”, “Apoptotic signaling”, “GPCR signaling,” and “antigen-receptor mediated signaling” (Supplemental Fig. S15A).

A summary of the EMP cluster in differentiating *Atf3*-KO cells showed profiles of impaired migration, reduced membrane import and export, reduced lipid and carbohydrate metabolism, reduced myeloid properties and some immune functions. In contrast, the activated response to virus infection, viral replication, and interferon pathways increased. Nuclear activity, such as cell cycle related processes, RNA-production and DNA-replication also increased (Supplemental Fig. S15A).

### *Atf3*-KO-derived Late Mesoderm

The top-GSEA (Fig. 6D-F; see also Supplemental Table S5D-F) results showed differences between various gene sets, with some replaced by others. For instance, in the EMP cluster, “cell cycle”, “RNA”, and “ribosome-related” gene sets were replaced by “Interferon-alpha and gamma”, “TGF” and “NFB (TNF-mediated) signaling” and “responses to symbiotic and viral stimuli” sets (Fig. 6D-F).

The “cell junction-related” gene sets were negatively enriched (Supplemental Fig. S15B; see also Supplemental Fig. S14D-F). The “Developmental” category revealed a wide diversity, surpassing EMPs in its scope, comprising angiogenesis, cardiac, mesenchyme, EMT, epithelium development, myogenesis, neurogenesis, osteoblast development, gastrulation, mesoderm and somatic tissue development, sensory organ, and urogenital development. These all showed negative enrichment, suggesting delayed maturation (Supplemental Fig. S15B).

As observed for EMP, the “Immune system” category shows positive enrichment in the “Interferon-alpha, beta, and -gamma” gene sets, as well as in “responses to viruses and viral replication processes,” the “Isg15 antiviral mechanism”, “antigen processing”, and “host defense”. Notably, myeloid and interleukin gene-sets showed no enrichment (Supplemental Fig. S15B; see also Supplemental Fig. S14D). In the “Signal transduction” category, positive enrichment was evident in Hedgehog family signaling, Erad, and Runx2 (gene) expression and activity, whereas negative enrichment was observed in the MAPK, Erk1-Erk2 cascade, TNF-signaling, serine/threonine-kinase signaling, including TGF-BMP and downstream SMAD, and apoptotic signaling, and P53 (Supplemental Fig. S15B).

In summary, the GSEA of the Late mesoderm profile revealed active cellular processes with increased collagen and extracellular matrix activity, along with elevated endoplasmic reticulum (ER), Golgi apparatus, and lysosome activities. The metabolic activity was increased, particularly in lipid and carbohydrate metabolism, with elevated glycolysis, glycosylation, and aerobic respiration. Nuclear activity and developmental pathways were slowed down, whereas the immune system was active in responding to related stimuli, primarily via interferon signaling.

### ZFP711

Both *Zfp711* alleles were knocked out by using Crispr/Cas9 and gRNAs by deleting 19 kb in total, encompassing exon-3 to the terminal coding exon (Fig. 7A), resulting in transcripts encoding only the N-terminal part of Zfp711, which - if translated - now lacks the DNA-binding domains. scRNA-seq (at D7) showed for *Zfp711* the highest expression at cluster level in Endothelium, followed by *Hoxb*^+^ Mesoderm, as sub-clusters EndRsad2, EndCdh5 and bStard8 (see Fig. 23, column 22; see also Supplemental Fig. S4C, pink box; Supplemental Table S9B).

**Figure 7.**
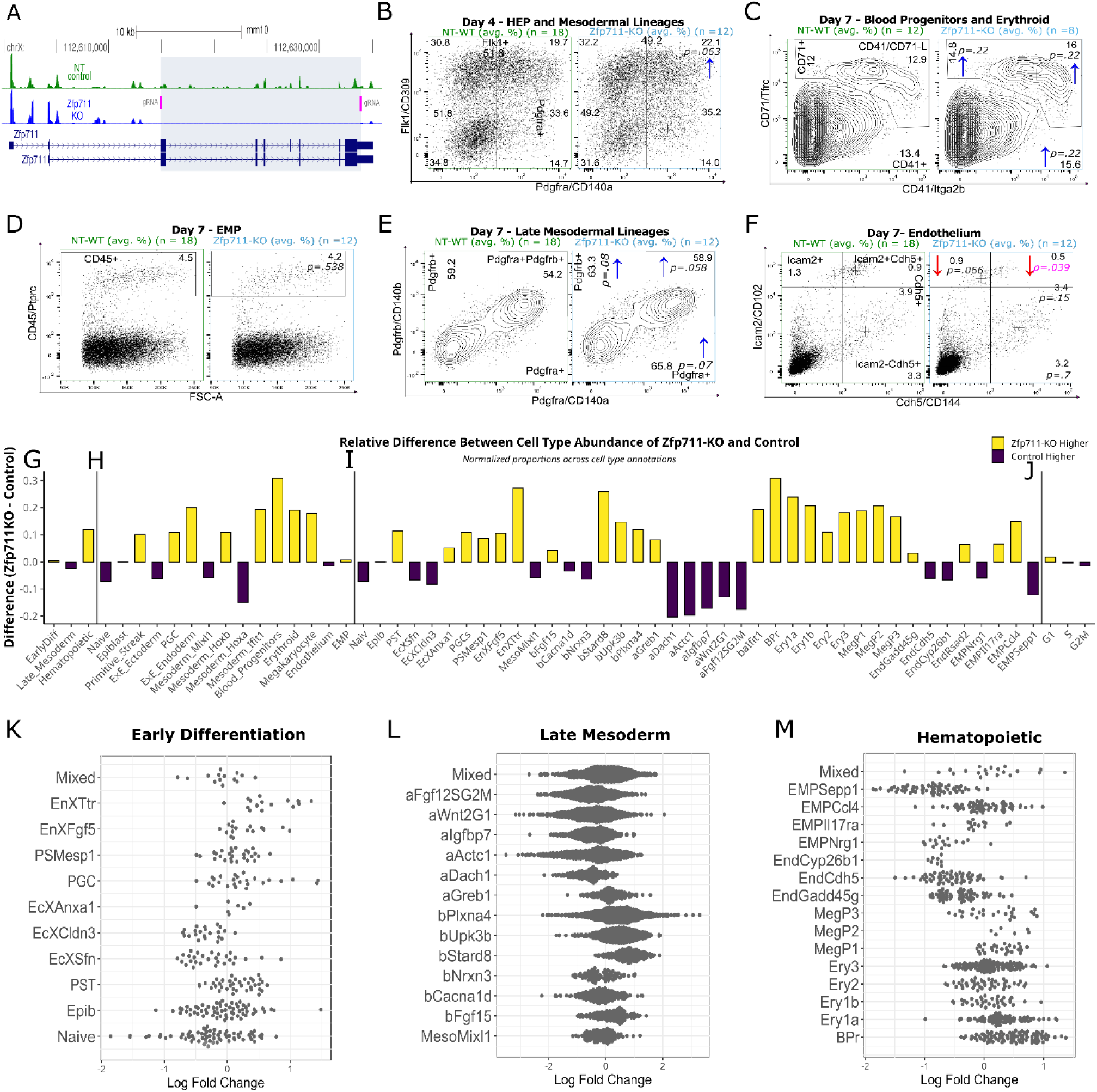
*Zfp711*-KO cells: results from Flow Cytometric Analysis and Differential Abundance Analysis. **Panel A** shows the knockout approach for *Zfp711*. The positions of the gRNAs are shown in pink. The peaks represent the scRNA-seq result for the control mESC in green and the KO in blue. The genome map of *Zfp711* is shown in dark blue. **Panel B** shows examples of FACS plots (using Pdgfra and Flk1 as cell surface markers for sorting) from *Zfp711*-KO and NT-Control cells after four days of differentiation. The numbers on the four corners show the average percentage of the respective quadrant. The numbers in the middle, right, and top show the percentage of Pdgfra- or Flk1-positive cells. The arrows indicate an increase (blue) or decrease (red) in the percentage of cells in a quadrant, and its significance. Panels **C-F** shows similar plots for D7 Blood progenitors - Erythroid cells, EMPs, Late Mesodermal lineages, and Endothelium. **Panels G-I** compare the number of NT-Control cells versus the *Atf3*-KO cells as groups, clusters, and sub-clusters. **Panel J** shows the comparison of the number of cells in the respective phase of the cell cycle. **Panels K-M** show the fold-change in the Differential Abundance Analysis in the different groups while creating small neighborhoods (small groups of cells) and are shown as dots (each dot representing in most cases 50-100 cells). Spatial FDR < 0.1, higher values are shown in grey. The text on the y-axis indicates sub-clusters, and the log fold change is shown on the x-axis. “Mixed” on the y-axis indicates a neighborhood that does not belong to any sub-cluster (using a 50% threshold).

### Zfp711-deficiency affects Late Mesoderm, Endothelium, and BP-Erythroid cells

Surface marker staining and FCA on D4 showed increases of 12% (NT-Control = 19.7 versus Zfp711-KO = 22.1 (19.7/22.1)) in Flk1^+^Pdgfra^+^ cells, 4.8% (30.8/32.3) in Flk1^+^Pdgfra^-^ cells, and 4.7% (33.7/35.2) in Pdgfra^+^ cells as compared to D4 control cells (Fig. 7B; see also Supplemental Fig. S16; Supplemental Table1C, D). At D7, the differentiated *Zfp711*-KO ESCs showed an increase of 9.9% (39.6/43.5) in the Pdgfra^+^Pdgfrb^+^ cells in Late Mesoderm and a decrease of 43% (0.85/0.48) in Cdh5^+^Icam2^+^ Endothelium. Interestingly, the BP- Erythroid cells also increased (as indicated by CD41 16.2% (13.4/15.6), CD71 23.6% (11.9/14.8) (Fig. 7B- F; see also Supplemental Fig. S16; Supplemental Table S1C,D).

scRNA-seq data for Zfp11 transcripts over the various annotated cell populations were checked similarly to those of Atf3 (Fig. 7G-M; see also Supplemental Figs. S17 and S18; Supplemental Tables S2I-P and S3D-F). At the group level, there was an increase of 27.2% (12.4/15.8) in hematopoietic lineages and a decrease of 4.5% (77.4/73.9) in the Late Mesoderm (Fig. 7G). At the cluster level, whilst ignoring the early-differentiation cell group, an increase was seen in Blood Progenitors of 89% (1.11/2.11), Erythrocytes 47.2% (4.17/6.15) and megakaryocytes 44% (1.02/1.47) (Fig. 7H). A 26.1% (44.3/32.7) decrease is observed in *Hoxa*^+^ mesoderm, whereas *Hoxb*^+^ mesoderm increased by 24.4% (33.0/41.1) (Fig. 7H; see also Supplemental Fig S17B; Supplemental Table S2I,N).

At the sub-cluster level, the aforementioned decrease was confirmed for *Hoxa*^*+*^ cells, i.e., in “aIgfbp7” (29.2%), “aFgf12SG2M” (29.8%), “aWnt2G1” (22.9%), “aActc1” (32.9%) and “aDach1” (33.7%) cell populations, while the increase was also confirmed for some *Hoxb*^+^ sub- clusters, i.e. for “bPlxna4” (27.04%), “bUpk3b” (34.4) and “bStard8” (69.7%). For the Hematopoietic Cells, there are increases for Erythroid and Megakaryocyte sub-clusters “BPr” (89.4%) and “Ery3” (44.6%). Some Endothelial and EMP sub-clusters like “EMPSepp1” (21.5%) and “EndCdh5” (11.4%) show a decrease (Fig. 7I,K-M; see also Supplemental Fig S17C; Supplemental Table S2J,O)), whereas there was no change in the total cell cycle phase (Fig. 7J; see also Supplemental Fig. S17D).

### Effects of *Zfp711* inactivation on *Hoxb*^+^ and *Hoxa*^+^ mesoderm

The highest numbers of DEGs (padj < 0.05) mapped to the Late Mesoderm group (Fig. 8A). The sub-cluster “bPlxna4” has the next-highest DEGs, followed by *Hoxb*^+^ Mesoderm. In Late mesoderm, 96 genes were downregulated, and 43 were upregulated. The Hematopoietic clusters and sub-clusters showed only small numbers of DEGs (Fig. 8B; see also Supplemental Table S4L-T).

**Figure 8.**
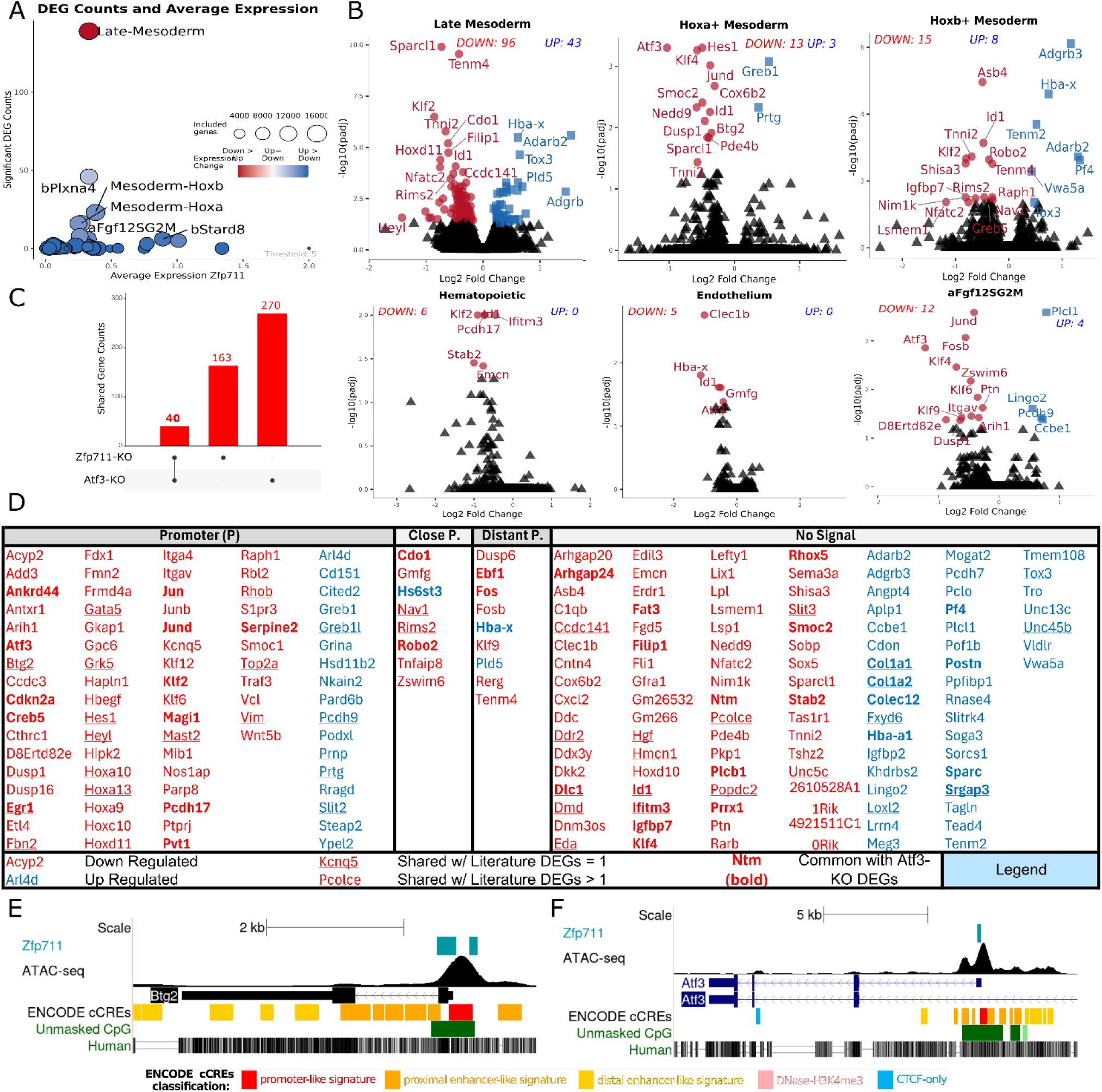
Results and categorization of the Differential Gene Expression Analysis of the *Zfp711*-KO cells. **Panel A** shows the average expression level and significant DEGs (padj < 0.05) in groups, clusters, and sub-clusters. The size of the circles indicates the number of genes included in the analysis after removing low-expressed genes in the group, cluster, or sub-cluster. The circle’s color indicates up- or downregulation (*blue* and *red* respectively). **Panel B** shows the up- or downregulation of genes in the Endothelium and *Hoxa*^+^ mesoderm clusters and Late Mesoderm group. **Panel C** shows the number of DEGs shared between *Atf3*-KO and *Zfp711*-KO. **Panel D** shows the total list of Significant DEGs from pseudobulk scRNA-seq at day 7, which was categorized/visualized with Zfp711 binding sites, up-downregulation, shared *Atf3*-KO DEGs list, and shared with ZNF711-related literature DEGs. **Panel E-F** shows two examples of Zfp711 binding sites with ATAC-seq peaks in three examples of DEGs.

Effects on the *Hoxa*^+^ and *Hoxb*^+^ Late Mesoderm showed that in *Hoxa*^+^ Mesoderm and its sub-clusters a number of genes are downregulated, e.g. the TF-encoding genes *Atf3, Klf4/6/9, Hes1, Jund, Fosb, Zswim6*, and the growth/proliferation-related genes *Dusp1, Hbegf*, whereas *Hoxb*^+^ Mesoderm did not show such changes. Instead, *Hoxb*^+^ Late Mesoderm showed decreased *Klf2, Asb4*, and *Robo2*, and increased *Tox3* and *Adgrb3* (Fig. 8B). Table 3, Supplemental Tables S4L-T and S10C,D show the DEGs from various clusters that can affect gene regulation and include genes like *Id1, Atf3, Jun, Ebf1, Fos*, and *Egr1*.

### Characterization of the *Zfp711*-KO DEGs

In order to functionally categorize the intact-*Zfp711* dependent DEGs, we repeated a similar processing as for intact-*Atf3* dependent DEGs (a total of 203 genes (padj <0.05)). Briefly, ZNF711 ChIP-seq data were collected (see Material Methods and Supplemental Table S8B), converted to the mouse genome, and intersected with ATAC-seq and *Zpf711* motif-containing genomic regions. Regions appearing positive in these three assays were selected and analyzed to establish whether they were located within or close to individual DEGs.

Most Zpf711 binding sites are located downstream of a TSS (defined here as up to +500 bp; Supplemental Fig. S19A,B). Using this information and visual confirmation of human and mouse genomic locations, the DEGs again fell into four discernable categories (Supplemental Table S7F-J, S4L). These were “Promoter” and “Close Promoter” (− 300 to +500 bp), “Distant Promoter” (−300 bp to −2Kb), and No Signal (−2kb to the next gene) (Fig. 8D, see e.g. panels 8E and 8F, Table 4). DEG lists from published *ZNF711* knockout and *ZNF711* knockdown studies were also collected; they overlapped with the DEGs in this study (albeit following conversion to the mouse Znf711 ortholog (Fig. 8D; see also Supplemental Fig. S19; Supplemental Table S6B). Interestingly, *Atf3* was among the DEGs and has a Zfp711 binding site in its promoter (Fig. 8F). The DEG list of the *Zfp711*-KO and *Atf3-*KO differentiated ESCs shared 40 genes (30 of which are up, 10 are down), including *Atf3* itself (Fig. 8C; see also Rhie et al. 2018). Noteworthy, Rhies et al. (2018) only showed an upregulation of *ATF3* in cell lines, but did not look into stem cell-derived development.

### GSEA of *Zfp711*-KO cells in Late Mesoderm

The GSEA of *Zfp711*-KO of Late Mesoderm showed a decrease of “TNF signaling with NFB”, “Interferon-Gamma Response”, “Estrogen Late and Early response”, “Apoptosis”, “Cholesterol metabolism”, and “Somatic tissue development”. “RNA/ribosome-related pathways”, “Intracellular transport-related pathways”, and “Glycosylation” were enriched (Fig. 9, see also Supplemental Table S5G-I, Supplemental Fig. S20). The KO of *Zfp711* caused the downregulation of *Atf3*; however, rather than an expected upregulation of the “Interferon-related pathways,” as we observed for the *Atf3*-KO cells, these pathways were downregulated. This difference is likely due to other factors, such as *Bst2* and *Ifitm3*, which are upregulated in the *Atf3*-KO, but downregulated in the *Zfp711*- KO cells (Fig. 5B,H).

**Figure 9.**
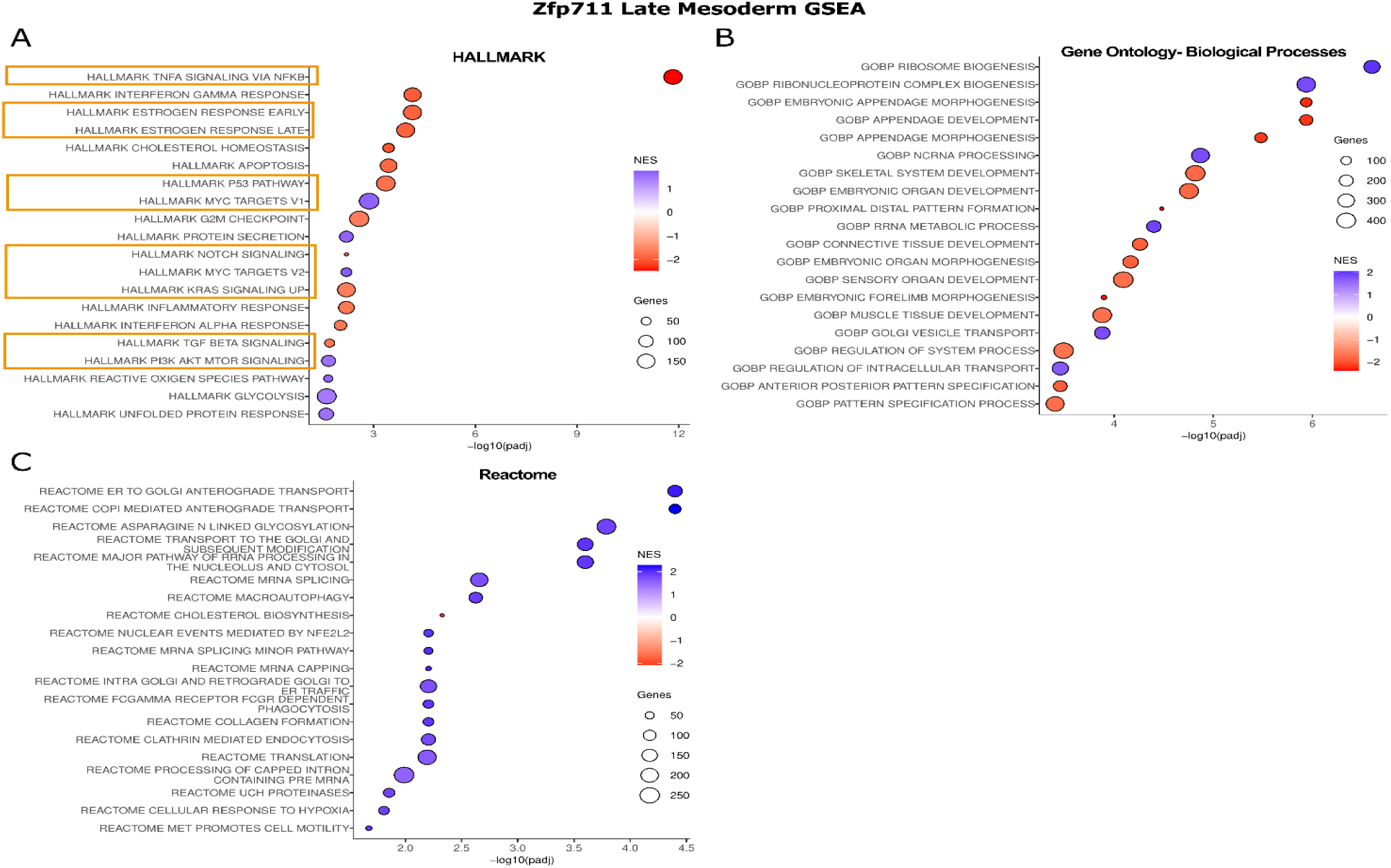
GSEA Results of the *Zfp711*-KO cells as Late Mesoderm. **Panels A-C** show the significance (−log10 (padj)) of the GSEA for three collections of gene sets (from MSigDB, https://www.gseamsigdb.org/) and the top-20 gene sets in that collection (on the left). The size of the circle indicates the number of genes in a gene set (after removal of the genes that are expressed at a very low level). Blue to red shading indicates the increase or decrease of the Normalized Enrichment Score (NES). The orange-colored boxes highlight “signaling pathways.”

## BCL6B

### *Bcl6b*-KO cells show no observable changes

*Bcl6b* was inactivated as shown in Supplemental Fig. S21A. *Bcl6b*-KO cells showed minimal changes compared to controls in terms of cell composition, as determined by scRNA-seq (Supplemental Fig. S21B-H, S23, and S24; Supplemental Tables S2Q-X and S3G-I). Surface marker staining (Supplemental Fig. S22; see also Supplemental Table S1E,F) did not change significantly (Supplemental Fig. S21B-J). The DGEA of groups, clusters, and subclusters, respectively, showed only 18 genes affected by the KO of *Bcl6b* (Supplemental Fig. S21I,J; see also Supplemental Table S4U,V). These KO cells do not appear to display any statistically significant change in the tissue where *Bcl6b* is expressed highly, i.e. only a small decrease was seen in Endothelium (Supplemental Fig. S21C).

## DISCUSSION

### Technical Aspects of the Analysis

This study shows that single-cell multiplexing technology can be used to study continuous cell differentiation as an entire system, with minimal batch effects. At the same time, it allows the investigation of multiple samples with biological monoclonal replicates. However, several potential technical problems should be kept in mind. Importantly, during the demultiplexing step of the scRNAseq analysis, we observed that Naive Pluripotent ESCs have a lower CMO labeling percentages than other cell types. Another observation is that Erythroid cells and, to a lesser extent, EMPs failed to end up in a single droplet more often than other cell types (see Supplemental Fig. S25). It is not clear why, but perhaps the changes in the plasma membranes of blood cells cause them to interact more with each other, in particular after cell labeling.

Another note is that particular annotated cell identities or characteristics may get lost during the extended labeling procedure. For example, different cardiac cell types could not be distinguished in late mesodermal lineages even though the D7 EBs, as observed under the microscope, were pulsating. In contrast to our previous D7 scRNA-seq analysis of the source mESCs (129×1/Svj origin, see Material Methods; but *without* multiplexing), we identified such cardiac cell types (see Supplemental Fig. S2A-C). A potential explanation for this difference could be the extended duration of the dissociation of the EBs during the capture of the single cells for scRNA-seq, as well as their labeling, causing a collective loss of recognizable cardiac cells.

Another technical step that warrants attention is the quality control of scRNA-seq itself. The potential strategy of using the ratio of “detected gene number” to “unique RNA-molecule count” can lead to discarding highly-specialized cells, especially when present in low numbers. We discarded the use of this ratio for preprocessing (filtering low RNA and gene counts, doublet removal, etc.), because Erythroid lineages showed a lower “detected gene number” to “unique RNA-molecule count” ratio than other clusters. It is important to consider the potential presence of lineages with a large bias in gene expression, i.e., Erythroid cells with a large bias of globin mRNAs versus the rest of the transcriptome. Such cells are filtered out when this ratio is used in the quality control of single cells.

Yet another potential problem is the presence of ambient RNA even after using removal tools for them (e.g., in Late Mesoderm Hba-x is present where it should be absent). An interesting observation is the presence of the “MegP2” sub-cluster. This cell population expresses genes that are seen in the advanced stages of Erythroid and Megakaryocyte Progenitors, suggesting they may represent a separate cell identity; however, we cannot firmly exclude that they simply would result from leftover ambient RNA.

DGEA frequently returns top-upregulated genes such as *Fam71a* for the *Atf3*-KO cells, *Pof1b* for *Zfp711*-KO, and *Slc16a13* for *Bcl6b*-KO. The common feature of these three genes is that they are known to be epistatic (here: downstream). Thus, their increased expression may be caused by the deletion of part of the neighboring DNA, causing changes in the local dynamics of the transcription (e.g., read-through, deletion of a regulatory region, reduced distances between promoters and upstream enhancers).

### Atf3

The *Atf3*-KO resulted in the most significant change in this very early differentiation by affecting cell type abundance and leading to the highest number of DEGs (Fig. 10A) when compared with the other two KO’s (see below) (Fig. 10B,C). The differentiation of *Atf3*-KO ESCs showed a change in cell composition at D4 and D7 (see Fig. 4) and gene expression (Fig. 5), particularly in the EHT of the EMPs and Endothelium. DGEA showed many more downregulated than upregulated genes in EMP lineages (see Fig. 5A,B). The “Late mesoderm” showed the opposite trend. Thus, the effects of the *Atf3*-KO can be separated into two categories: the Late Mesodermal and the Hematopoietic lineages, respectively. There is an increased differentiation towards “Late Mesodermal lineages” by a higher percentage on D4 of Pdgfra^+^ cells and on D7 of CD140a^**+**^CD140b^**+**^ cells (see Fig. 4). The number of EMPs (CD45^**+**^) decreased due to a delay in EHT transition or maturation, which gets worse during further EMP differentiation. The downregulation of *Runx1, Mafb*, and *Sox4* (and other genes) may explain a significant decrease in EMP lineage. For example, *Mafb* has many Atf3 binding sites, suggesting a potential direct regulatory role of Atf3. The decrease in EMP agrees with studies by Yin and co-workers (Yin et al. 2020), although they alternatively suggest that the decrease in EMPs is caused by a decrease in glucose utilization and a change in the redox state.

**Figure 10.**
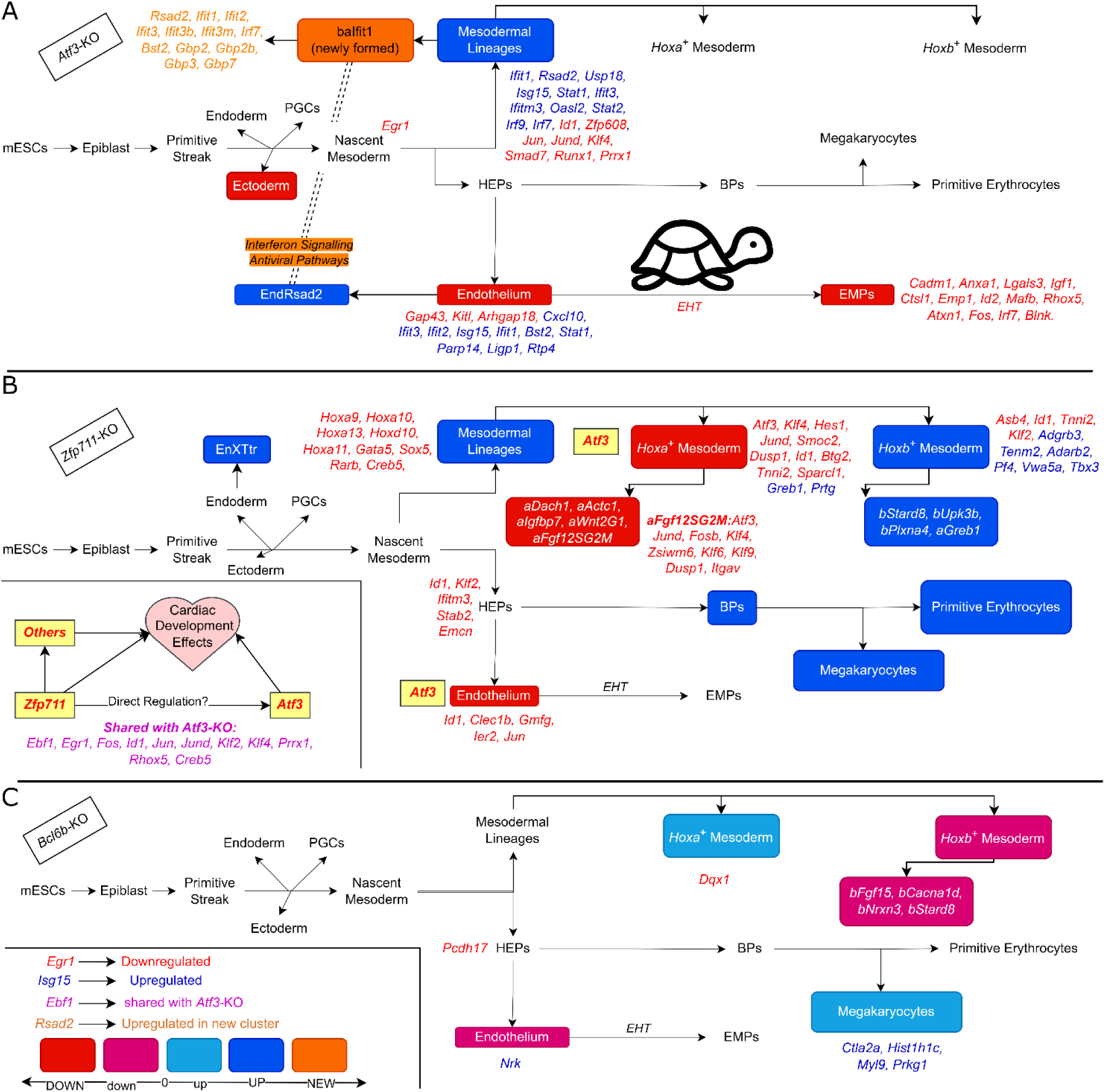
Summary of effects of Atf*3-*KO, *Zfp711-*KO, and *Bcl6b-*KO in ESCs submitted *in vitro* to hematoendothelial differentiation. **Panel A** shows detected changes in *Atf3*-KO on the developmental trajectory diagram (Turtle indacates slowing down in EHT). **Panel B** shows detected changes in *Zfp711*-KO on the developmental trajectory diagram. **Panel C** shows detected changes in *Bcl6b*-KO on the developmental trajectory diagram.

The same team also showed an atrial dilated cardiomyopathy and an increased ratio of immature myeloid cells. Our findings on the EMP lineage overlap with their findings. Our data also show changes to Late Mesoderm, including cardiac lineages. However, the lack of clearly identifiable cardiac lineages in our scRNA-seq data regrettably prevents conclusions about specific cardiac lineages (as discussed above).

Two subsets of Endothelial cells (EndCdh5 and EndGadd45g) decreased, whereas one small cluster was highly enriched in *Atf3*-KO cells (EndRsad2), the latter probably being derived from EndCdh5, as based on mapping proximity (see Supplemental Fig. S5C,G, clusters 34 and 36). Similarly, a novel cluster appeared in Late Mesoderm (balfit1) on D7, mostly consisting of *Atf3-KO cells*, probably derived from the clusters Mesoderm *Hoxa*^**+**^ and *Hoxb*^**+**^ (Supplemental Fig. S4A,B). In addition to EMP, many other differentiation processes are affected, e.g., angiogenesis, epithelial to mesenchymal transition, cardiac, and mesenchyme development (Supplemental Fig. S15), suggesting that Atf3 plays an essential role during the development and maturation of many tissues. These effects may be due to metabolic changes upon *Atf3*-KO ESC differentiation in Late Mesoderm, as suggested by an increase in “Lipid and carbohydrate metabolism and cellular energy consumption” (Supplemental Fig. S15B). In addition, both EMP and Late Mesoderm show an increase in “Interferon and response to viruses”.

GSEA suggests that the top-hallmark results are “Cell cycle and related pathways”, which are upregulated in *Atf3*-KO EMP (Supplemental Fig. S15A), but this is not seen in Late Mesoderm. Instead, it shows “Interferon response” as the top-hallmark, while “Cell cycle gene sets” is not in the top 20 (Fig. 6 and Supplemental Fig. S14). This observation raises the question of whether the upregulation of cell cycle-related genes is the effect of Atf3 deficiency. The likely explanation for these opposite results for the two groups is that the *Atf3*-KO has much fewer cells in the “EMPsepp1” sub-cluster. Therefore, the total number of mutant cells in the G1-phase will also be lower than in the NT-Control.

The DEG approach combines the gene counts of all the cells and creates pseudo-bulk data. Next, the pseudobulk data of the NT-Control and the *Atf3*-KO are compared, ranking genes that are up- or downregulated. In GSEA, these lists are compared to curated gene-sets, the Hallmark gene-sets, the GO biological processes gene-sets, and the Reactome gene-sets. GSEA results in a significance and normalized enrichment score for each gene-set. If it gives a positive score, it suggests many upregulated genes in the particular gene set.

The late EMP cells (EMPsepp1) show that 70% of these cells are in the G1-phase (Supplemental Fig. S5B), suggesting they have matured to EMP. The *Atf3*-KO cells lead to fewer such EMPsepp1 cells (Fig. 4F,I,M). Thus, during the DGEA on EMP, the *Atf3*-KO will have fewer cells in G1-phase, suggesting that a change in the cell cycle is due to the absence of Atf3, while this needs not be the case. GSEA, therefore, indicates what processes may be affected, but a conclusion can only be made after including changes in cell type frequency before they are linked to a biological process.

In the “Late Mesoderm” group, the predominantly upregulated DEGs point to the role of Atf3 as a repressor. Notably, the TGF (/BMP) family component genes (*Id1, Smad7, Id2*) and TNF -NF B (*Ier5, Id2, Plk2, Klf4, Fos, Jun, Gadd45b*) pathways show negative enrichment. “Interferon signaling” and “Response to viruses” are upregulated alongside collagen formation in accordance with Sood and co-workers (Sood et al. 2017). Finally, positive enrichment was clear for the “Signal transduction” category, i.e., in Hedgehog family signaling, the Erad pathway and *Runx2* expression and activity pathways. In contrast, negative enrichment was observed in the MAPK, Erk1Erk2 cascade, TNF signaling, serine/threonine-kinase signaling, TGF, BMP and downstream SMADs, apoptotic signaling, and P53-related pathways, which explains why the “Development” category (Supplemental Fig. S15B) is downregulated.

### Zfp711

The main effect of the *Zfp711*-KO was apparent for Late Mesoderm (using DGEA). Effects on the “Late Mesoderm” sub-clusters differ in terms of the abundance and differences in the DEGs. “*Hoxa*^+^ Mesoderm” and its subclusters show downregulation of the TF-encoding genes *Atf3, Klf4/6/9, Hes1, Jund, Fosb, Zswim6*, and the growth/proliferation-related genes *Btg2, Dusp1, Hbegf*. “*Hoxb*^**+**^ Mesoderm” does not show this.

*Atf3* is one of the *Zfp711-*dependent DEGs in *Hoxa*^**+**^ Mesoderm. This finding is supported by Rhie and coworkers (Rhie et al. 2018), who observed *ATF3* downregulation in their *ZNF711*-KO experiments in human cell lines. ChIP-seq data in humans and mice (Supplemental Fig. S19A,B) showed that the *Atf3* promoter contains binding sites for Zfp711. Modulated Atf3-DEGs and Zfp711-DEGs also showed that another 39 genes overlap (Fig. 8C,D). The decreased expression of *Atf3* in Endothelium would explain why the number of endothelial cells has decreased (Fig. 7F and Supplemental Fig. S16).

Snabel et al. (2024) showed that when *ZNF711* is knocked out in human pluripotent stem cells that cardiomyocyte differentiation was affectedwhen the cells are differentiated (without retinoic acid). DEGs intersection showed that 37 out of 204 genes were shared between the *ZNF711*KO and the Zpf711-KO ESCs shown here, and included *Hes1, Heyl, Klf6, Id1, Jun, Tshz2, Tox3*, and some other essential genes for cardiac cells such as *Dmd* (Dystrophin) and *Unc45b* (Myosin Chaperone B).

This *Atf3*-KO and results from Yin et al. (2020), show that the cardiac lineages are among the affected mesodermal lineages. Additionally, *Zfp711*-dependent DEGs included a number of genes related to the family of nuclear retinoic acid receptors (e.g., *Rarg* (Retinoic Acid Receptor Gamma), *Rarb*), supporting the potential role of the *Zfp711* with retinoic acid in cardiac development as reported by Snabel et al. (2024). The lower level of *Atf3* expression may explain part of the Zfp711-deficiency effects on the cardiac lineage observed here. The blood lineages are not increased in the *Atf3*-KO because this KO decreases essential hematopoietic genes such as *Runx1* or *Mecom*. Their decrease is not observed in the *Zpf711*-KO.

The endothelial sub-cluster EndCyp26b1 displayed a distinct profile compared to two other smaller sub-clusters (baIfit1 and EndRsad2). A closer examination of the markers for EndCyp26b1 suggests that this sub-cluster may be guiding endothelial cells toward an endocardial fate. The increased expression of genes such as *Pdlim3, Ccdc80, Ednrb, Gata4, Hand2, Meis2, Tbx20*, and *Sox6*, along with the reduced expression of *Mef2c, Zfp711*, and *Gadd45g*, supports this observation (Supplemental Fig. S26). The downregulation of *Zfp711* during the endocardial transition suggests that *Zfp711* may play a crucial role in endothelial and cardiac lineage development (Daniel et al. 2020; Pawlikowski et al. 2019).

### Bcl6b

*Bcl6b* genetic inactivation in ESCs showed minimal effect (only 18 DEGs were found) in this cell differentiation system (using FCA and scRNA-seq). In fact, the *Bcl6b* KO samples function as a spike-in of the analysis of *Zfp711*KO and *Atf3*-KO cells.

### Conclusion

Adaptation of our multiplexing scRNA-seq technology to investigate three TF-encoding genes within one experiment, with multiple biological replicates of *in vitro* differentiation, successfully captures cell types from the naïve pluripotent state to differentiated states without observable batch effects. This again shows the strength of the differentiation of mouse ESCs to mimic developmental steps that occur *in vivo*. The presence of Erythroid, Megakaryocytes, Endothelium, and transitioning cell types like EMPs and Endocardium (EndCyp26b1) highlights the presence of the rich cell type environment of the *in vitro* differentiation system, which is directed without additional signaling molecules other than ascorbic acid and transferrin (see Material Methods), suggesting that a similar approach can be used in other directed differentiation systems.

We developed a workflow combining data from ATACseq, mRNA expression analysis at the single-cell level, and publicly available ChIP-seq to study the role of three TFs during in early development. We asked three questions: **1**) Are these three TFs important for early lineage choices between hemato-endothelial lineages and late mesodermal lineages? **2**) Are they important for later stages of the differentiation? 3) Which genes and pathways are changed upon their perturbation?

***Atf3*** clearly has a role in the early stages of differentiation by increasing the late mesodermal lineages. At later stages, it regulates the number of EMPs. Its absence delays EHT and forms novel clusters in Late Mesoderm. Affected pathways and genes, include downregulation of genes like *Runx1 or Mafb* and signaling pathways such as TGF /BMP and TNF -NF B pathways, while it up-regulates interferon signaling and response to viruses. ***Zfp711*** has a role in *Hoxa*^***+***^ Mesoderm and endothelial cells. Its absence affects endothelial lineages and Late Mesodermal lineage derivatives such as cardiac lineages, and interestingly, it down-regulates *Atf3* and other genes that are also downregulated in the *Atf3* KO. ***Bcl6b*** appears to have almost no role (Fig. 10C), although its absence was shown to affect neovascularization in ocular vascular diseases (Tanaka et al. 2023), a tissue that is absent in the mESC system.

Multiplexed scRNA‐seq and *in vitro* differentiation reveal distinct roles for Atf3 and Zfp711 in the early hemato‐endothelial and mesodermal specification, underscoring the versatility of our integrative workflow. We hope these insights will inspire further exploration of embryogenesis and spark innovative strategies in regenerative medicine.

## MATERIAL AND METHODS

### Cell Line Generation with CRISPR-Cas9 knockouts

Two guide RNAs were designed to delete each of the three genes using the IDTCustom Alt-R™ CRISPR-Cas9 guide RNA tool, including two non-targeting gRNAs. “IDT’s “Alt-R CRISPR-Cas9 System: Cationic lipid delivery of CRISPR ribonucleoprotein complexes into mammalian cells” guideline (version 6) followed downstream analysis with Alt-R® CRISPR-Cas9 tracrRNA, ATTO™ 550 (1075927), Alt-R™ CRISPR-Cas9 crRNA, Alt-R™ S.p. Cas9 Nuclease V3, (1081058), and 1075927 Lipofectamine™ CRISPRMAX™ Cas9 Transfection Reagent (CMAX00008) were used for the transfection of mESCs. Transfected cells were sorted after 24 hours into 96-well plates, which are MEF coated and contain 2i+mESCs Medium. After 5-7 days, each well was checked for monoclonal growth, transferred into a new 96-well plate, and two copies of a 96-well plate were prepared; one plate was frozen down within 10% DMSO in FBS, and the other 96well plate was used for genomic DNA isolation. Following genotyping of the candidate clones with PCRs, KOpositive clones were thawed, expanded, genotyped again, and frozen in multiple vials.

The following guide RNAs were used to target;

Atf3 GGGACTAGTTTTCACAACGCTGG and TCATTATTGAGGTTGTCCAATGG, Bcl6b: AGGTACCAGACTTTACCTGGGGG and AAGGCCCTGGTACCAACTACAGG, Zfp711: GATGGGATAACTCTCGATCATGG and GTTAAGACACCTTATTTGATTGG, NT-Control: IDT-control crRNAs (1072544, 1072545).

The primers used for genotyping PCR reactions of Atf3, Zfp711, and Bcl6b were as follows. For Atf3, the primers were Atf3_out_f5_#601 (CTGGAGACCTCTGTGCAAGA), Atf3_out_r3_#602 (TGCCCTGTCACTGAGTATGG), Atf3_ins_f2_#604 (ACAGCTTGCCATGAAACCTG), and Atf3_ins_r1_#605 (CTCCTCAATCTGGGCCTTCA). For Zfp711, the primers used were Zfp711_out_f4_#577 (CCTGGTTTTGGCCTTACAGA), Zfp711_out_r2_#578 (TGCATTTTCTTTCCCCACCC), Zfp711_ins_f2_#580 (AAACTGCCGAACAAGGACTG), and Zfp711_ins_r2_#581 (TCACAATGCCTACACTGGTG). Finally, for Bcl6b, the primers were Bcl6b_out_f3_#583 (CTCATCCTCGGGTGCTTAT), Bcl6b_out_r3_#584 (TCAAACCCAGCTGTTCATGC), Bcl6b_ins_f2_#586 (GGCAGTTCTTATCGCTTGCA), and Bcl6b_ins_r1_#587 (ACACCGTCTCCTAGTCGTTG).

### Media

mESCs-Medium: DMEM HyClone™ (cytiva SH30081), 15% Fetal Bovine Serum (FBS) (Capricorn Scientific FBS-12A, CP18-2152), 1X MEM Non-Essential Amino Acids Solution (100X) (Gibco 11140-035), 2mM GlutaMAX™ Supplement (Gibco 35050061), 10mM HyClone™ HEPES Buffer (cytiva SH30237.01), 1X Penicillin-Streptomycin (100X) (Sigma-Aldrich P0781), 0.1mM 2-Mercaptoethanol (50 mM) (Gibco 31350010), 1000U.ml - ESGRO® Recombinant Mouse LIF Protein 10^7 units/ml (Sigma-Aldrich ESG1107).

MEF-Medium: DMEM (Capricorn Scientific DMEM-HPSTA), 15% - FBS, (Capricorn Scientific FBS-12A, CP18-2152), Penicillin-Streptomycin (100X) (Sigma-Aldrich P0781), 2mM - GlutaMAX™ Supplement (Gibco 35050061).

2i+mESCs-Medium: mESCs-Medium with 3 μM - CHIR 99021 (biotechne-TOCRIS 4423) and 1 μM PD0325901 (selleckchem S1036).

Freezing Medium: FBS (Capricorn Scientific FBS-12A, CP18-2152) with 10% DMSO (general lab consumable)

Dissociation Buffers: Trypsin-EDTA (0.05%), phenol red (Gibco 25300054) or TrypLE™ Express Enzyme (1X), phenol red (Gibco 12605010)

Differentiation Medium: IMDM, GlutaMAX™ Supplement (Gibco 31980048), 15% FBS (Capricorn Scientific FBS-12A, CP18-2152), 1X - Penicillin-Streptomycin (100X) (Sigma-Aldrich P0781), 0.1 mM - 2-Mercaptoethanol (50 mM) (Gibco 31350010), 50 μg/ml - L-Ascorbic acid (Sigma-Aldrich A5960-25G), 150 g/mL -Transferrin (Roche 10652202001)

Wash Buffer: 10% FBS (Capricorn Scientific FBS-12A, CP18-2152) in PBS (Capricorn Scientific PBS-1A or Gibco 14190-094)

### mESCs and *In Vitro* Differentiation

129 X1/Svj mESCs (The Jackson Laboratory) were grown on irradiated mouse embryonic fibroblasts at 37°C and 5% CO2 and were split every two days. Two or three passages before the differentiation, 2i+mESCs medium was used for one passage and then switched back to the mESCs medium. When the differentiation was started, the following steps were performed: the mESCs were harvested and made into a single-cell suspension, strained with 40 μM cell strainer (Greiner 542040), washed three times, and counted in Trypan Blue (Bio-Rad 1450021) with a Countess II Automated Cell Counter. One million live cells were plated in 12 ml in a petri dish (Greiner 633102) freshly prepared differentiation medium, and plates were placed on an orbital shaker at 60 rpm, 37°C, and 5% CO2. On the second day of the differentiation, 2 ml of prewarmed differentiation medium was added to each plate. On day 4, HEP and mesodermal cell differentiation efficiency were checked using half of the embryonic bodies. The remaining half was transferred to new plates, 13 ml differentiation medium was added, and the mixture was kept in the same condition until day 7.

The harvested EBs were washed three times with PBS, and prewarmed TrypLE was added in a 50 ml falcon tube and placed in a 37°C water bath while shaking with intermittent pipetting for disassociation. Four minutes later, 3 ml Trypsin/EDTA was added, and the dissociation process continued for at least 7 minutes while visually checking the dissociation of the EB. The disassociation buffer was inactivated with PBS+10%FCS, strained with a 40 μM cell strainer to create a single-cell suspension, washed three times, and stained for cell surface marker staining or prepared for scRNA-seq.

### Surface Marker Staining and Flow Cytometric Analysis

On day 4, Flk1 and Pdgfra antibodies were used for cell type identification, while on day 7, CD144, CD102, Pdgfra, Pdgfrb, CD41, CD71, CD45 and CD93 antibodies were used. Briefly, a million cells were incubated in 100 μl PBS+10%FCS with the antibody combinations for half an hour on ice, followed by three washing steps. Stained samples were measured with BD LSRFortessa™ Cell Analyzer. The following antibodies were used following the manufacturer’s recommendation: CD309 APC (BioLegend 136405), CD309 AF647 (Biolegend 121910), CD140a PE (Biolegend 135905), CD140a PE/Cy7 (Biolegend 135911), CD140b APC (Biolegend 136007), CD102 AF647 (Biolegend 105611), CD144 PE/Cy7 (Biolegend 138015), CD41 PE (Biolegend 133905), CD93 PE (Biolegend 136503), CD45 APC (Biolegend 157605), CD45 APC/Cy7 (Biolegend 103115), CD71 APC (Biolegend 113820), CD202 PE (Biolegend 124007).

Flow cytometric analysis results were analyzed using FlowJo 10.8.1 software. Staining panel frequencies were compared to mixed-model statistical cells using the lme4 R package (Bates et al. 2015) with the setting “~ Conditions + (1|Differentiation)”.

### Differentiation for scRNA-seq and CMO Multiplexing

For scRNA-seq, three clones were selected for each condition (Atf3-KO, Bcl6b-KO, Zfp711-KO, NT-Control), and three replicate differentiation plates were generated for each clone. These replicate plates were merged on day four, and the differentiation continued until day 7. After harvesting EBs and generating a single-cell suspension, samples were washed three times and the cells were counted. The “3’ CellPlex Kit Set A” (10XGenomics 1000261) kit was used according to the manufacturer’s instructions to label the samples. The cell concentration was measured again, and samples were pooled, targeting equal cell numbers to proceed to the single-cell RNA seq capture.

### ScRNA-seq Capture, Library Preparation, and raw data analysis

scRNA-seq was performed using the 10X Genomics Chromium X Platform. The Chromium Next GEM Single Cell 3’Reagent kit v3.1(Dual indexing) kit with feature Barcode technology for cell multiplexing (10x Genomics) was used following the manufacturer’s instructions generating separate gene expression and cell multiplexing libraries. These libraries were subsequently sequenced on a Novaseq 6000 platform (Illumina) with cycles setting 28-10-10-90 cycles. Approximately, 25000 reads/cell and 5000 reads/cell were generated for respectively each of the GEX and CMO libraries. The raw data were demultiplexed using the CellRanger-7.0.1 mkfastq pipeline. The gene expression data were processed using the CellRanger-7.0.1 count pipeline. Gene expression data in combination with CMOs were processed using the CellRanger-7.0.1 multi pipeline. The mm10 reference genome was used for generating the counts matrix.

### scRNA-seq Analysis

scRNA-seq data were analyzed using the Seurat v5 R package (Satija et al. 2015; Butler et al. 2018; Hao et al. 2021, 2024; Stuart et al. 2019). In addition to CellRanger-7.0.1 multipipe line’s demultiplexing, a second demultiplexing step was performed using the HTODemux function. Seurat’s HTODemux function with a positive quantile = 0.99 threshold was used. Next, Median Absolute Deviation thresholding (Heumos et al. 2023; Germain et al. 2020) was used to remove lowquality cells with nFeature_RNA (the number of detected genes), nCount_RNA (the amount of detected RNA), and percent.mt (percentage of mitochondrial RNA) with thresholds 5 (initial threshold) and 3 (final threshold). Ambient RNA removal was performed on SoupX (Young and Behjati 2020) using default parameters. The scDblFinder package (Germain et al. 2021) was utilized to detect doublets, and 0.20 was set as the doublet threshold. After this step, the five libraries were merged and combined into one dataset with the JoinLayers function.

Before removing low-quality cells temporary mapping and clustering of the merged dataset was performed with the following functions of the Seurat: NormalizeData, ScaleData, FindVariableFeatures (vst, nFeatures=4000), RunPCA, FindNeighbors (dims=1:75), RunUMAP (dims=1:75, spread=2), FindClusters(resolution=4, algorithm=4, n.start=100, n.iter=100) and CellCycleScoring. Low-quality cells were removed using the following classifications: singlet-assigned cells from the Cell Ranger pipeline’s “Confidence Assignment Table”; singlet-assigned cells from HTODemux; singlets assigned cells from scDblFinder and a Median Absolute Deviation threshold of 3. Following this, cell numbers from clusters were compared before and after removing the low-quality cells, cell clusters containing mostly low-quality cells were also removed from the dataset. Markers of the cell types were detected using the FindMarkers/FindAllMarkers functions of Seurat.

Following the cleanup, the following steps were performed: cell cycle regression with ScaleData, FindVariableFeatures (nFeatures=4000), RunPCA, RunUMAP(dims =1:75, spread=1, n.neighbors=50, n.epoch=1000), FindNeighbors(dims=1:75, k.param=50), FindClusters(resolution=1.25, algorithm=4, n.start=100, n.iter=100). Clusters that require higher resolution were further divided using the FindSub-clusters function. Later, three levels of cell annotation were constructed: groups, clusters, and sub-clusters. The groups were then divided into three categories: “Early differentiation”, “Late Mesoderm” and “Hematopoietic.” Each of the three going through FindVariableFeatures, RunPCA(npcs=50), FindNeighbors(dims=1:50), and RunUMAP(dims=1:50). Markers of the annotated cells were detected with FindMarkers and FindAllMarkers function.

For the Differential Gene Expression Analysis (DGEA), each TF-KO was compared to the NT-Control with pseudo-bulked DESeq2 (Love et al. 2014) in all cell type annotations in a three-level (groups, clusters, and subclusters) resolution. Single-cell expression counts were converted to a pseudo-bulk counts matrix with Seurat’s “AggregateExpression” function. Then, a DESeq2 object was created using the “DESeqDataSetFromMatrix,” a function of DESeq2. Genes with low read counts were removed. The “DESeq” function is applied without extra arguments, and padj < 0.05 is used to detect the Differentially Expressed Genes (DEGs).

The marker detection of the three small sub-clusters (EndRsad2, EndCyp26b1, baIfit1) performed with Seurat’s FindMarkers function was utilized with the Wilcox method with min. pct and log.fc.threshold equal to 0.

Gene Set Enrichment Analysis (GSEA) was performed with the fgsea R package (Korotkevich et al. 2021) on MsigDB’s (Subramanian et al. 2005; Mootha et al. 2003) Hallmark, GO-Biological Processes, and Reactome gene sets. Results of the DGEA with “stat” values were used to rank the gene list. Low-expression genes were removed if a gene had a base value of less than 15. The fgsea function was run with minSize = 15 and maxSize = 500 settings. The top 20 results of each gene set were used for visualization. Results that showed pad < 0.001 were manually curated into categories (i.e., Metabolism, Signal).

Speckle (Phipson et al. 2022) and miloR (Dann et al. 2022) packages were used for the differential abundance analysis (DAA) (or Compositional Analysis). For speckle, default settings were used with the propeller function.

For MiloR, the following process was repeated for each KO-TF compared to Control in EarlyDiff, Late-Mesoderm, and Hematopoietic groups. KNN graphs were constructed with “buildGraph” (parameters: k=35, d=50) and” makeNhoods “(prop = 0.3, k=35, d=50, refined=TRUE). The “k” value was calculated with the 5 x (number of samples) formula. Euclidian distances were calculated with” calcNhoodDistance” (parameters: d=50). Differential neighborhood abundance was tested with” testNHoods.” The dependency between the average number of cells per sample and the logFC was checked using the “plotNhoodMA” function with MAPlot. With “buildNhoodGraph,” an abstract graph of the neighborhoods visualization was created and visualized with “plotNhoodGraphDA” (parameters= alpha (SpatialFDR) = 0.1). For annotation of the neighborhoods, the “annotateNhoods” function was used to annotate the neighborhoods with the sub-clusters. In case the neighborhoods did not contain at least 50 percent of the majority from any sub-clusters, it is labeled as “Mixed.” “plotDAbeeswarm” was used for visualization for the distribution of log fold changes across neighborhood annotations. “findNhoodGroupMarkers” was used for the marker gene detection on selected neighborhoods (SpatialFDR <0.1).

The *in vivo* scRNA-seq mouse gastrulation atlas (Pijuan-Sala et al. 2019) and extended atlas (Imaz-Rosshandler et al. 2023) were used for visualization, aligning *in vitro* and *in vivo* data through the FeaturePlot function of Seurat. Pseudotime construction was performed with Palantir (Setty et al. 2019).

### DEG, Literature, and ChIP-seq Integration

Public human and mouse datasets were used to compare the DEGs list with each TF KO based on results from the literature and DEGs’ relationship to binding sites of the KO-TFs.

To establish the binding sites of Atf3, the following datasets were used from the ENCODE portal (Hitz et al. 2023; Luo et al. 2019; Dunham et al. 2012) (https://www.encodeproject.org/) with the following identifiers: ENCAN506YWT, ENCSR480LIS, ENCSR879YNX, ENCSR000BUG, ENCSR000BKC, ENCSR000BJY, ENCSR402ZCY, ENCSR000BKE, ENCSR568ZXG, ENCSR632DCH, ENCSR000BNU, ENCSR000DOG, ENCSR028UIU (Supplemental Table 8A). Peaks of these data sets were uploaded to the Galaxy web platform (https://usegalaxy.org/)(Community et al. 2024) and were combined (~210.700 peaks) and then merged with the mergeBED function of the bedtools (~97K peaks) (Quinlan and Hall 2010). These coordinates were converted from human hg38 to mouse mm10 using UCSC Genome Browser-LiftOver (Hinrichs et al. 2006) with default settings for conversion across the species. Additionally, Zhang and co-workers (Zhang et al. 2023) ChIP-seq data in a mouse with mm9, which were converted to mm10 with LiftOver using default parameters for the same species conversion. Later, all converted data were combined and merged, resulting in 65307 peaks in mm10. As a next step, Atf3 peaks in mm10 were intersected with ATAC-seq data (manuscript in preparation), which covers daily chromatin accessible data from day 3 to day 7 in whole EB with the bedtools Intersect interval function with a minimum overlap fraction of 0.1, which resulted in 28136 peaks.

Furthermore, these peaks were checked whether they contain Atf3 motifs or not with the intersection of the following process: human and mouse motifs were selected from Jaspar (Castro-Mondragon et al. 2021; Fornes et al. 2019; Khan et al. 2017; Khan and Mathelier 2017) (MA605.1, MA0605.2, MA0605.3, MA1988.1, MA1988.2, FIMO (Grant et al. 2011) used for each motif on ATAC-seq peaks with an output threshold of 0.01, including reverse complement strand parameters. The resulting motifs were merged later. This merging resulted in 27943 peaks in mm10, which were used to link them to the nearest gene with the GREAT web tool (http://great.stanford.edu/public/html/) (McLean et al. 2010; Tanigawa et al. 2022), which has a single nearest gene option within a 1000kb distance. The output of this process was then used to overlap it with the DEG list of Atf3-KO. Also, the ChIPseeker package (Yu et al. 2015; Wang et al. 2022) was used for the peak distance to the TSS and peaks positions genomics features characterization with hg38 and mm10 Ensembl genomic annotations (Harrison et al. 2023).

Harmonize 3.0 from Rouillard A et al. 2016, ENCODE Transcription Factor Targets (Rouillard et al. 2016) the database used for ATF3 targets. The mouse orthologs are converted from human with syngoportal-version-12-12-2020 (Koopmans et al. 2019) and intersected with the DEGs list of the Atf3-KO.

Furthermore, five studies of the knockout and knockdown ATF3 in humans and mice later intersected with the DEGs list (Xu et al. 2011; Sood et al. 2017; Marcantonio et al. 2021; Badu et al. 2024; Labzin et al. 2015).

A similar process was followed for Zfp711, which is described for Atf3. ChIP-seq data were used from GSE102616, GSE20673, GSE145160 (Rhie et al. 2018; Kleine-Kohlbrecher et al. 2010; Ni et al. 2020). The motif of the ZNF711 has been described (Ni et al. 2020). For ZNF711 literature DEGs, the following GEO accession codes/studies were used GSE102616, GSE145160, and Snabel and co-workers (Snabel et al. 2024; Ni et al. 2020; Rhie et al. 2018).

## DATA AVAILABILITY

scRNA-Seq data generated in this study have been deposited in the European Nucleotide Archive (ENA) database under the accession number PRJEB82953. ATAC-seq data used in this study is a separate project and will be deposited in the ENA database upon completion of the manuscript preparation. UCSC browser sessions can be accessible with the following links https://genome.ucsc.edu/s/mdrcetin/hg38_ATF3, https://genome.ucsc.edu/s/mdrcetin/mm10_Atf3, https://genome.ucsc.edu/s/mdrcetin/mm10_Zfp711, https://genome.ucsc.edu/s/mdrcetin/hg19_ZNF711, and https://genome.ucsc.edu/s/mdrcetin/hg38_ZNF711 The data supporting the findings of this study are available from the corresponding author upon reasonable request.

## CODE AVAILABILITY

The code that was used in this study is available at https://github.com/mdrcetin/Cmo_Atf3_Zfp711_Bcl6b and https://ridvan-cetin.github.io/CMO_Atf3_Zfp711_Bcl6b/. Our scRNAseq can be explored with shinnyapp-link at https://ridvan-cetin.github.io/CMO_Atf3_Zfp711_Bcl6b/.

## COMPETING INTEREST STATEMENT

The authors declare no competing interests.

### ACKNOWLEDGMENTS

We are grateful to Alex Maas and Tsung Wai Kan for their assistance with the FACS, Cengizhan Acikel for statistical advice, and Dilek Akyol for financial support.

## AUTHOR CONTRIBUTIONS

R.C. and F.G. conceived the project. R.C. conducted the experiments and performed the data analysis. G.P., J.v.S., E.B., and Y.F. assisted in conducting the experiments. R.H., G.v.B., M.S. and A.K. assisted in the data analysis. R.C. and F.G. wrote the manuscript; all authors, including J.G., D.H., J.v.H. and E.M., contributed to the writing and provided feedback.

---

**Supplemental Figure S1.**
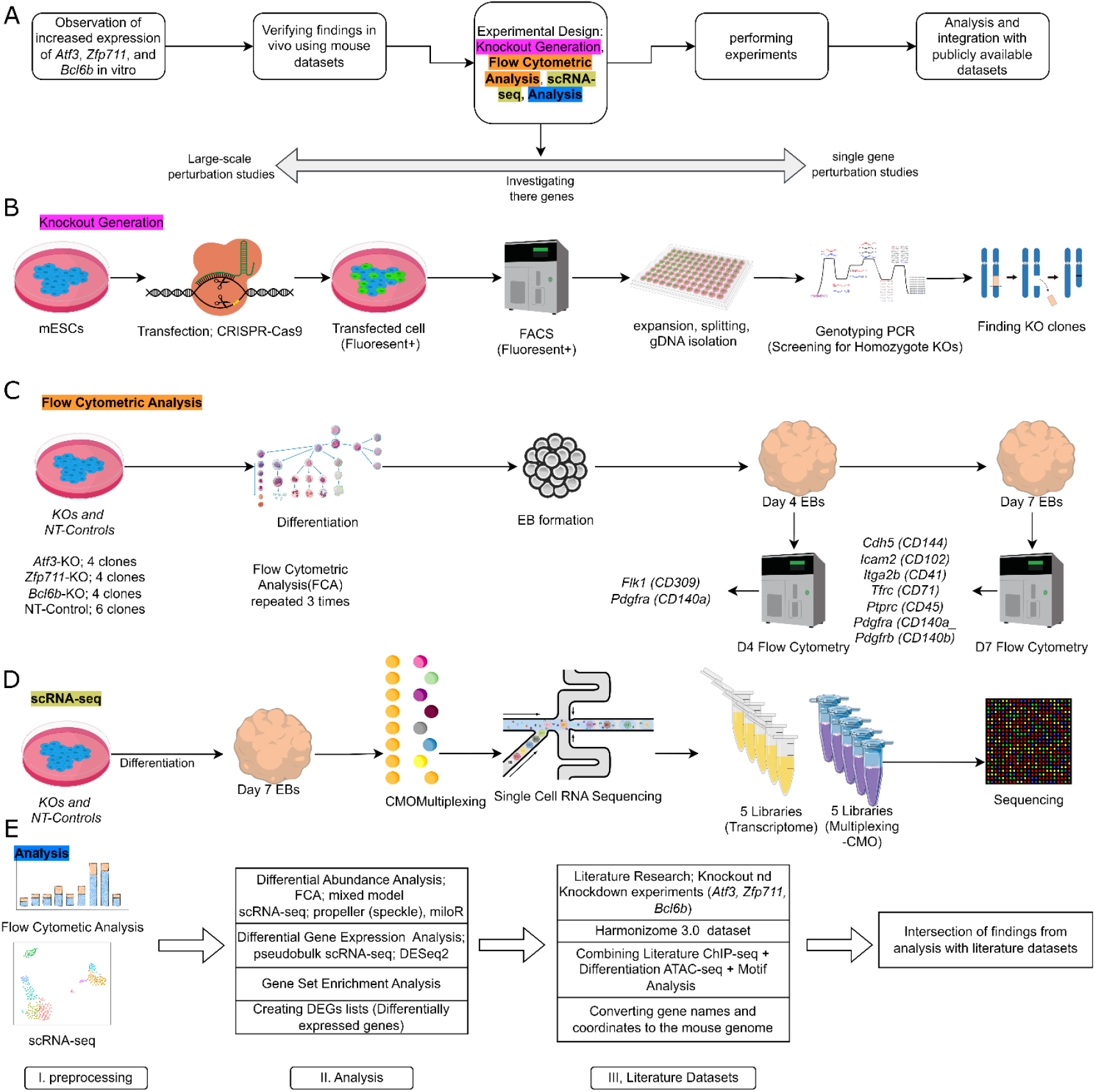
Steps of the study, and details of the individual steps. **Panel A** shows the overall timeline of the study. **Panel B** shows the knockout cell line generation steps. **Panel C shows** the Flow Cytometric Analysis workflow. **Panel D** shows the multiplexed scRNA-seq workflow. **Panel E** shows the analysis workflow of the FCA and scRNA-seq datasets.

**Supplemental Figure S2.**
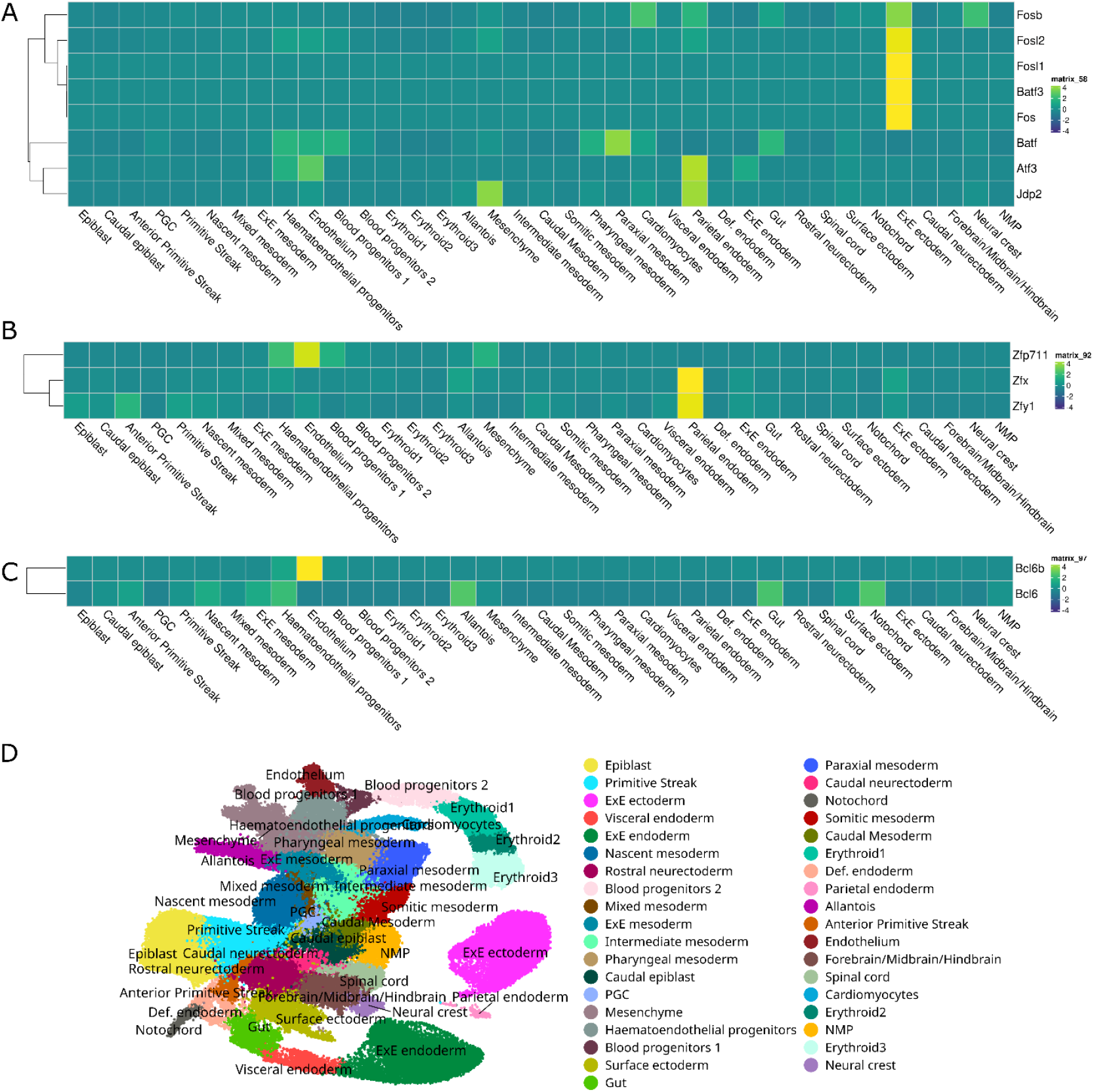
Selected TFs and the expression of their paralog(s) *in vivo*. **Panel A** shows the expression of *Atf3* and its paralogs, as documented in the mouse gastrulation atlas. **Panel B** shows the expression of *Zfp711* and its paralogs (*Zfy2* has been removed since no expression could be detected). **Panel C** shows the expression of *Bcl6b* and its paralog. **Panel D** shows the annotation of the cell types in UMAP from the mouse gastrulation atlas (Pijuan-Sala et al. 2019).

**Supplemental Figure S3.**
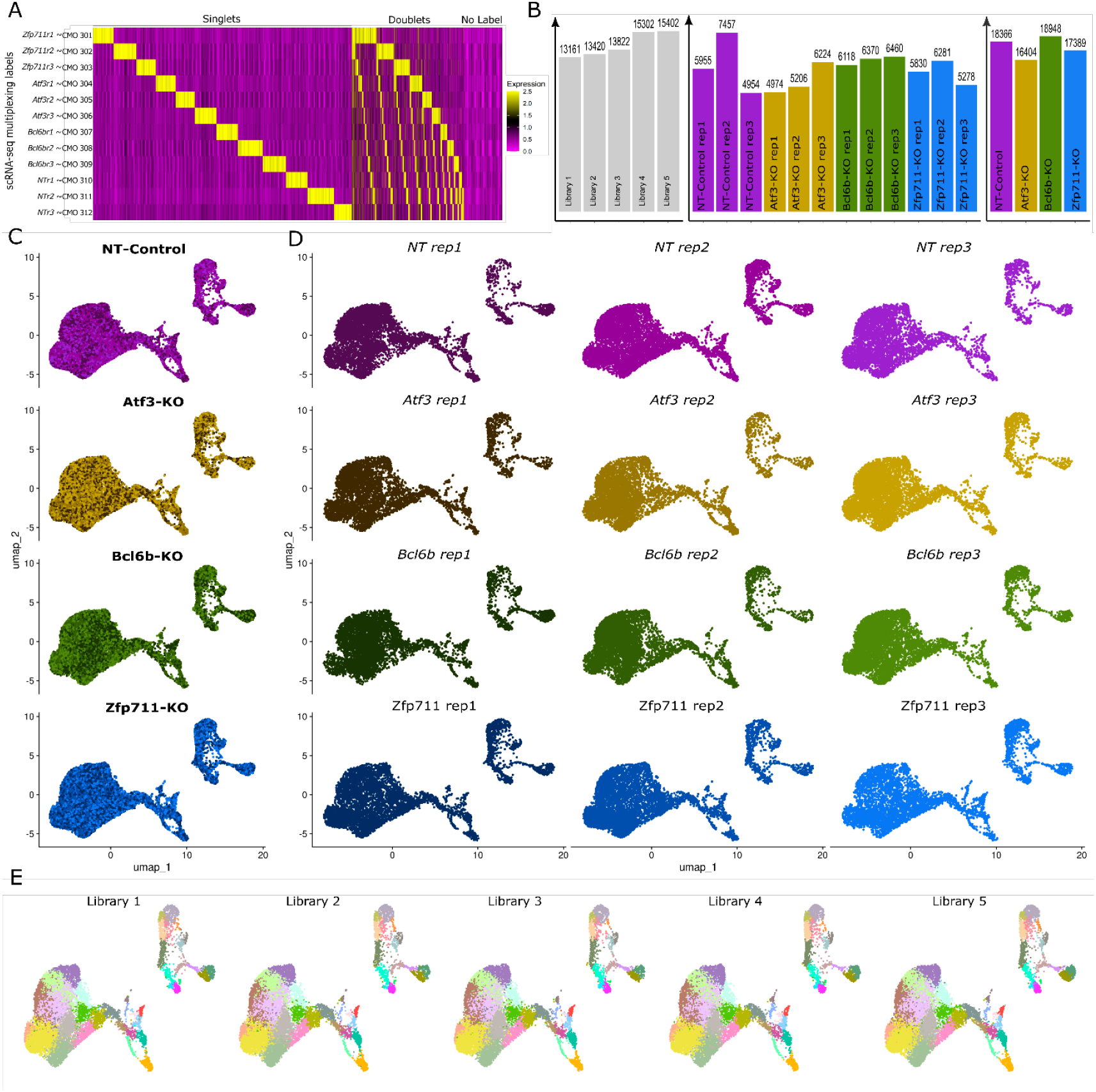
Demultiplexing scRNA-seq and cell distribution. **A**. An example of demultiplexing, using the Seurat-HTODemux function for a library. The Y-axis shows the samples and CMO labels. The X-axis shows individual cells. The heatmap shows CMO label counts and categorizes cells into singlets, doublets, and negative/No Signal/unassigned. **B**. Distribution of number of cells after quality control. The panel on the left shows the number of cells in each library. The panel in the middle shows the number of cells from each sample/clone. The panel on the right shows the number of cells for each experimental condition. **C**. UMAPs are split by conditions. **D**. UMAPs are split by samples. **E**. UMAPs are split by lanes. Different colors represent different sub-clusters (see Figure 2C).

**Supplemental Figure S4.**
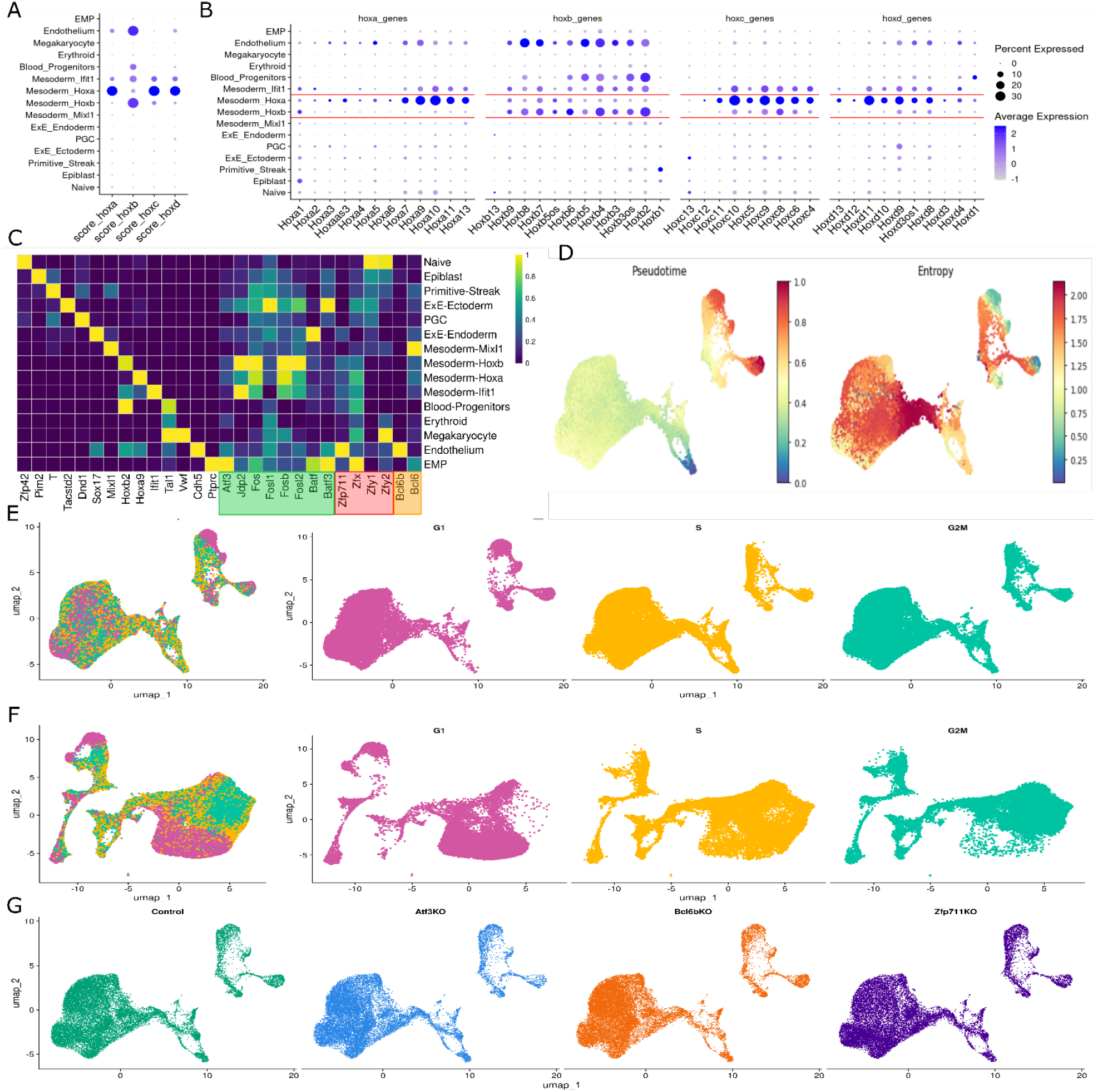
Marker genes for clusters, pseudotime, and cell cycle. **A**. Expression score of *Hoxa, Hoxb, Hoxc*, and *Hoxd* genes in clusters (y-axis). **B**. Details of the individual *Hoxa, Hoxb, Hoxc*, and *Hoxd* genes expression in clusters (y-axis). *Hoxa*^+^ Mesoderm and *Hoxb*^+^ Mesoderm are highlighted with red lines. **C**. The expression at the cluster level cell annotation of known marker genes, *Atf3* and some of its paralogs (green), *Zfp711* and its paralogs (red), and *Bcl6b* and its paralog (orange). **D**. UMAP mapping of the pseudotime from early (blue) to late (red) and entropy (differentiation potential). **E**. Cell cycle phases present in cell cycle-regressed data. **F**. Cell cycle phases present in non-regressed data for the cell cycle. **G**. UMAPs are split by conditions.

**Supplemental Figure S5.**
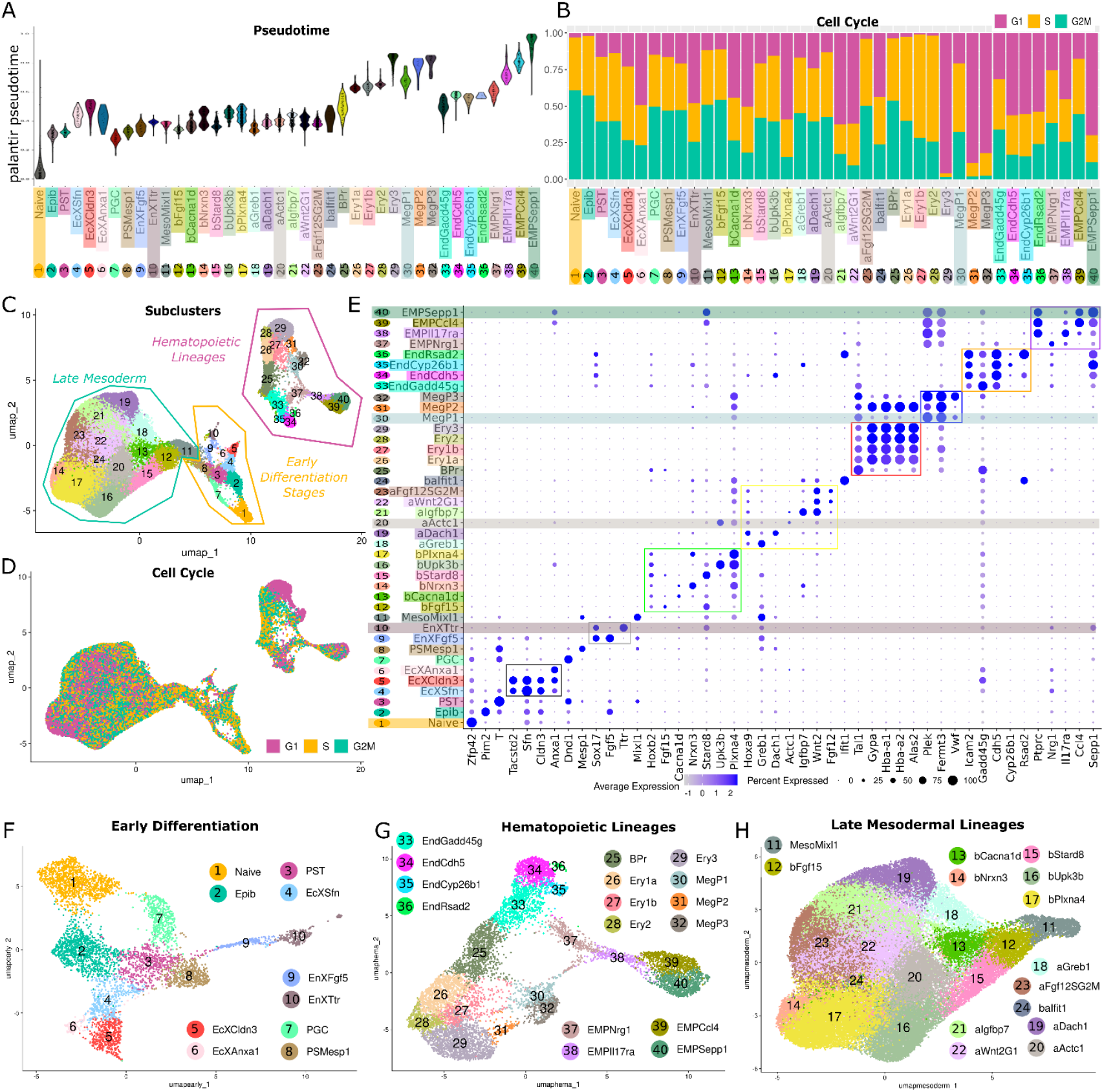
Details of the data used for sub-cluster level annotation with numeric labels. **A**. Pseudotime trajectory beginning from the naïve sub-cluster to later stages of the differentiation. **B**. Distribution of the cell cycle phases per sub-cluster. **C**. UMAP shows sub-cluster level annotation (numbers) and their group-level annotation (lines). **D**. Mapping of the cell cycle distribution on UMAP. **E**. Marker genes are used for sub-clustering and naming the sub-cluster-level cell type annotation (colored boxes and highlights are used as guidance). **F-G**. scRNA-seq data was divided into three subsets based on the groups, and new UMAPs were created for each.

**Supplemental Figure S6.**
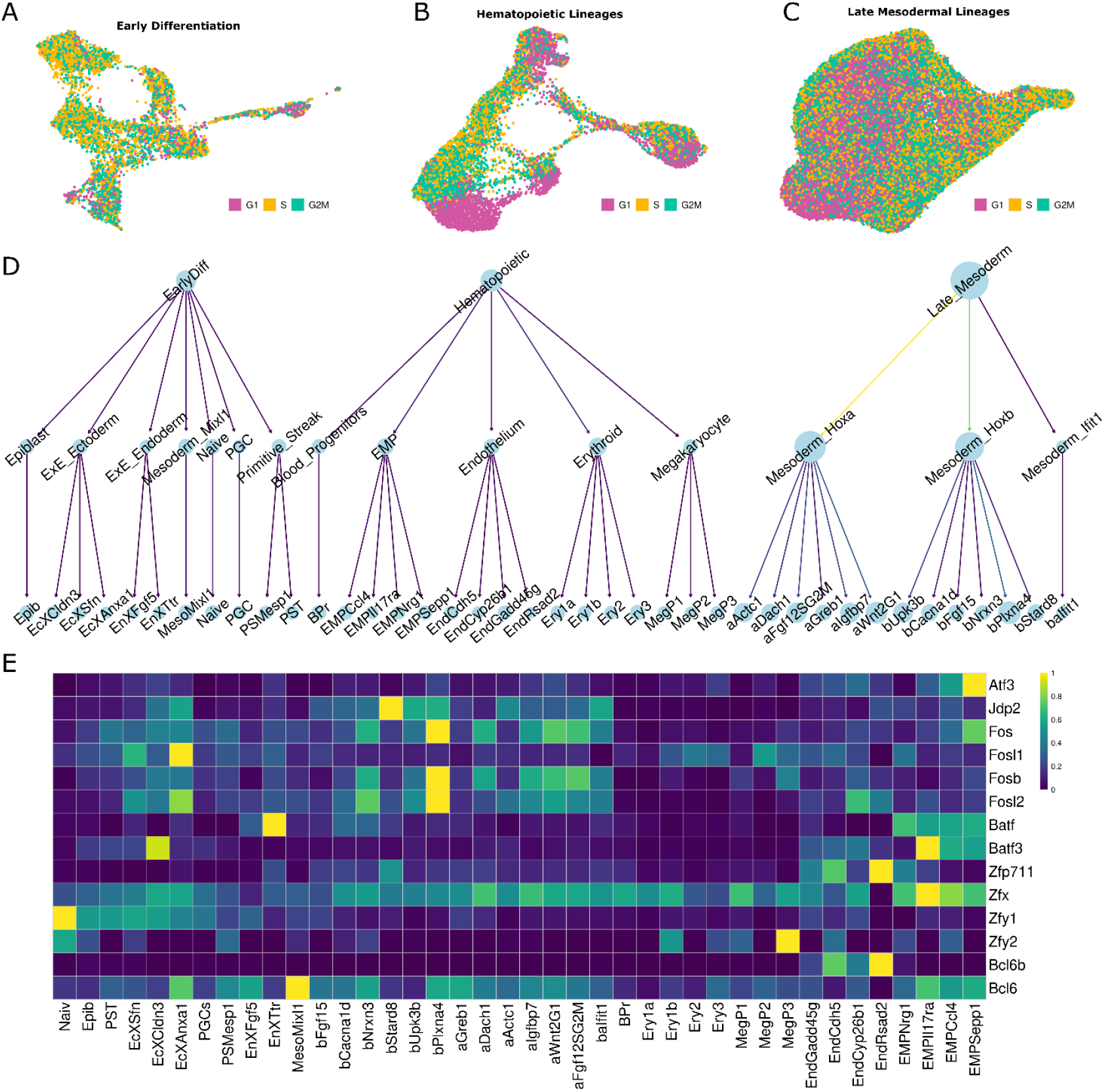
Mapping the cell cycle score on group-divided UMAPs and cell type annotation hierarchy. **A-C**. Cell cycle scores are mapped on group-divided UMAPs. **D**. Cell type annotation hierarchy with three groups, 15 clusters, and 40 sub-clusters (the size of the circles indicates the number of cells). **E**. Expression of *Atf3, Zfp711, Bcl6b*, and their paralogs at sub-cluster level.

**Supplemental Figure S7.**
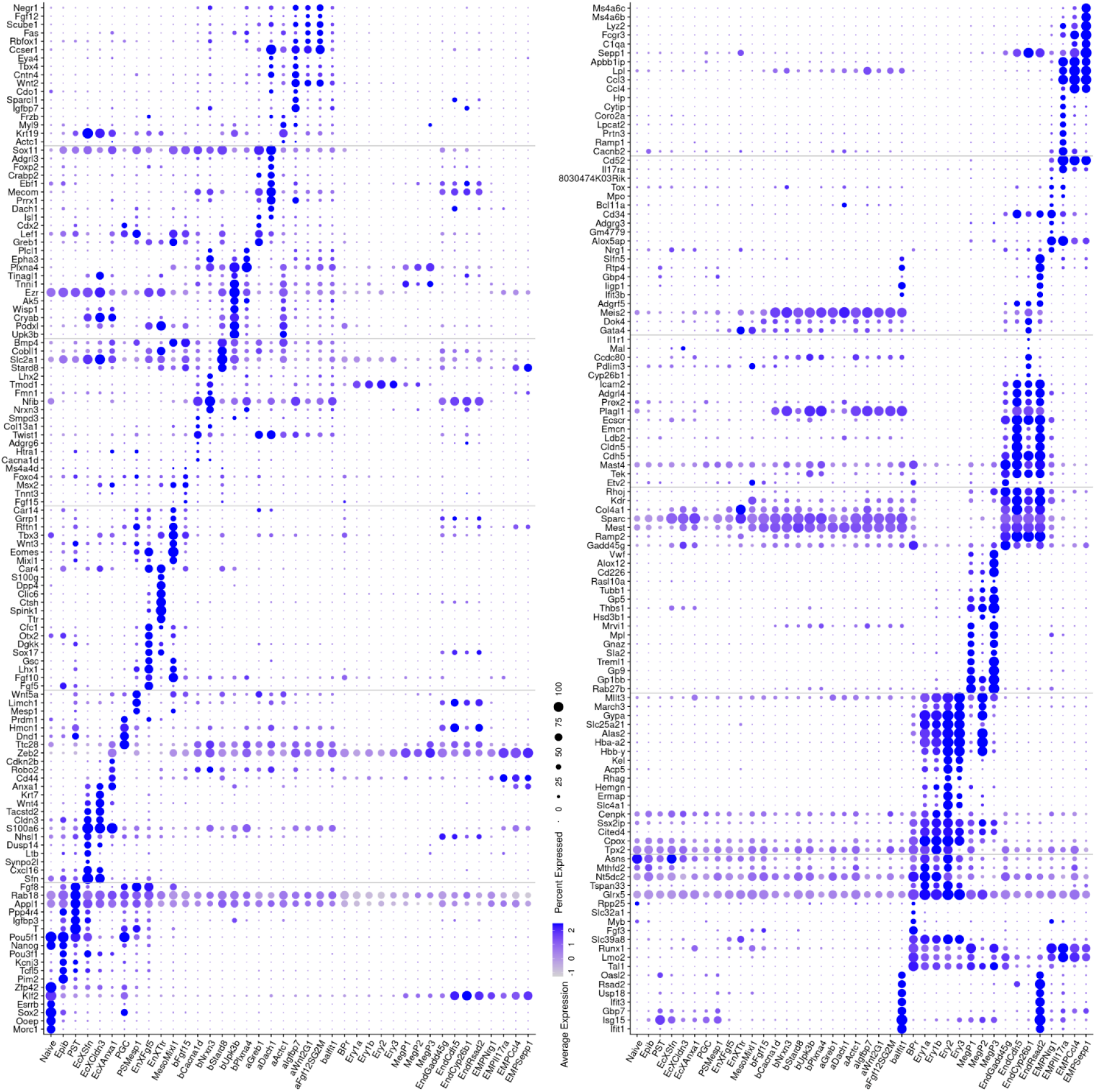
Extended list of markers for sub-cluster-level annotation.

**Supplemental Figure S8.**
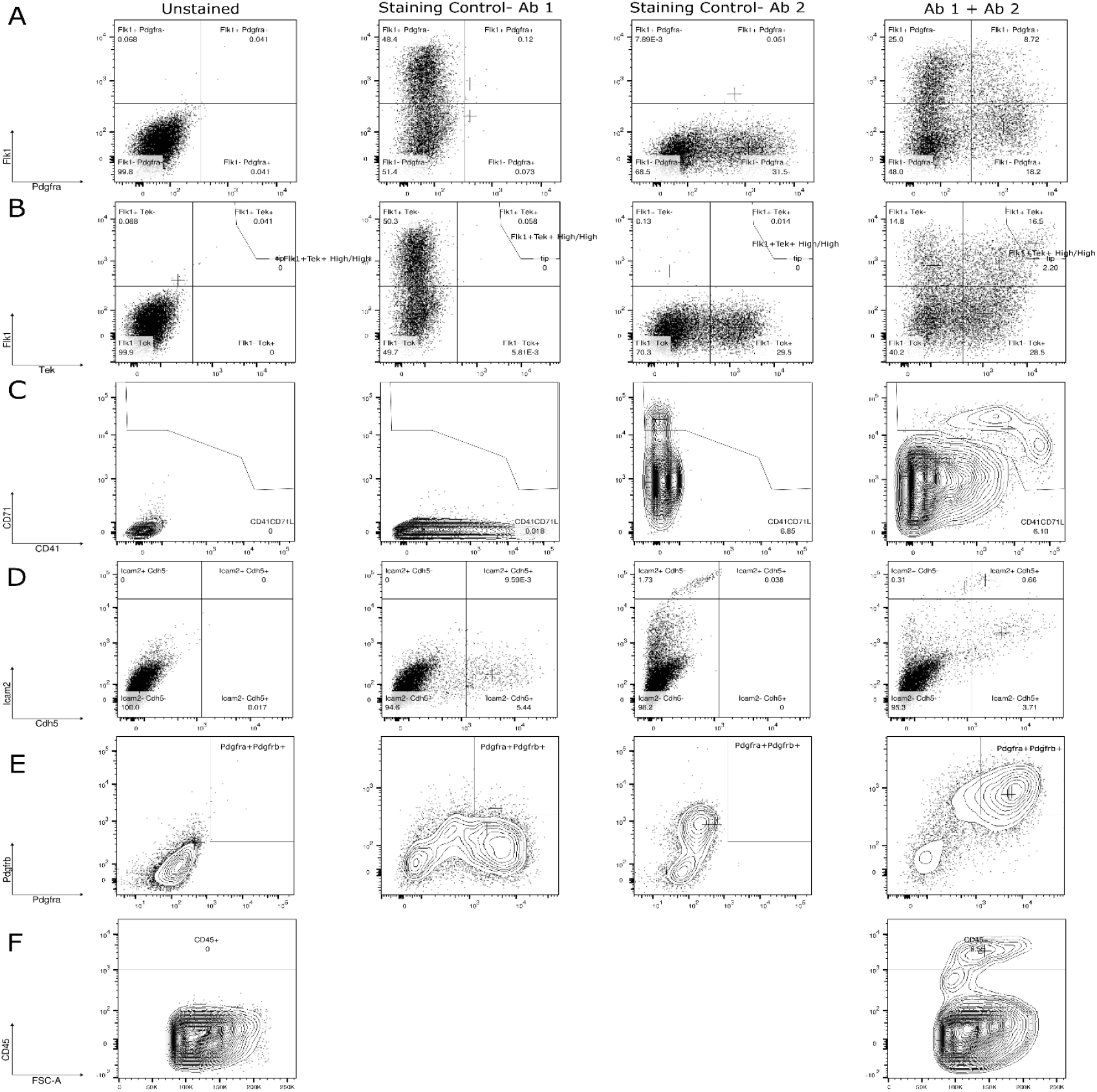
Staining for cell surface markers and example gating. **A**. Flk1/Pdgfra staining on day (D) 4. **B**. Flk1/Tek staining on D4. **C**.CD41/CD71 staining on D7. **D**. Icam2/Cdh5 on D7. **E**. Pdgfra/Pdgfrb on D7. **F**. CD45 staining on D7.

**Supplemental Figure S9.**
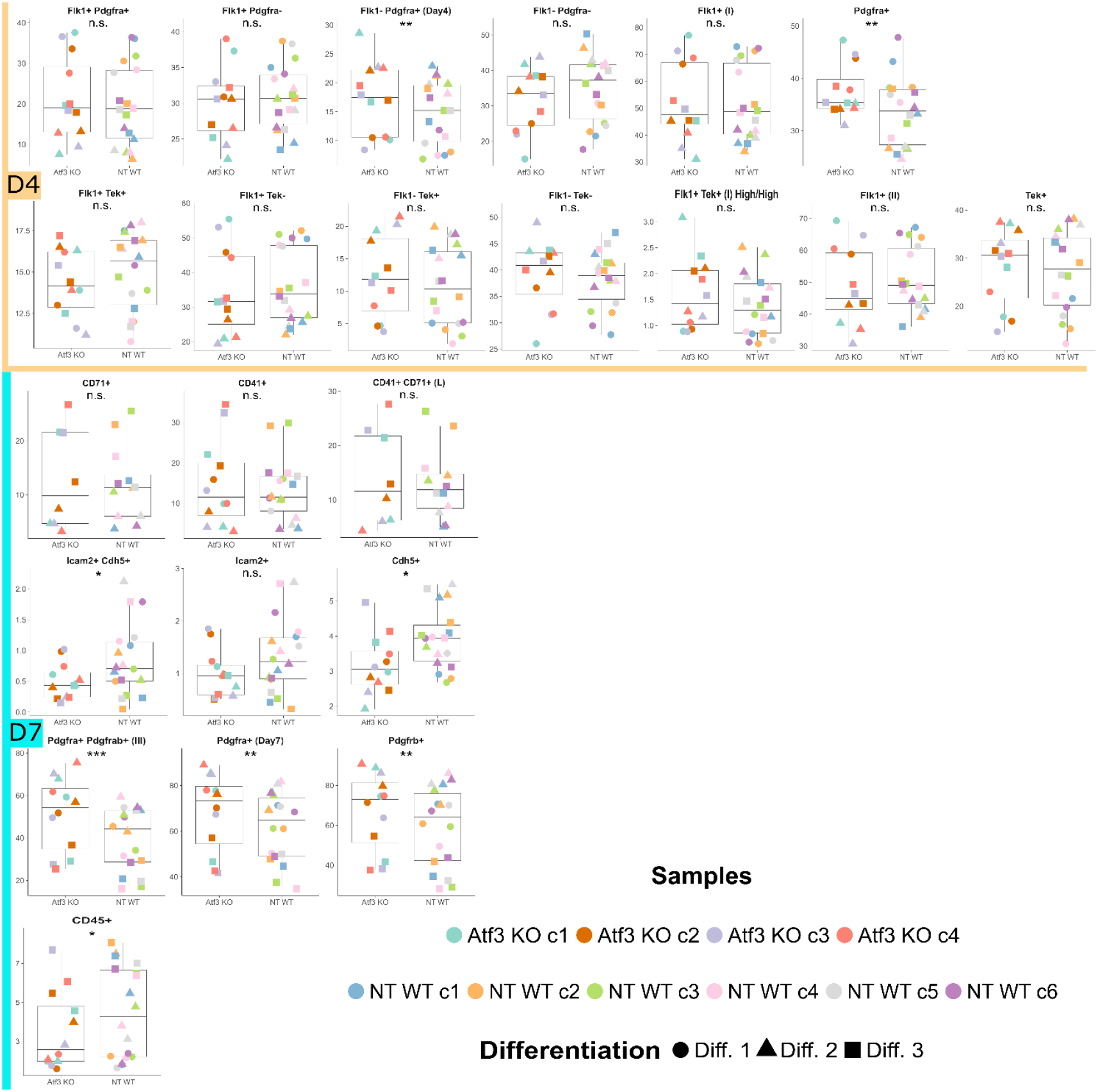
Flow Cytometric Analysis results of the *Atf3*-KO versus NT-Control cells (at D4 (orange) and D7 (turquoise)). Details of the dataset and statistical analysis can be accessed in Supplemental Table S1A,B. (n.s.) not significant: P > 0.05, (*) P < 0.05, (**) P < 0.01, (***) P < 0.001.

**Supplemental Figure S10.**
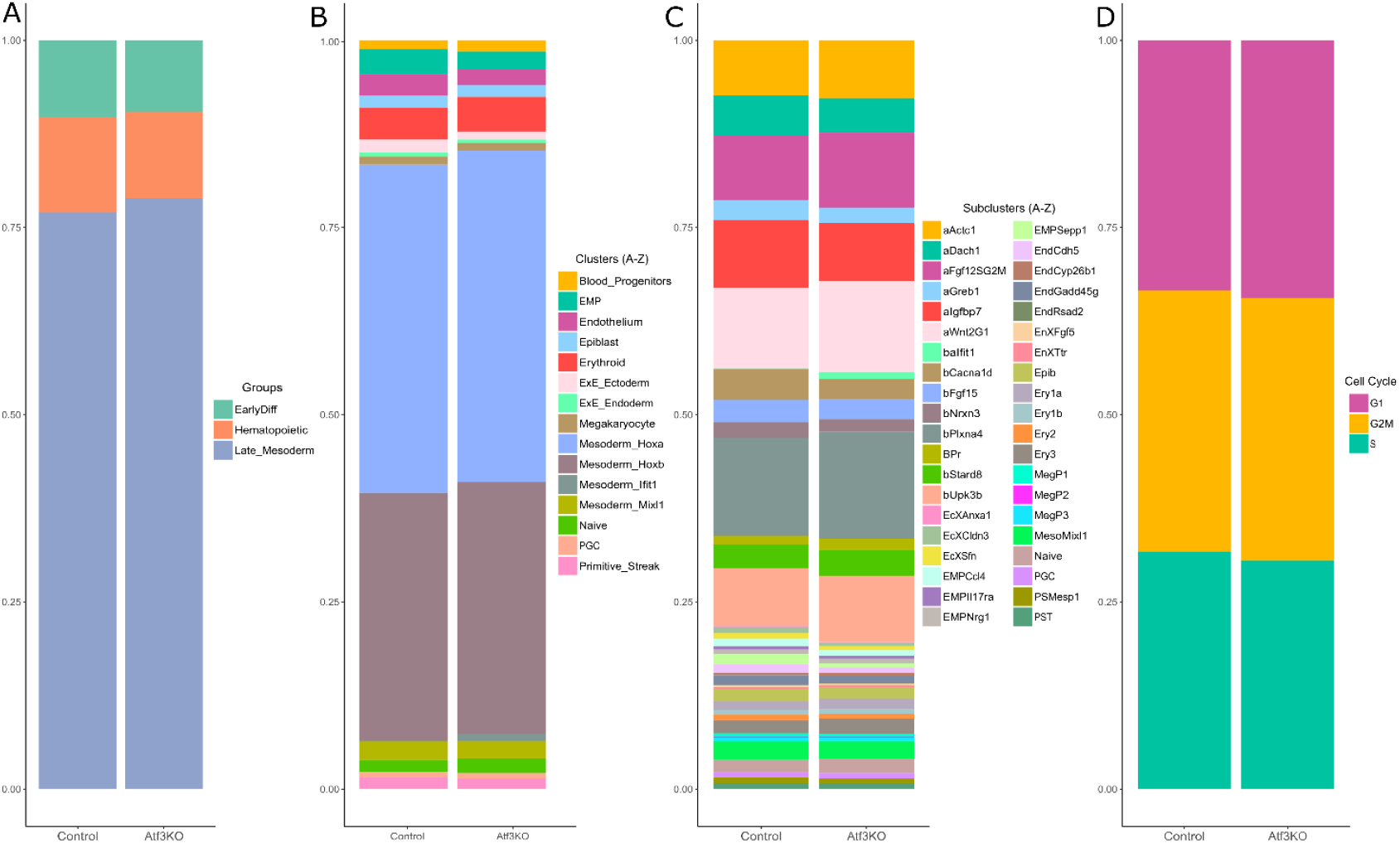
*Atf3*-KO cells: Differential Abundance Analysis results using Speckle R Package. **A-D**, Fraction of annotated cell types in groups (**A**), clusters (**B**), sub-clusters (**C**), and cell cycle (**D**).

**Supplemental Figure S11.**
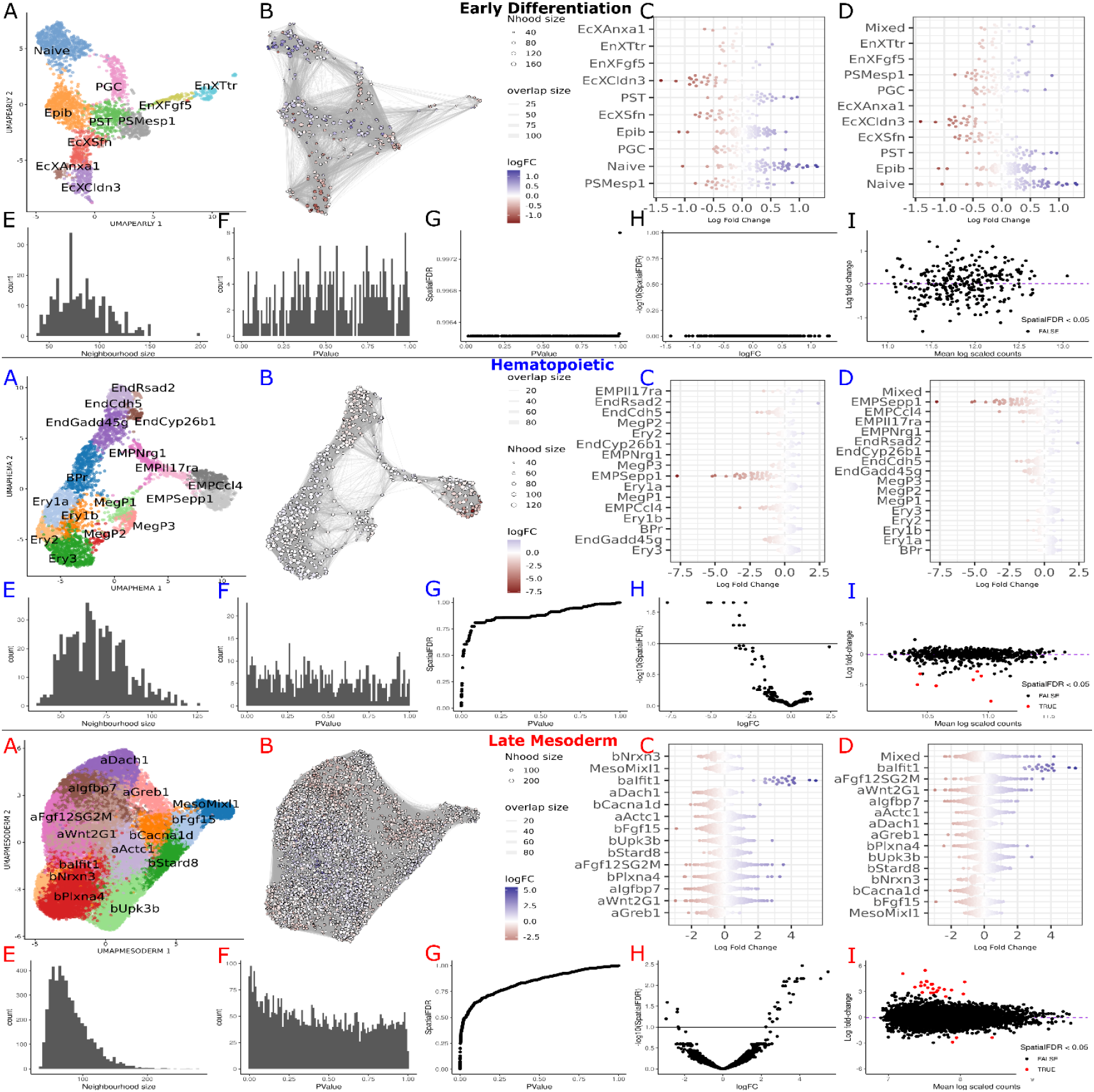
Results of the *Atf3*-KO cells Differential Abundance Analysis by miloR, in the groups Early Differentiation (Black), Hematopoietic (Blue), and Late Mesoderm (Red). **A**. UMAP with cell type annotation. **B**. miloR neighborhoods visualization on UMAP coordinates. **C**. Neighborhoods categorization to sub-clusters. **D**. Neighborhoods categorization into sub-clusters (50% threshold). **E**. Distribution of the size of the neighborhood. **F**. P-value histogram of the neighborhoods. **G**. P-value and Spatial FDR relationship for neighborhoods. **H**. Fold change and Spatial FDR relationship. **I**. Mean log-scaled counts and log fold-change relation of the neighborhoods. The expectation is that the neighborhoods will be located around log fold-change 0. This figure is Supplemental to Figure 4K-M, and the dataset that outputs analysis can be seen in Supplemental Table S3A-C.

**Supplemental Figure S12.**
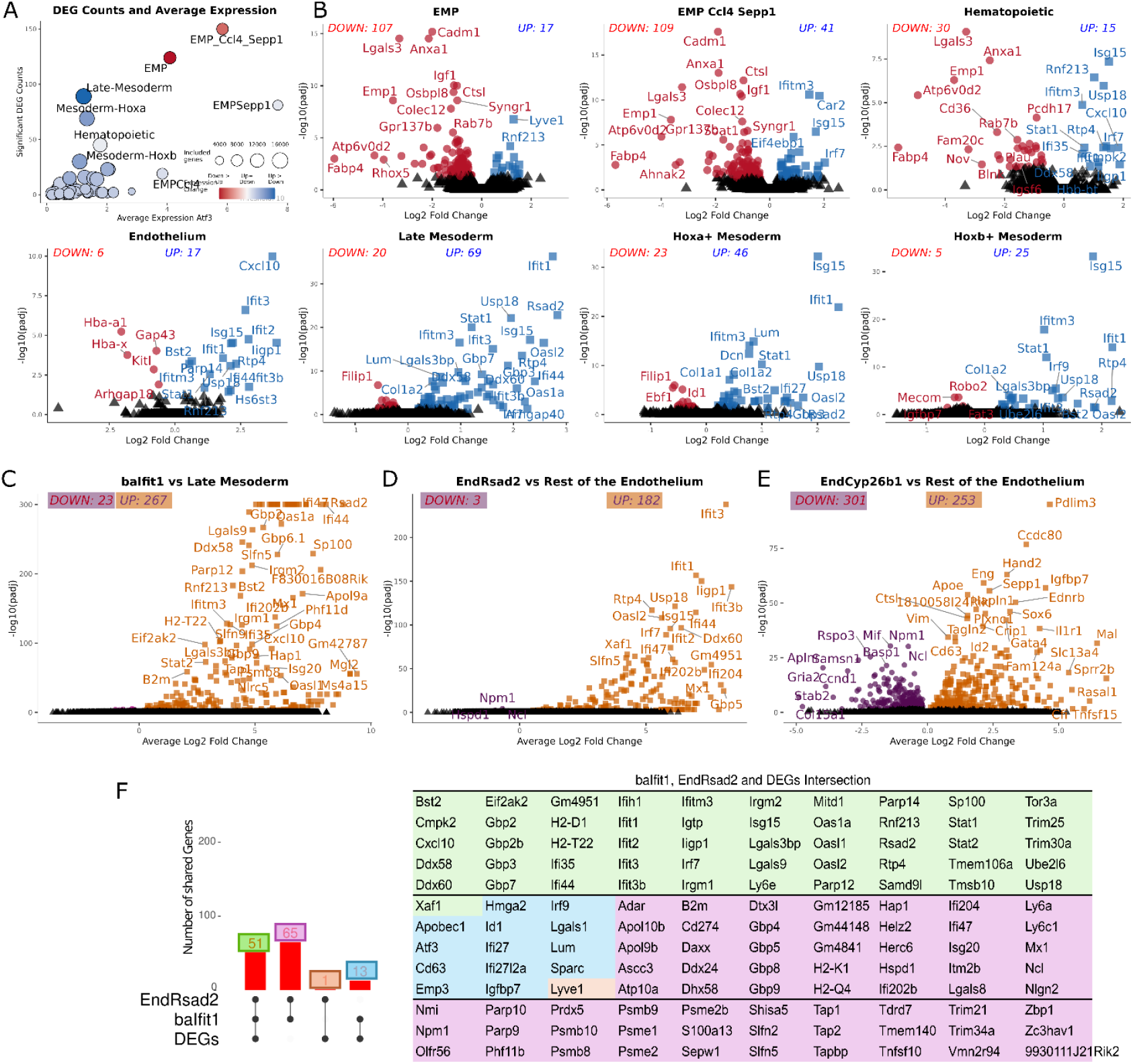
Atf3-KO cells: Differential Gene Expression Analysis and Unique Cluster Markers. **Panel A** shows the average expression level and number of the significant DEGs (padj < 0.05) in groups, clusters, and sub-clusters. The size of the circles indicates the number of genes included in the analysis after removing low-expressed genes in the group, cluster, or sub-cluster. The circle’s color indicates up- or downregulation (*blue* and *red* respectively). **Panel B** shows the up- or downregulation of genes in multiple conditions as a volcano plot. **Panels C-E**. Marker genes of the small/unique sub-clusters are baIfit1, EndRsad2, and EndCyp26b1. **Panel F**. The number of shared genes between EndRsad2, baIfit1, and DEGs with color code. Supporting data are shown in Supplemental Table S4A-K.

**Supplemental Figure S13.**
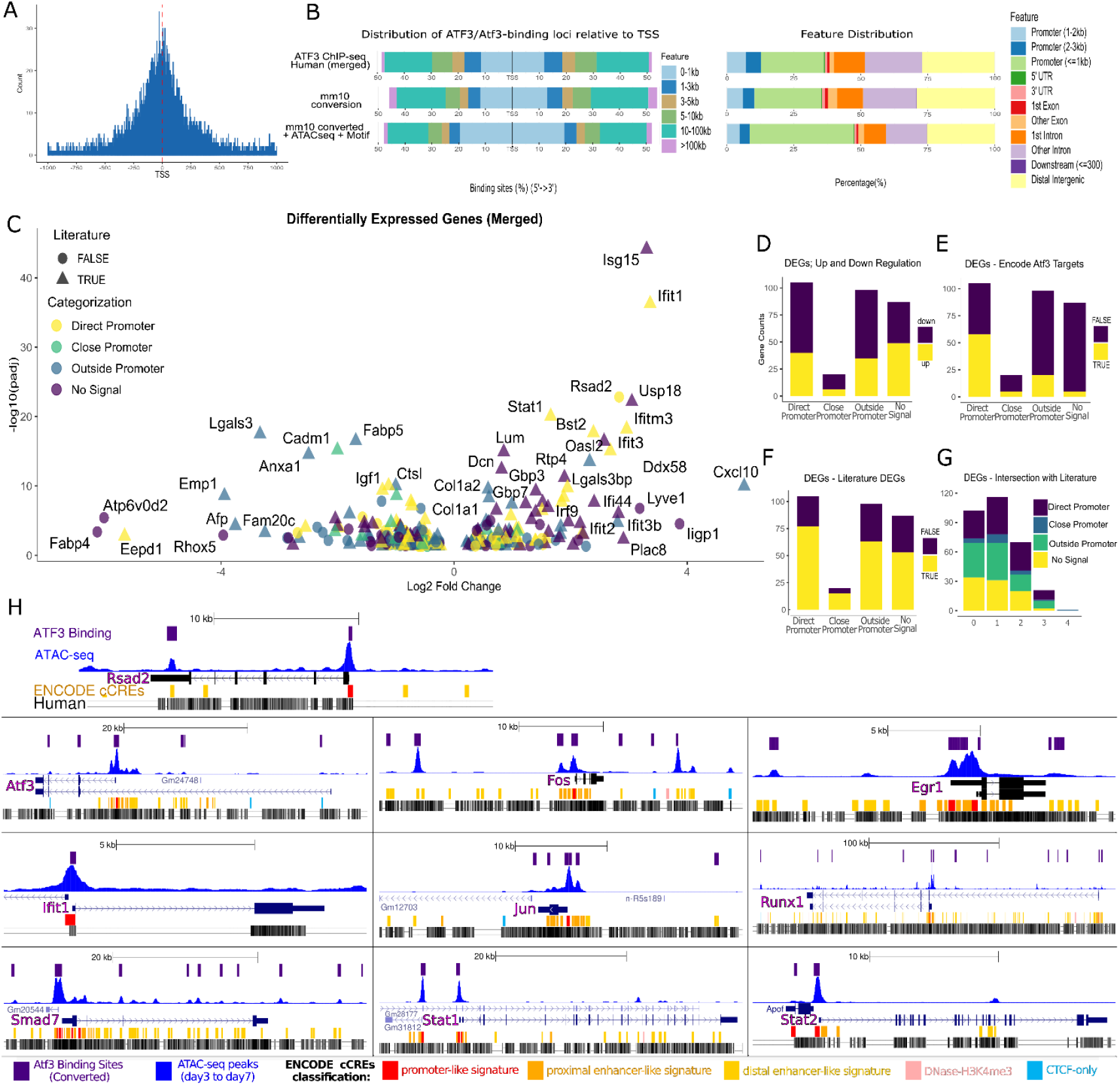
Results and categorization of the Differential Gene Expression Analysis of the *Atf3*-KO cells. **A**. Atf3 binding regions distribution histogram relative to the Transcription Start Site (TSS). **B**. Atf3/ATF3 ChIP-seq peak regions location and distance to TSS in three stages of the analysis going from human to mouse. **C**. Volcano plot of all the significant DEGs. **D-G**, Distribution of the DEGs, Atf3-binding categorization, TF Targets dataset, and DEGs found in the literature. **H**. Some examples of Atf3 binding sites with ATAC-seq peaks in three example DEGs. This figure is Supplemental to Figure 5E-H. Supporting data can be seen in Supplemental Table S4A-K, Supplemental Table S6A, and Supplemental Table S7A-E.

**Supplemental Figure S14.**
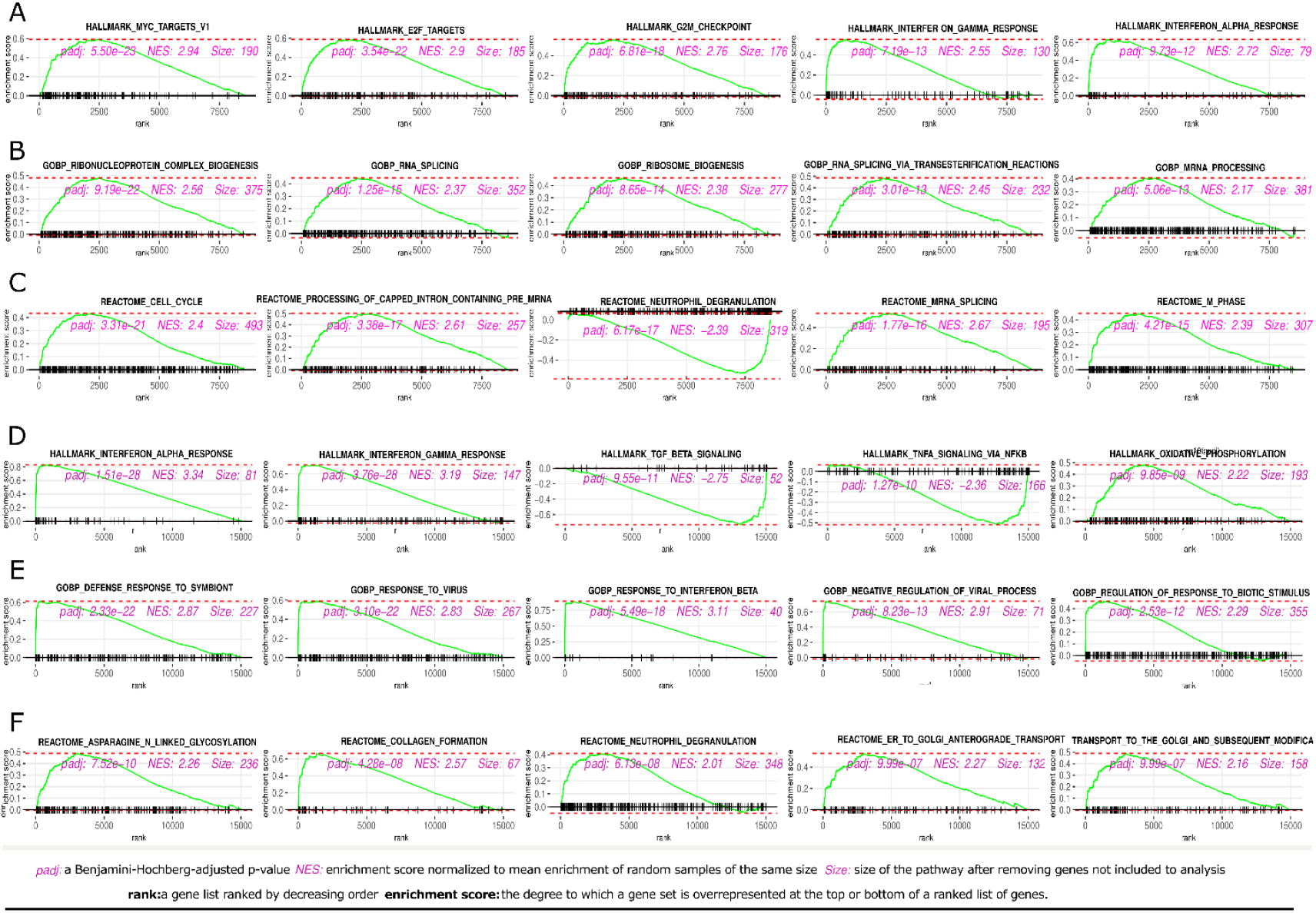
GSEA Results of the *Atf3*-KO cells on EMPs and Late Mesoderm. **Panels A-C** show enrichment plots of the top-5 results on EMPs. **Panels D-F** show for Late Mesoderm the GSEA results, similar to panels A-C. This figure is Supplemental to main Figure 6, and supporting data can be seen in Supplemental Table S5A-F.

**Supplemental Figure S15.**
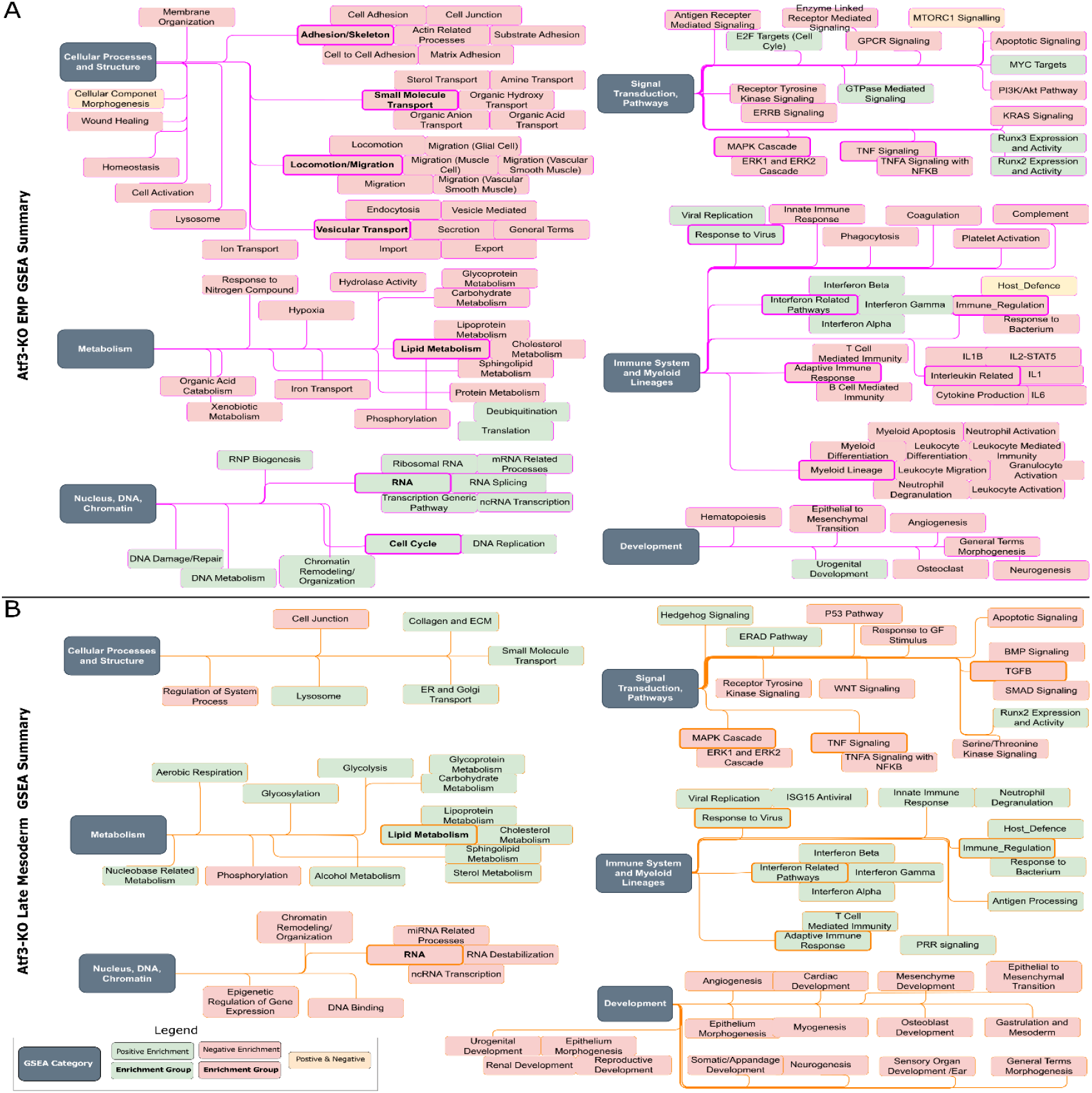
Manually grouped GSEA results of the *Atf3*-KO cells. **Panel A** shows the categorization of EMPs GSEA results. **Panel B** shows the categorization of Late Mesoderm GSEA results. This figure is Supplemental to Figure 6, and supporting data can be seen in Supplemental Table S5A-G.

**Supplemental Figure S16.**
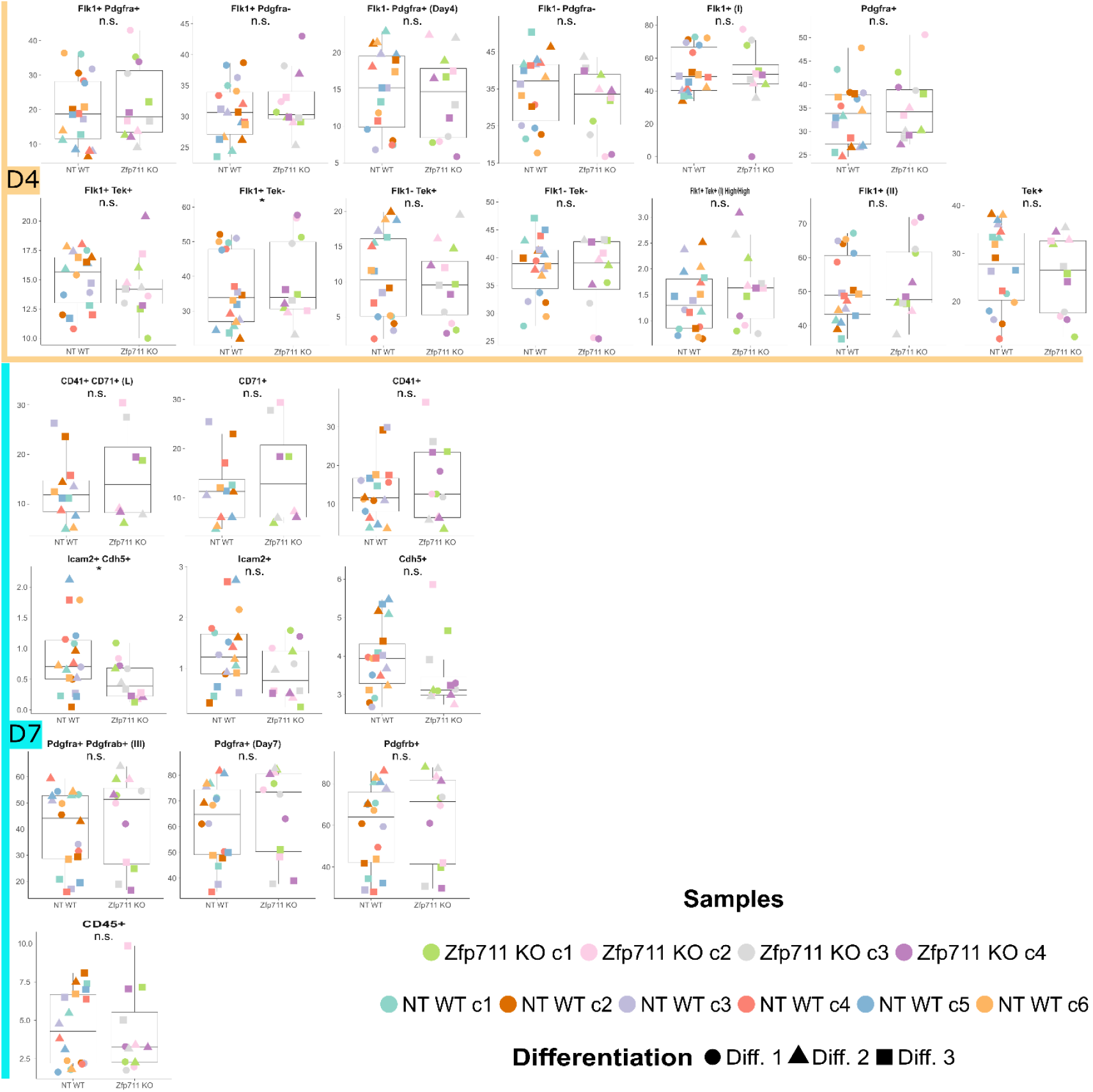
Flow Cytometric Analysis Results of the *Zfp711*-KO versus NT-Control cells (D4 (orange) and D7 (turquoise)). Supplemental to Figure 7B-F. Details of the dataset and statistical analysis can be accessed in Supplemental Table S1C-D.(n.s.) not significant: P > 0.05, (*) P < 0.05, (**) P < 0.01, (***) P < 0.001.

**Supplemental Figure S17.**
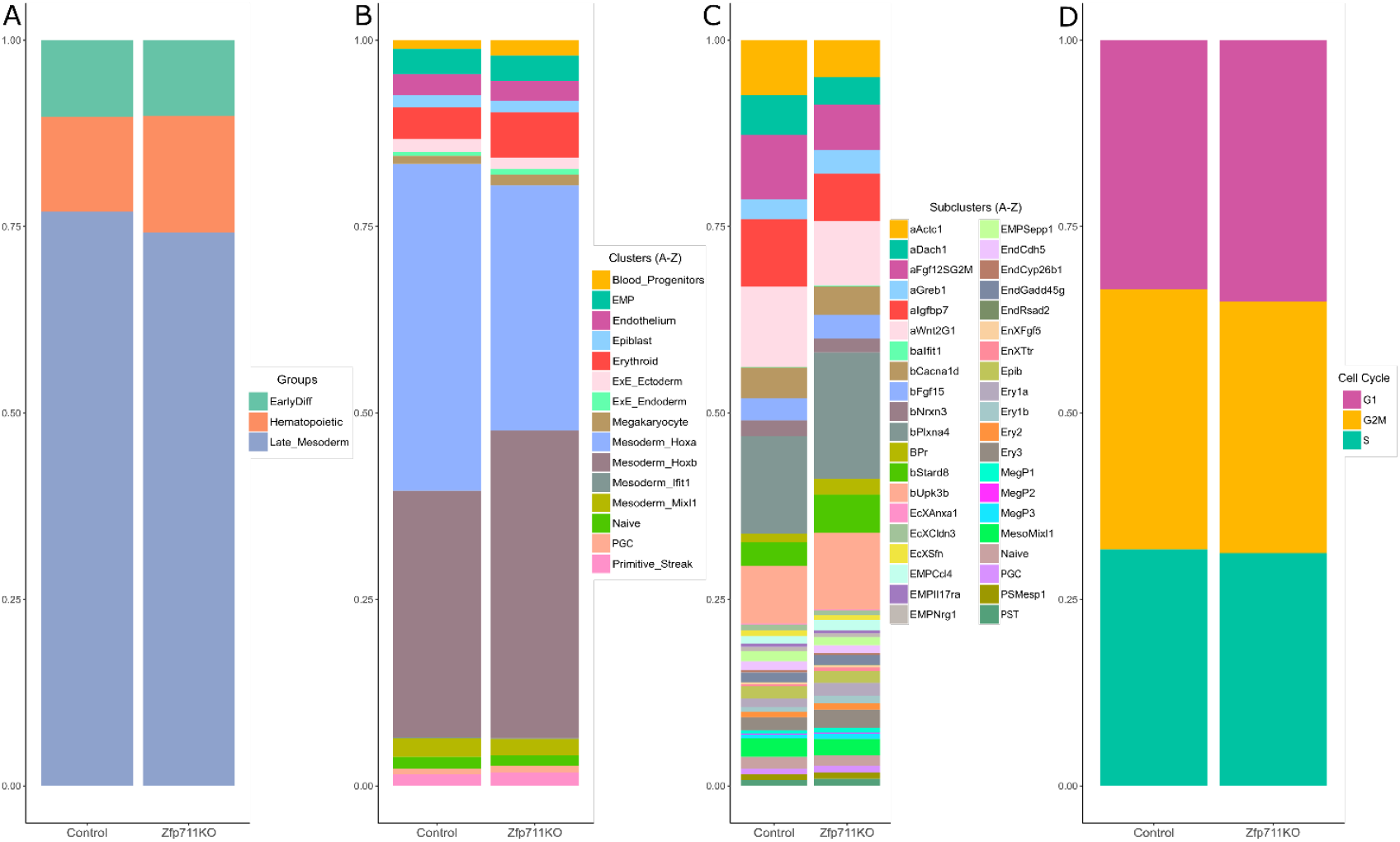
*Zfp711*-KO Differential Abundance Analysis results with Speckle R Package. **A-D**. Fraction of the annotated cell types in order of groups (A), clusters (B), sub-clusters (C), and cell cycle (D).

**Supplemental Figure S18.**
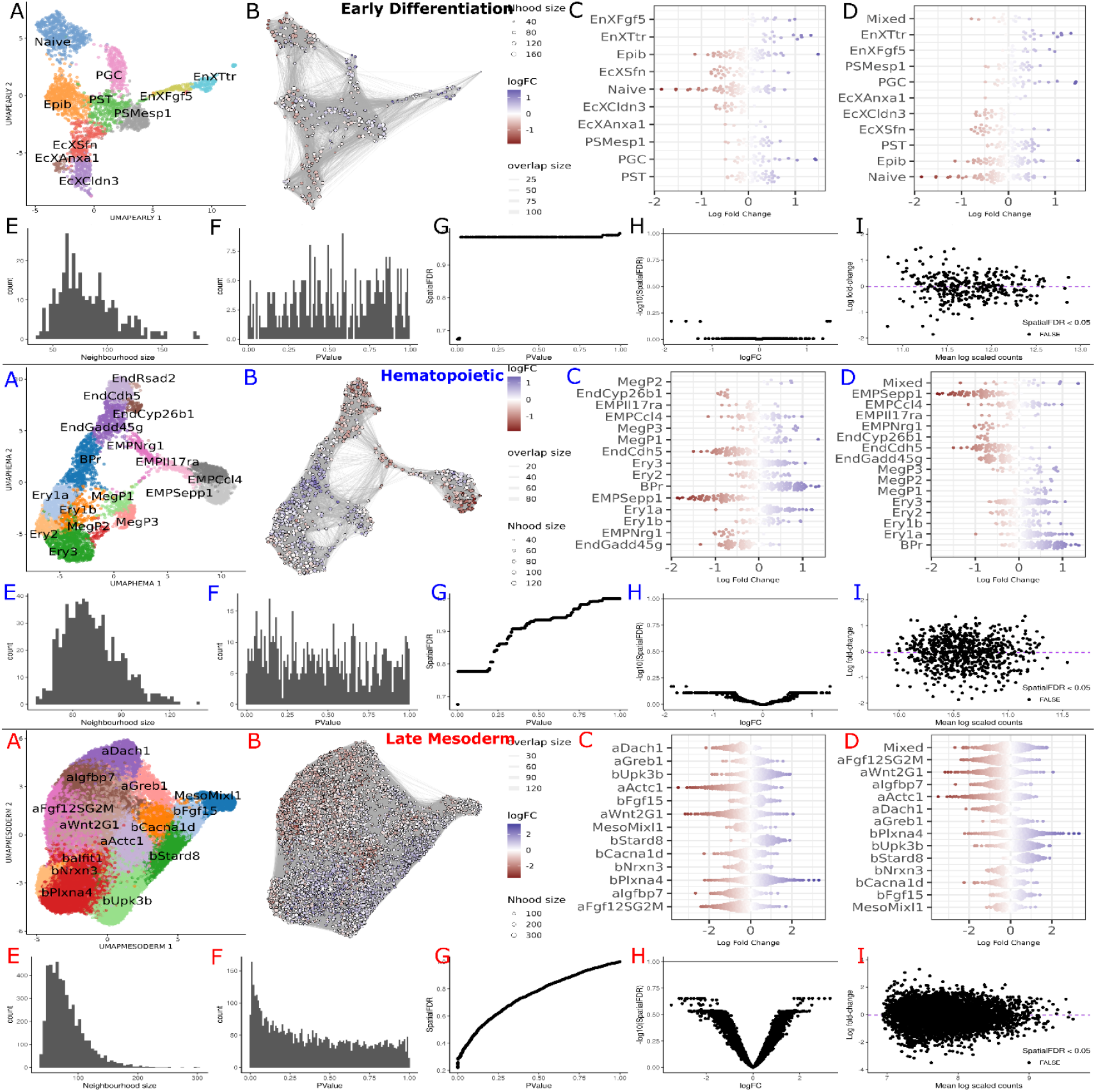
*Zfp711*-KO miloR Differential Abundance Analysis: detailed results in different groups on Early Differentiation (Black Label A-I), Hematopoietic (Blue Label A-I), and Late Mesoderm (Red Label A-I). **A**. UMAP with cell type annotation. **B**. miloR neighborhoods visualization on UMAP coordinates. **C**. Neighborhoods categorization into sub-clusters. **D**. Neighborhoods categorization to sub-clusters and mixed (50% threshold). **E**. Distribution of the size of the neighborhood. **F**. P-value histogram of the neighborhoods. **G**. P-value ~ Spatial FDR relationship for neighborhoods. **H**. Fold change and SpatilFDR relationship. **I**. Mean log scaled counts and log fold-change relation of the neighborhoods. The expectation is that the neighborhoods will be located around log fold-change 0. This figure is Supplemental to Figure 7K-M, and the dataset that outputs the analysis can be seen in Supplemental Table S3D-F.

**Supplemental Figure S19.**
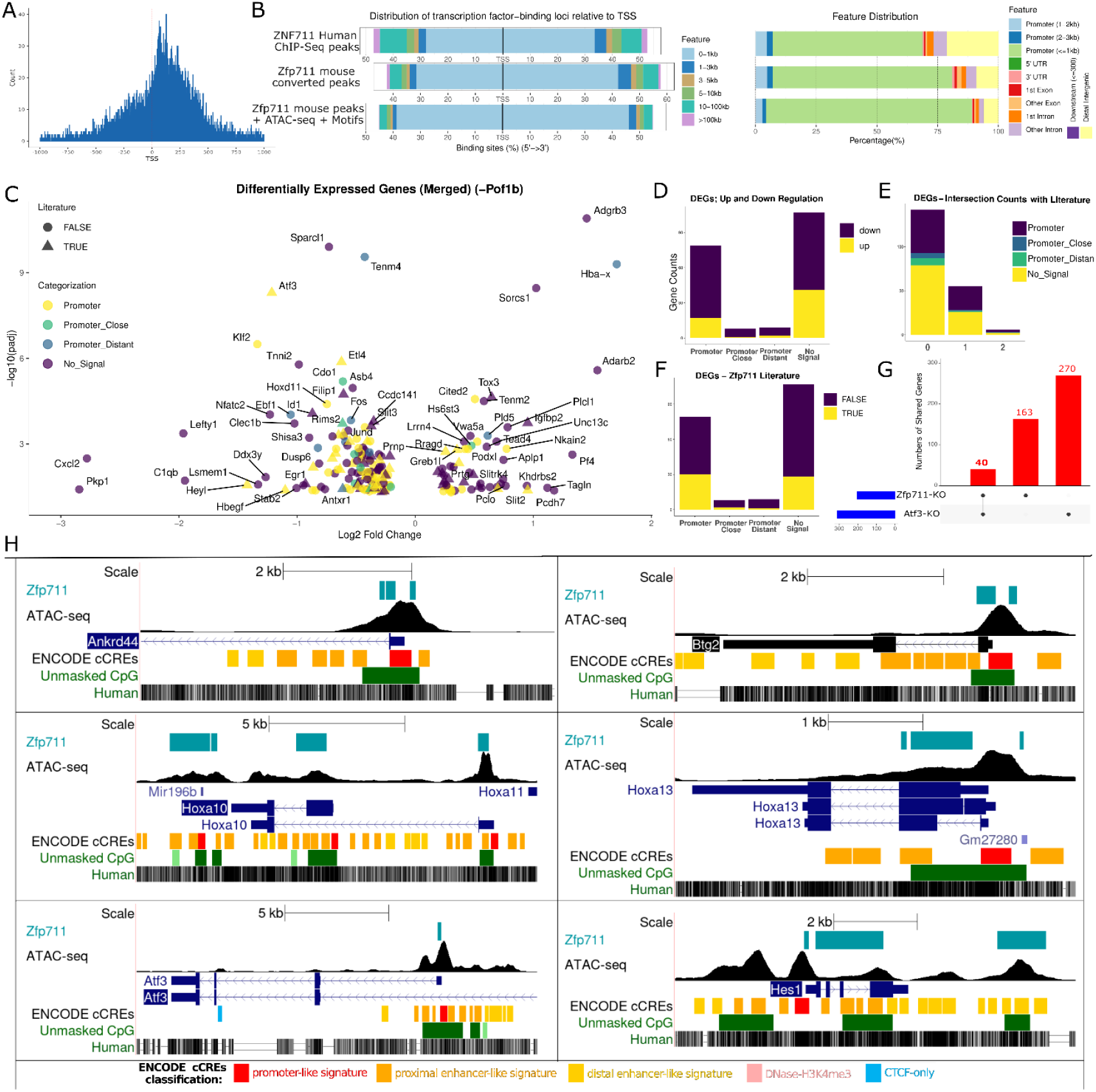
Results and categorization of the Differential Gene Expression Analysis of the *Zfp711*-KO cells. **A**. Zfp711 binding regions distribution histogram relative to the Transcription Start Site (TSS). **B**. Location and distance to the TSS of the Zfp711/ZNF711 ChIP-seq peak regions in three stages of the analysis from human to mouse. **C**. Volcano plot of the total significant DEGs. **D-G**, Distribution of the DEGs, Zfp711-binding categorization, shared DEGs with *Atf3*-KO cells, and DEGs (as reported in the literature). **H**. Some examples of Zfp711 binding sites with ATAC-seq peaks in three example DEGs. This figure is Supplemental to Figure 8D-F. Supporting data can be seen in Supplemental Table S4L-s, Supplemental Table S6B, and Supplemental Table S7F-J.

**Supplemental Figure S20.**
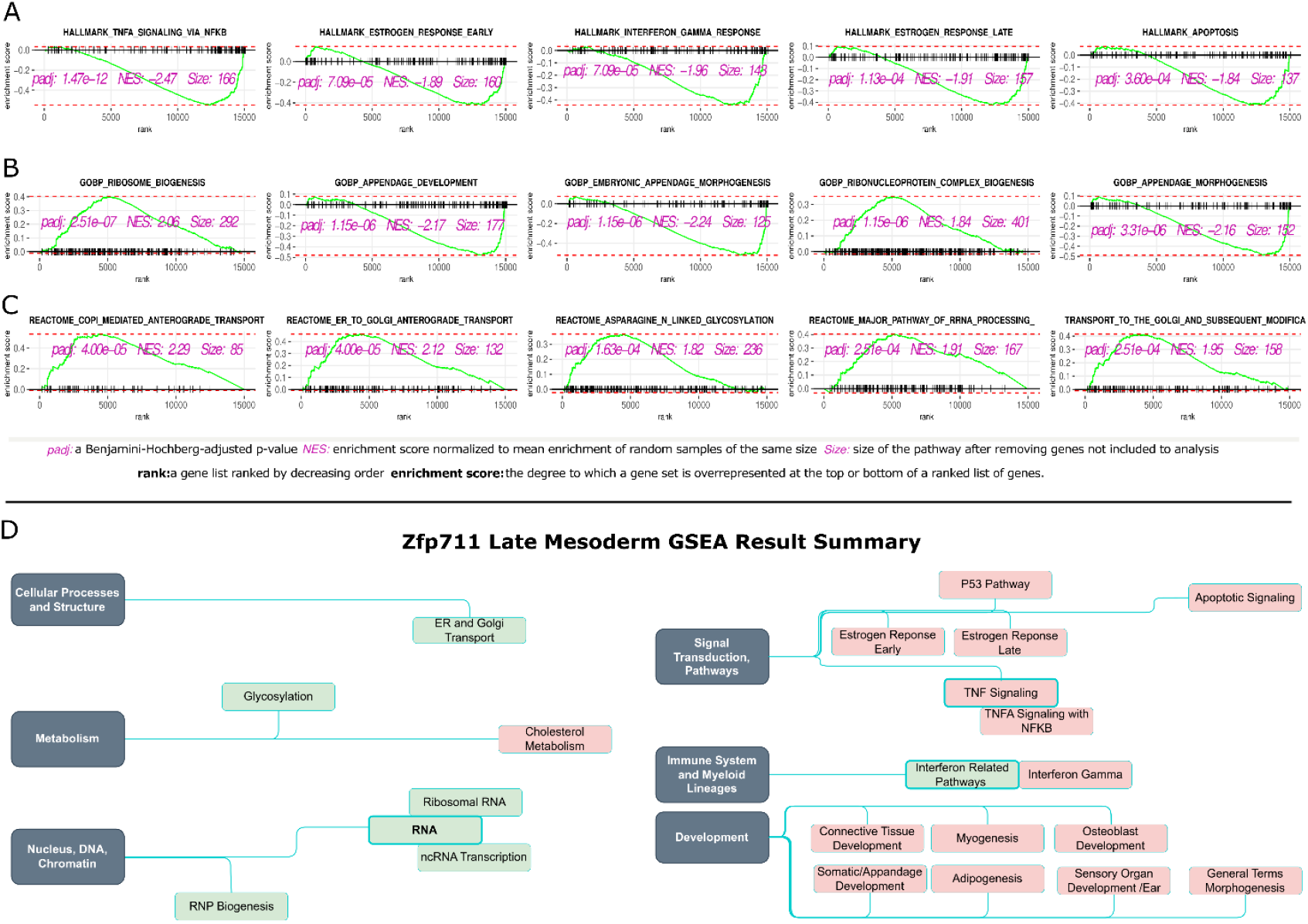
GSEA results of the *Zfp711*-KO cells as Late Mesoderm. **Panels A**-**C** show enrichment plots of the top-5 results from Late Mesoderm GSEA results. **Panel D** shows the manually grouped GSEA results of the *Zfp711*-KO cells on Late Mesoderm. This figure is Supplemental to Figure 9, and supporting data can be seen in Supplemental Table S5G-I.

**Supplemental Figure S21.**
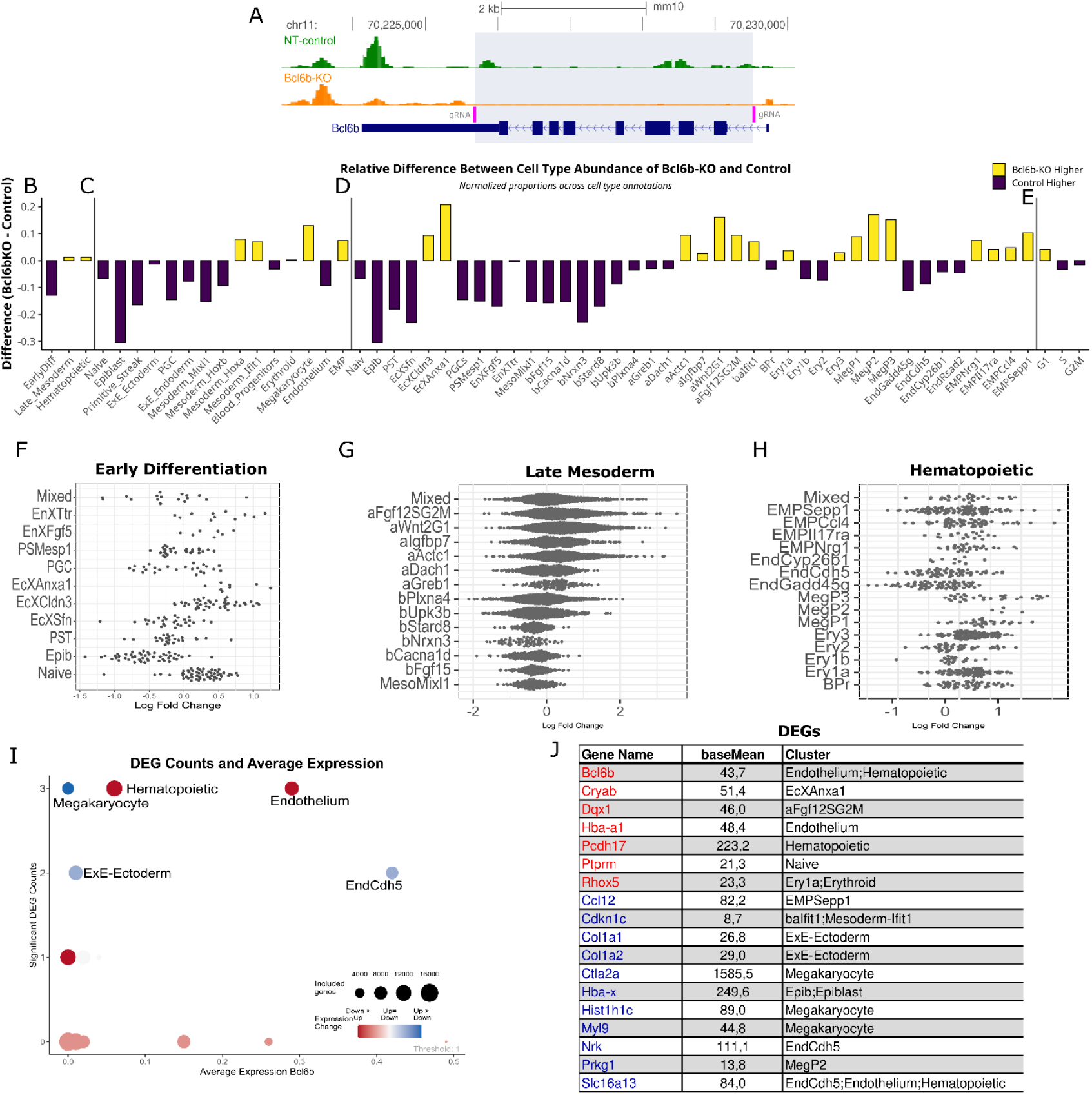
Results with *Bcl6b*-KO cells. **Panel A** shows the knockout approach for *Bcl6b*; the position of the gRNAs is shown in pink. The peaks represent the scRNA-seq result for the control mESCs in green and the KO in orange. The gene structure of *Bcl6b* is shown in dark blue. **Panels B-D** compare the number of NT-Control versus the Bcl6b-KO cells in groups, clusters, and sub-clusters. **Panel E** compares the number of cells in the respective phases of the cell cycle. **Panels F-H** show the fold-change in the differential abundance analysis in the different groups while creating small neighborhoods (small groups of cells) shown as dots (each dot represents mainly 50-100 cells). Spatial FDR < 0.1, higher values are shown in grey. The text on the y-axis indicates sub-clusters, and the log fold change is shown on the x-axis. “Mixed” on the y-axis indicates a neighborhood that does not belong to any sub-cluster (using a 50% threshold). **Panel I** shows the average expression level and significant DEGs (padj < 0.05) in groups, clusters, and sub-clusters. The size of the circles indicates the number of genes included in the analysis after removing the low-expressed genes in the group, cluster, or sub-cluster. The color of the circle indicates up- or downregulation (*blue* and *red*, respectively). **Panel J** shows all the DEGs, their expression level (*baseMean*), and to which cluster they belong.

**Supplemental Figure S22.**
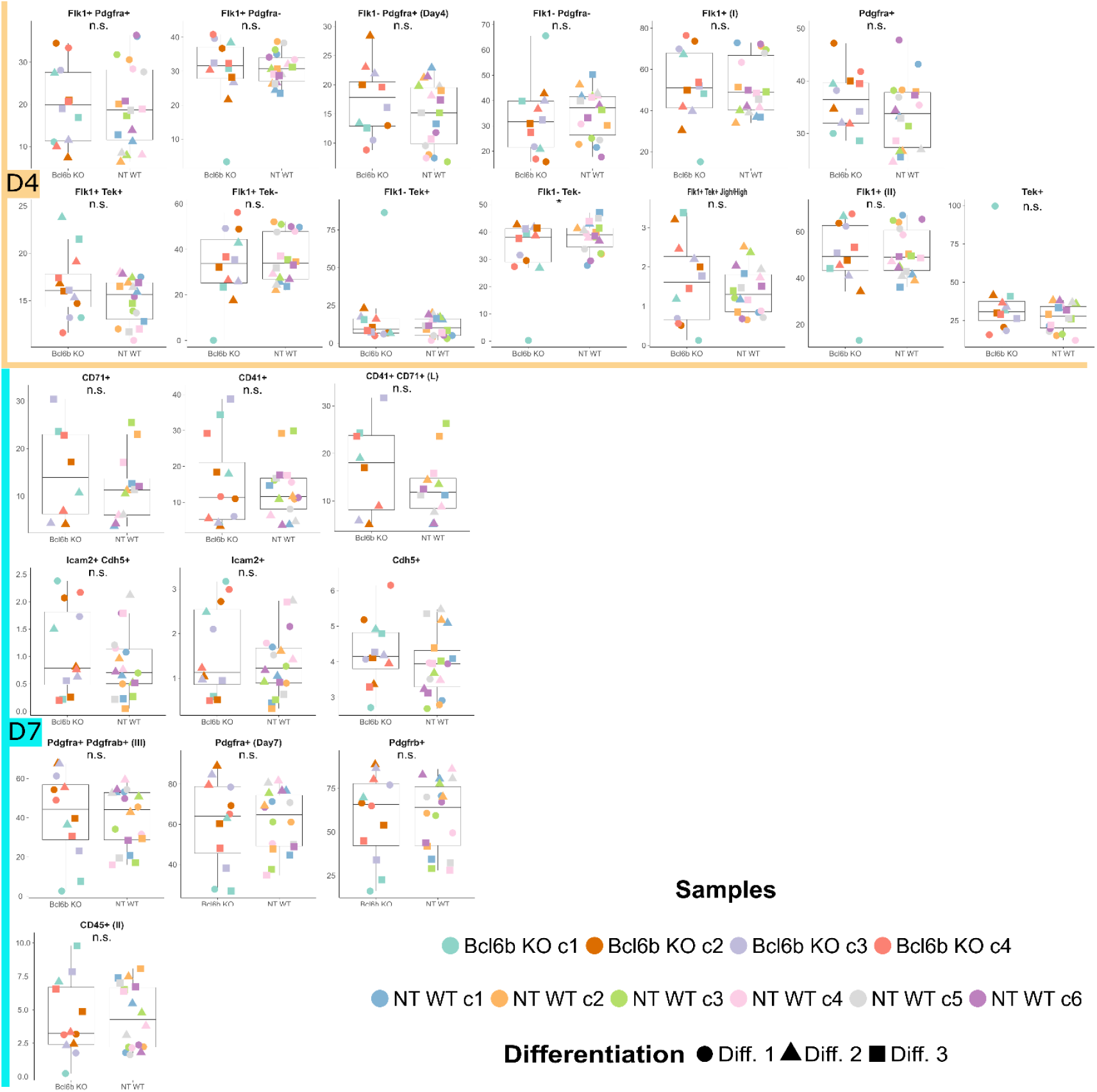
Flow Cytometric Analysis results of the *Bcl6b*-KO versus NT-Control cells (D4 (orange) and D7 (turquoise)). Details of the dataset and statistical analysis can be accessed in Supplemental Table S1E-F. (n.s.) not significant: P > 0.05, (*) P < 0.05, (**) P < 0.01, (***) P < 0.001.

**Supplemental Figure S23.**
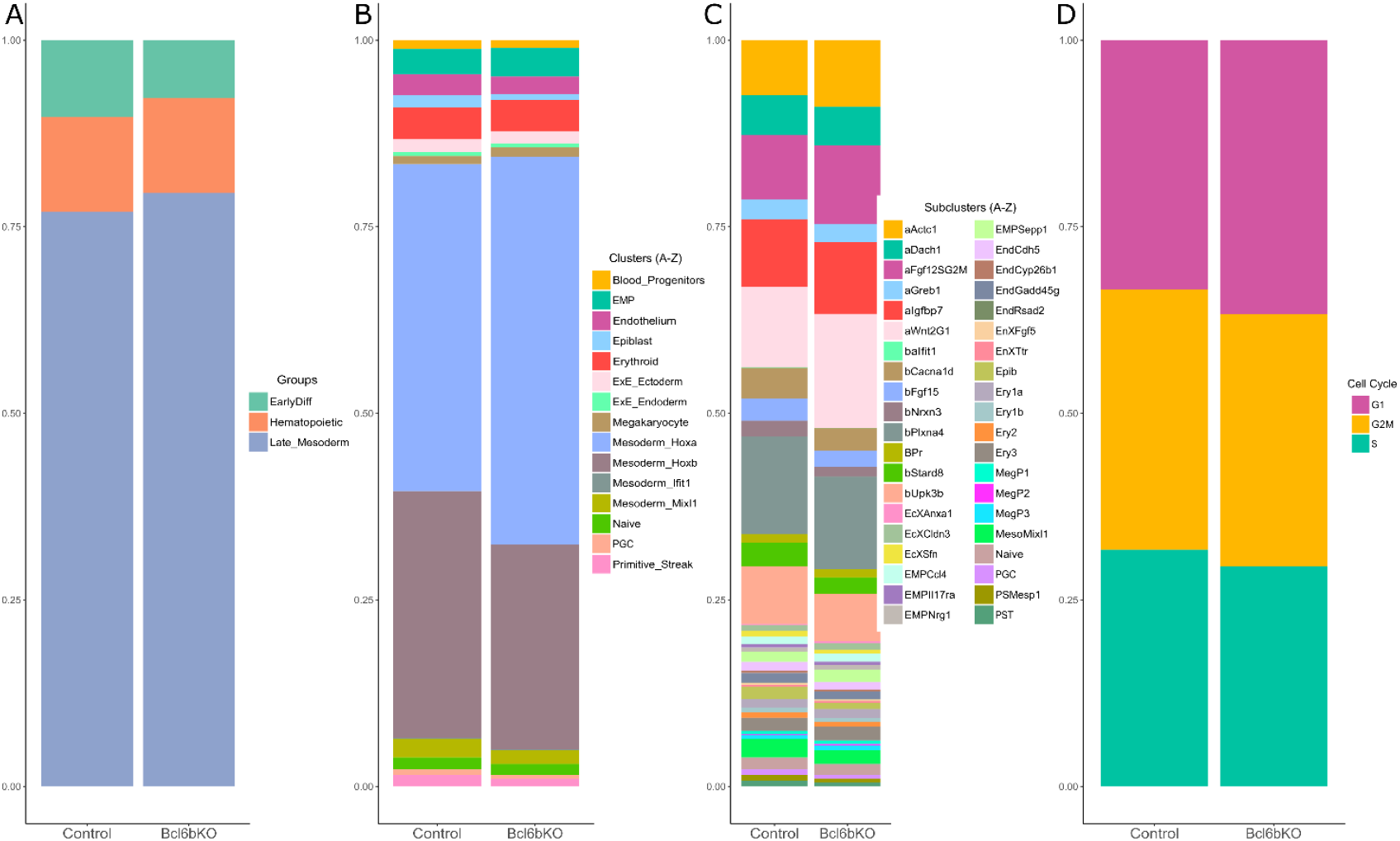
*Bcl6b*-KO cells: Differential Abundance Analysis results with Speckle R Package. **Panels A**-**D**. Fraction of the annotated cell types in order of groups (**A**), clusters (**B**), sub-clusters (**C**), and cell cycle (**D**). This figure is Supplemental to Figure 10B-E. Outputs of the Speckle differential abundance analysis can be seen in Supplemental Table S2Q-X.

**Supplemental Figure S24.**
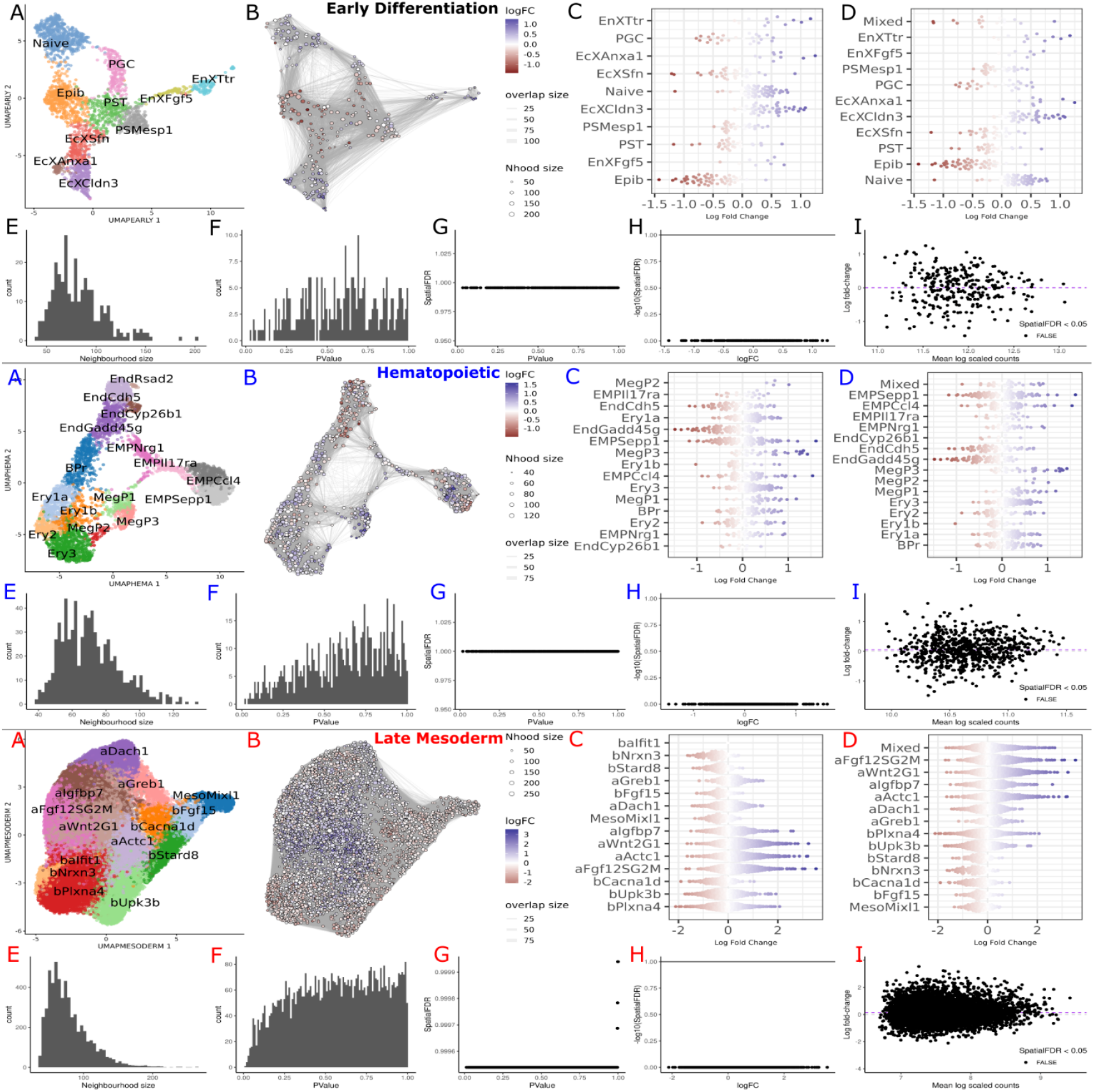
*Bcl6b*-KO cells: miloR Differential Abundance Analysis, detailed results in different groups, i.e. Early Differentiation (Black Label A-I), Hematopoietic (Blue Label A-I), and Late Mesoderm (Red Label A-I). **A**. UMAP with cell type annotation. **B**. miloR neighborhoods visualization on UMAP coordinates. **C**. Neighborhoods categorization into sub-clusters. **D**. Neighborhoods categorization into sub-clusters and mixed (50% threshold). **E**. Distribution of the size of the neighborhood. **F**. P-value histogram of the neighborhoods. **G**. P-value ~ Spatial FDR relationship for neighborhoods. **H**. Fold-change and Spatial FDR relationship. **I**. Mean log-scaled counts and log fold-change relation of the neighborhoods. The expectation is that the neighborhoods will be located around log fold-change 0. This figure is Supplemental to Figure 10F-H, and the dataset that outputs the analysis can be seen in Supplemental Table S3G-I.

**Supplemental Figure S25.**
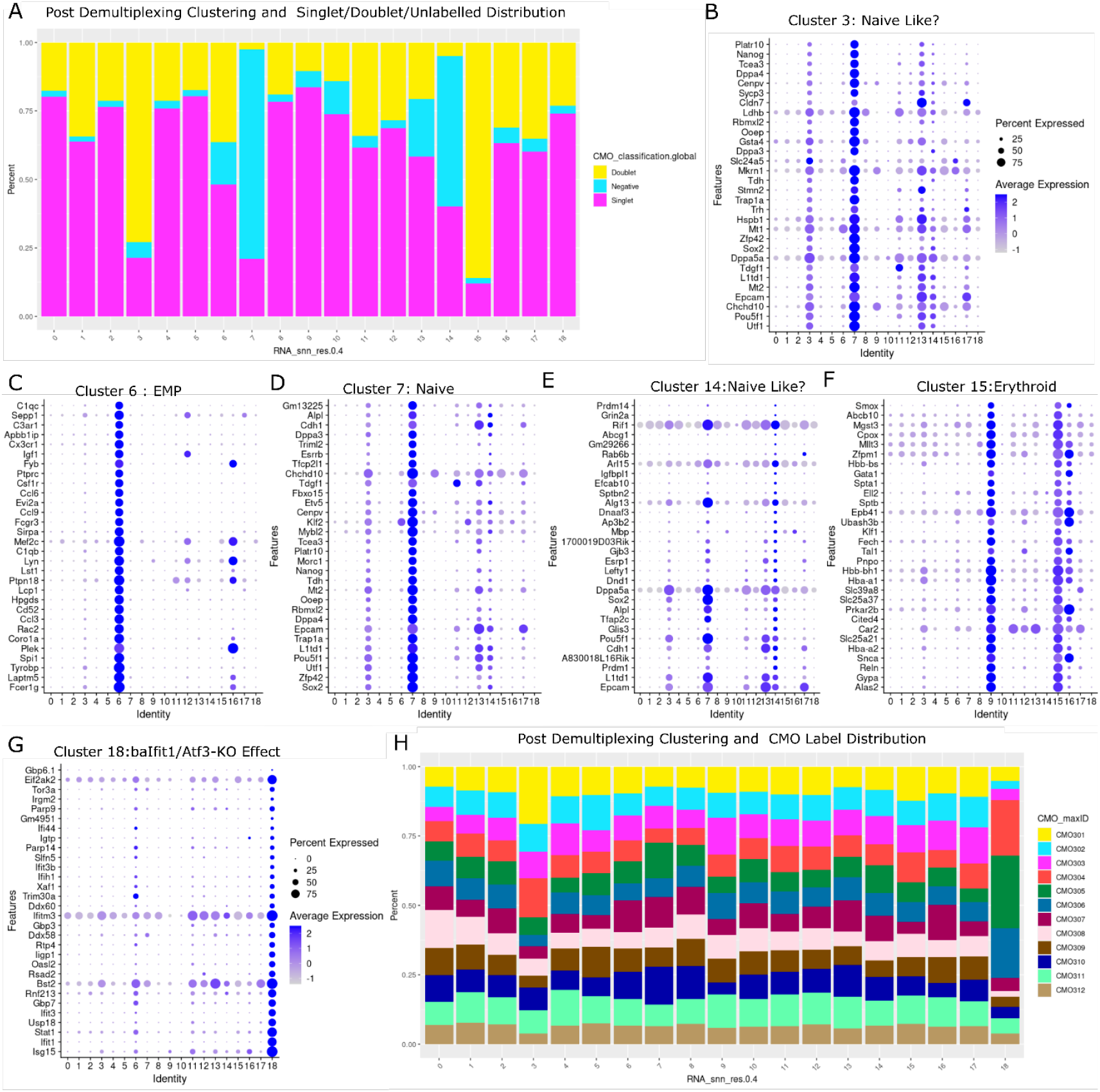
Demultiplexing results of CMO labels and their distribution across cell types. **Panel A**. Distribution of demultiplexed CMO labels into doublets, singlets, and negatives (unlabeled) following quality control steps. The X-axis represents clusters (0-18), and the Y-axis shows the percentage distribution of doublets, singlets, and negatives. **Panels B-C**. A plot of the highly expressed genes in the clusters identified in Panel A. **Panel G:** Displays the highly expressed genes in Cluster 18. **Panel H** Shows the distribution of CMO labels across clusters.

**Supplemental Figure S26.**
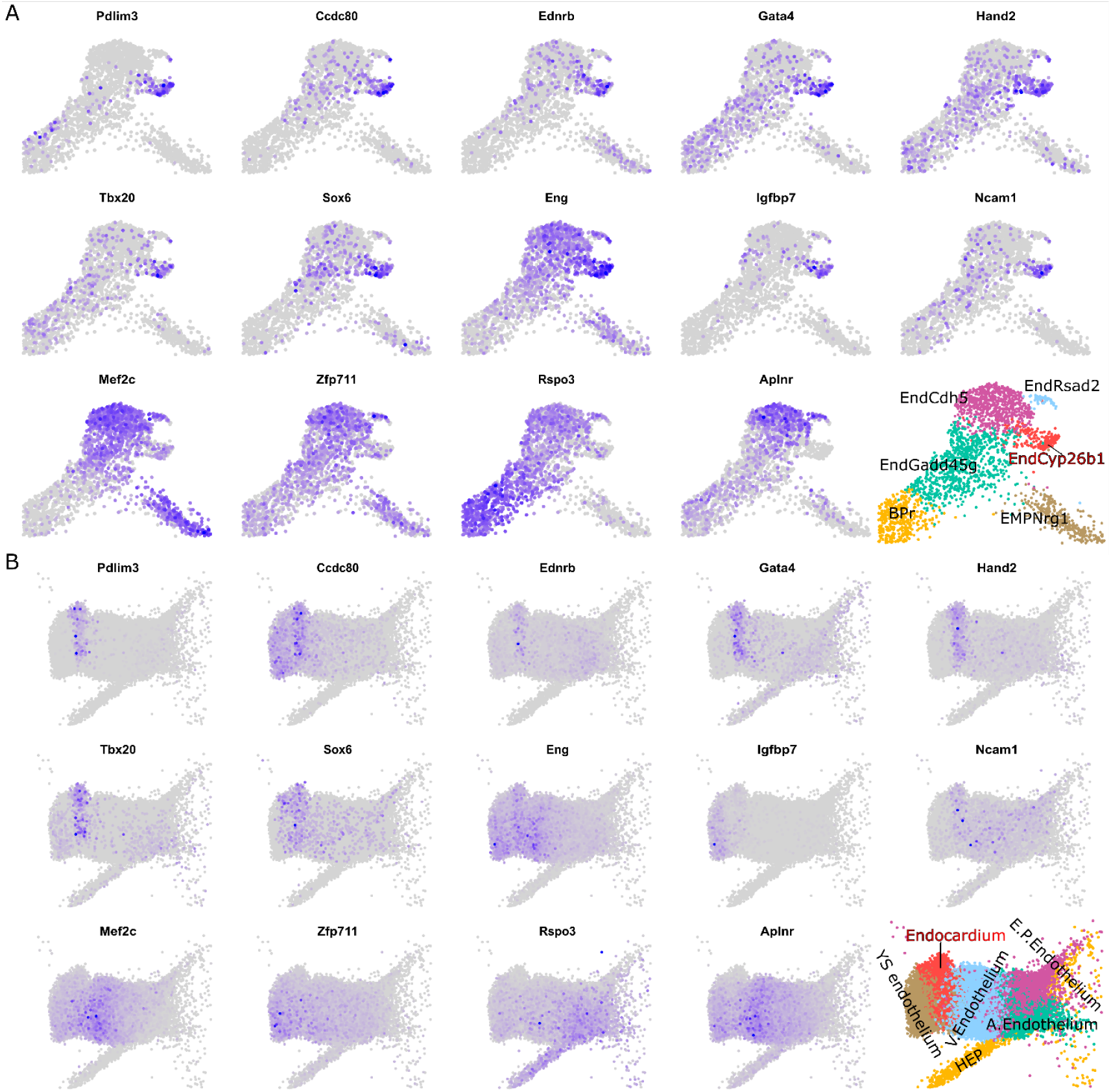
Mapping of marker genes in the EndCyp26b1 sub-cluster. **Panel A**. Mapping the 10 up- and 4 down-regulated genes on the zoomed-in Hematopoietic Group UMAP, focusing on the endothelial sub-clusters, and highlighting the EndCyp26b1 sub-cluster. **Panel B**. Mapping similar to A, using data from the Extended Mouse Atlas (Imaz-Rosshandler et al. 2023), demonstrating the spatial distribution of the marker genes within the endothelial context.

